# Ridge Redundancy Analysis for High-Dimensional Omics Data

**DOI:** 10.1101/2025.04.16.649138

**Authors:** Hayato Yoshioka, Julie Aubert, Hiroyoshi Iwata, Tristan Mary-Huard

## Abstract

**Motivation:** Redundancy Analysis (RDA) is a popular reduced rank regression approach for modeling the relationships between two sets of variables. In omics studies, RDA provides a flexible framework for investigating associations between high-dimensional molecular data. However, omics datasets often exhibit strong multicollinearity and suffer from the “large *p*, small *n*” problem, where the number of predictor variables *p* exceeds the number of samples *n*. Ridge RDA addresses multicollinearity by introducing a ridge penalty in addition to the rank restriction. Despite these advantages, its application to high-dimensional omics data remains challenging due to i) difficulties in manipulating the large coefficient matrix of dimensions *p* × *q*, where *q* represents the number of response variables, and ii) the need to jointly select the rank and the regularization parameter that both influence the performance of the method.

**Results:** We propose an efficient computational framework for ridge RDA that overcomes these challenges by leveraging the Singular Value Decomposition of the predictor matrix *X*. This approach eliminates the need for direct covariance matrix inversion, improving computational efficiency. Furthermore, we introduce a novel strategy that reduces the memory burden associated with the storage of the coefficient matrix from *pq* to (*p* + *q*)*r*, with *r* the chosen reduced-rank dimensionality. Our method enables an efficient grid search to jointly select the ridge penalty *λ* and *r* via cross-validation. The proposed framework is implemented in the R package **rrda**, providing a practical and scalable solution for the analysis of high-dimensional omics data.

**Availability:** The R package `rrda` is available on CRAN: https://CRAN.R-project.org/package=rrda/rrda. Scripts and data used for the analysis are available on GitHub: https://github.com/Yoska393/rrda/rrda.

## 1. Introduction

Redundancy Analysis (RDA; Van den Wollenberg 1977) is a widely used multivariate statistical method for investigating relationships between two sets of variables (Reinsel and Velu, 1998). Also referred to as reduced-rank regression (Anderson, 1951) and principal components of instrumental variables, RDA is a flexible technique for modeling multiple response variables as functions of multiple predictor variables. Unlike Canonical Correlation Analysis (Hotelling, 1936), which maximizes the correlation between two variable sets, RDA aims to optimize the predictability of one set from the other.

RDA is commonly applied in ecological studies, where the response matrix *Y* of dimensions *n* × *q* typically represents community composition, while the predictor matrix *X* of dimensions *n* × *p* contains environmental variables explaining the variation in *Y*. The method is based on the multivariate linear model *Y* = *XB* + *E*, where *B* is the coefficient matrix subject to the rank constraint rank(*B*) ≤ *r*, ensuring dimensionality reduction.

Rank restriction is beneficial as it efficiently captures and exploits redundant information, and facilitates the identification of dominant patterns in the data (Capblancq and Forester, 2021).

In recent years, RDA has been increasingly applied in various omics studies. The field of multi-omics has emerged as a powerful approach for deciphering complex biological pathways (Buescher and Driggers, 2016). Notable applications include gene expression analysis (Wen et al., 2023), DNA methylation studies (Ruiz-Arenas and González, 2017), and metabolomic research (Hilafu et al., 2020). However, omics data are typically high-dimensional and characterized by highly correlated variables, leading to challenges related to multicollinearity.

To mitigate multicollinearity, regularized redundancy analysis (Takane and Hwang, 2007), also referred to as reduced-rank ridge regression (Mukherjee and Zhu, 2011), has been considered in omics research. By introducing a regularization term, ridge regression reduces the impact of collinearity by shrinking the coefficients of correlated predictors, thus preventing overfitting and enhancing generalization performance. This regularization improves stability and generalization to new datasets, making it a valuable tool for analyzing complex biological systems (Wen et al., 2023).

In high-dimensional regression models, where the objective is to predict a single response variable from a large set of predictors, a primary challenge is the “large *p*, small *n*” problem, in which the number of predictor variables (*p*) exceeds the number of samples (*n*). This setting is particularly prevalent in genomics and other omics disciplines (Crossa et al., 2017; Zhao et al., 2019; Li et al., 2024). The *n* ≪ *p* problem has been widely addressed in the univariate response setting (Lawson and Hanson, 1995; Turlach, 2006), and efficient implementations are readily available to users (Friedman et al., 2023). However, in the multivariate setting, ridge RDA raises new challenges, as both the number of predictor variables (*p*) and the number of response variables (*q*) can be large (e.g., *p, q* > 10^5^). This results in a coefficient matrix *B* ∈ ℝ^*p* × *q*^ whose size grows multiplicatively with *p* and *q*, leading to significant computational and storage challenges (Yuan et al., 2007; Kong et al., 2020; Ke et al., 2022). Despite these limitations, efficient statistical frameworks that explicitly address both computational burden and memory requirements for ridge RDA remain underdeveloped, particularly in high-dimensional omics applications.

To address these challenges, we propose a novel approach for computing the regularization path and optimizing data storage for ridge RDA. We introduce a new decomposition of the low rank coefficient matrix *B* that reduces its storage in memory from *pq* to (*p* + *q*)*r*, where *r* denotes the rank of matrix *B*, making the approach amenable to high-dimensional settings. These computational enhancements open the way to efficient parameter tuning, where the ridge regularization parameter *λ* and the rank (*r*) are jointly selected through a cross-validation (CV) process. Our method is implemented in the R package **rrda**, publicly available on CRAN.

This article introduces an efficient ridge RDA framework designed for high-dimensional omics data. We begin by providing a theoretical foundation that demonstrates how our approach improves computational efficiency in ridge RDA. Next, we conduct simulation studies to compare the performance of our method with classical approaches across a range of dataset sizes. Finally, the method is illustrated through several applications to high-dimensional omics datasets of varying sizes and types, including DNA copy number, gene expression, metabolome, microbiome, and methylation-expression data. These applications demonstrate the ability of the approach to handle high-dimensional data.

## 2. Materials and Methods

### 2.1. Ridge redundancy analysis

Let *Y* be an *n* × *q* matrix of response variables, and let *X* be an *n* × *p* matrix of predictor variables. We assume that the columns of matrices *Y* and *X* are centered to have means of 0. The relation between *X* and *Y* is given by

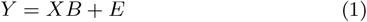

where *B* is the matrix of regression coefficients, and *E* is an error matrix. The ridge RDA aims to estimate *B* under both rank and ridge restrictions. The corresponding optimization problem is

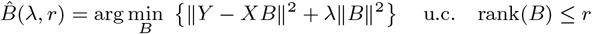

where ‖ *A* ‖^2^ =tr(*A*’*A*) is the squared Frobenius norm and *λ* is the ridge regularization parameter. As shown in Takane and Hwang (2007), the previous optimization problem can be reformulated as

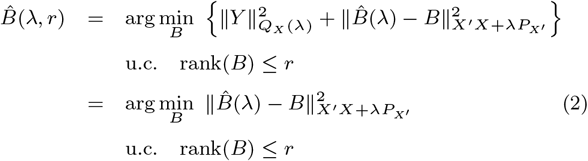

Where 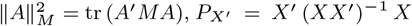, and 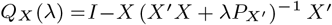. Matrix 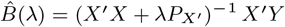 represents the ridge estimator of *B* without rank restriction.

The solution of (2) can be obtained by considering the Generalized Singular Value Decomposition (GSVD, Cailliez and Pages 1976; Greenacre 1984) of 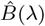 with respect to row and column metric matrices 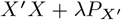 and *I*. See Appendix B for the definition of GSVD. This decomposition yields

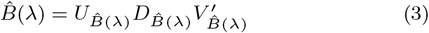

where 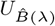 satisfies 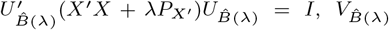 satisfies 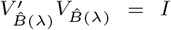, and 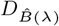 is a diagonal matrix. The ridge-penalized-*r*-rank estimate 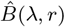 is then obtained by retaining only the GSVD components corresponding to the first *r* dominant singular values:

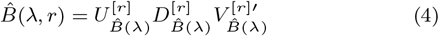

where 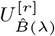 is a *p* × *r* matrix, 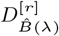 is an *r* × *r* diagonal matrix, and 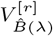 is a *q* × *r* matrix.

Computing the unconstrained ridge estimator requires the inversion of 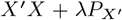, which is a large matrix of size *p* × *p*. The estimate 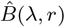 is also a large matrix (of dimensions *p* × *q*), and presents both computational and storage difficulties. Lastly, model parameter tuning requires repeating the inference procedure for each combination of the ridge (*λ*) and rank (*r*) constraints.

### 2.2. Adaptation to the high-dimensional setting

We now focus on ridge RDA in high-dimensional settings where *p, q* ≫ *n*. Similar to what is classically performed in the ridge regression setting (see Hastie et al. 2001, Section 18.3.5), we first perform a Singular Value Decomposition (SVD) of *X* as

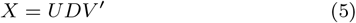

The different quantities appearing in Equation (2) can then be rewritten as follows:

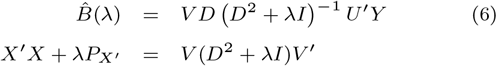

where we assume *λ >* 0. Note that this first SVD is performed only once, as it does not depend on parameters *λ* and *r*.

A second SVD is required for the computation of the GSVD of matrix 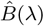. Indeed, one has

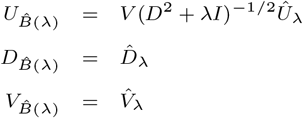

where 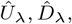 and 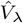 are the SVD components of matrix *M*_*λ*_ = (*D*^2^ + *λI*)^*−*1*/*2^*DU*^*’*^*Y*. As this matrix depends on *λ*, this second SVD must be computed for each specific value of *λ*. However, the computation of the matrix *M*_*λ*_ itself for different values of *λ* can be factorized as *M*_*λ*_ is the product of a diagonal matrix and a constant matrix *U*^*’*^*Y* that does not depend on *λ*.

The final expression of the 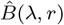 is then

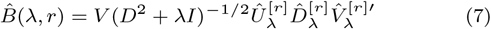

To summarize, the proposed formulation i) leverages the SVD of *X* to avoid the direct inversion of the 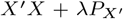 matrix, which is of size *p x p*, thereby significantly reducing computational cost, ii) reduces the computation of the GSVD that follows to the computation and storage of diagonal matrices (*D*^2^ + *λI*)^*−*1*/*2^ combined with the SVD of a *n x q* matrix *M*_*λ*_, offering efficiency gains over handling the *p x q* matrix *B*. See Appendices B and G for details.

### 2.3 Effcient storage of the high-dimensional coefficient matrix

In omics studies, where both *p* and *q* may be large, the direct storage of the ridge RDA coefficient *p x q* matrix may become challenging or even infeasible. In the Methylation application of Section 3.2.3, one has *p* = 392, 277 and *q* = 67, 528, leading to a coefficient matrix with *>* 10^10^ elements that would require a memory storage of ~ 100 GB. To address this issue, we propose an efficient storage method. From (4), define

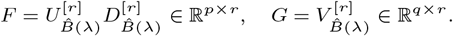

Denoting by *F*_*i*_ and *G*_*i*_ the *i*-th column of *F* and *G*, respectively, one has

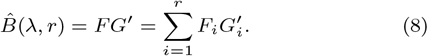

Moreover, the computation of any new prediction from the ridge RDA estimator can be obtained as

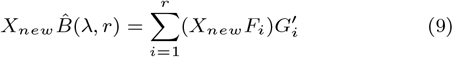

which does not require the computation of 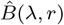. Consequently, the storage of the coefficient matrix can be replaced by the storage of the two matrices *F* = [*F*_1_, …, *F*_*r*_] and *G* = [*G*_1_, …, *G*_*r*_] of size *p* x *r* and *q x r*, respectively. This alternative representation drastically reduces the memory usage, moving, e.g., from 100 GB to 175 MB in the methylation application when considering *r* = 100.

The computation and storage of the 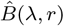 and of its usage for prediction are implemented in the R package `rrda` available on CRAN. The `rrda.fit` function computes the coeffcient matrix either in a component form, i.e., using the *F* and *G* matrices, or directly as a full *px q* matrix, for all combinations of *λ* and *r*. The `rrda.predict` function performs prediction using 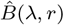 directly or following (9) whenever the component form is used.

An additional advantage of the low-rank representation is that it enables the extraction of a compact, interpretable submatrix for visualization. Define the symmetrically scaled factors as 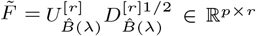 and 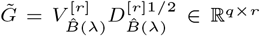, so that 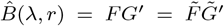. One can select the top *m*_*x*_ predictors and top *m*_*y*_ responses based on the row-wise *ℓ*_2_ norms of 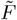 and 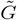, respectively. Denoting by 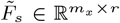 and 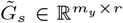 the submatrices formed by the selected rows, the reduced matrix

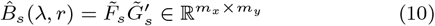

captures the dominant associations among the most relevant predictors and responses. Importantly, the procedure relies solely on 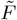 and 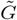, avoiding explicit construction of the full *p x q* coefficient matrix and thus preserving memory efficiency. This procedure is implemented in the `rrda.top` function of the `rrda` package. Further details are provided in Appendix F.

### 2.4. Parameter tuning

Based on the computational and storage shortcuts introduced in the previous section, we consider a *V*-fold CV method to jointly select *r* and *λ*. The criterion to optimize is

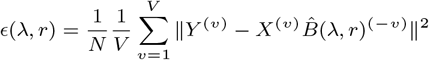

where (*v*) stands for the *v*^th^ subset of the data, 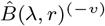 is the ridge RDA coefficient matrix estimated from all subsets except the *v*^th^ one, and *N* = *n*_*v*_ *x q* with *n*_*v*_ the number of samples in the *v*^th^ subset.

In practice, a sequence of values for *λ* is considered within a range [*λ*_min_, *λ*_max_] that must be provided by the user. Following the choice of the `glmnet` procedure (Friedman et al. 2010), the default value of the interval bounds can be set at

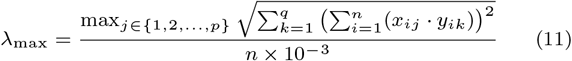

and *λ*_min_ = 10^−4^*λ*_max_, respectively

As the computational cost of (11) is *O*(*npq*), a random subsampling of the variables was implemented whenever max(*p, q*) *> m*, with a default value of *m* = 10^3^. In our experiments, this sampling approach significantly reduced the computation time without hampering accuracy (Figure 1S). Furthermore, we extended the `glmnet` “one-standard-error” rule (Friedman et al., 2010) to the selection of both the ridge penalty coefficient *λ* and the rank parameter *r*, selecting the largest *λ* and smallest *r* within one-standard-error of the minimum mean squared error (MSE).

**Fig. 1:**
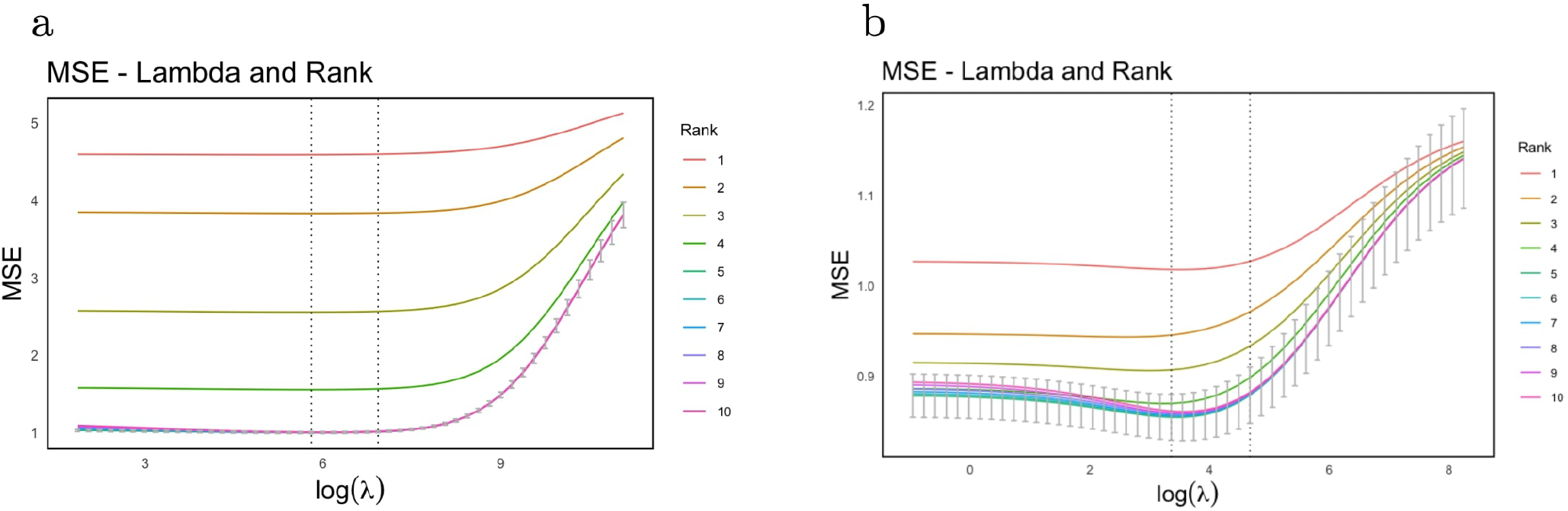
Visualization of the RRDA regularization path computed on two datasets simulated with model 1 (a) and model 2 (b), respectively. Each plot displays the 5-fold cross-validated mean squared error (CV-MSE) for all combinations of ridge penalty *λ* and rank. The left vertical dotted line indicates the ω value that minimizes the CV-MSE, while the right vertical dotted line corresponds to the *λ* selected by the one-standard-error rule. Error bars correspond to CV standard errors.

The optimal (*r,λ*) combination can be selected through CV using the `rrda.cv` function of the R package `rrda`, which also benefits from parallel computing (see Appendix G). In our experiments, we applied 5-fold CV via RRDA (hereafter `rrda.cv`) to jointly select *r* and *λ*, and via the classic non-ridge RDA (hereafter `c-rda.cv`, i.e., `rrda.cv` with *λ* = 0) to select *r*.

Alternatives to CV for model selection include information criteria. In what follows, we benchmarked the CV approach against five criteria that have been proposed in the context of reduced rank regression: Akaike Information Criterion (AIC; Akaike 1974), Bayesian Information Criterion (BIC; Schwarz 1978), Generalized Information Criterion (GIC; Fan and Tang 2013), Generalized Cross-Validation (GCV; Golub et al. 1979), and Bayesian Information Criterion with Penalty (BICP; An et al. 2008). Although GCV was originally introduced as an approximation to CV, here we regard it as a type of information criterion. The mathematical expressions of the different criteria are provided in Table 1.

**Table 1.** Evaluation criteria for tuning parameter selection. Here, *r*_*x*_ denotes the rank of the explanatory variable matrix *X, q* is the number of response variables, *p* the number of explanatory variables, *n* the sample size, and 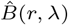 the estimator depending on rank *r* and penalty parameter *λ*.

| **Criterion** | **Formula** |
|---|---|
| AIC | $nq\log\left(\frac{\|Y-X\hat{B}(r,\lambda)\|^2}{nq}\right)+2\,r(r_x+q-r)$ |
| BIC | $nq\log\left(\frac{\|Y-X\hat{B}(r,\lambda)\|^2}{nq}\right)+\log(nq)\,r(r_x+q-r)$ |
| GIC | $nq\log\left(\frac{\|Y-X\hat{B}(r,\lambda)\|^2}{nq}\right)+\log(nq)\,\log(pq)\,r(r_x+q-r)$ |
| BICP | $nq\log\left(\frac{\|Y-X\hat{B}(r,\lambda)\|^2}{nq}\right)+2\,\log(pq)\,r(r_x+q-r)$ |
| GCV | $\frac{nq\|Y-X\hat{B}(r,\lambda)\|^2}{\left[nq-r(r_x+q-r)\right]^2}$ |

### 2.5. Competing methods

Two competing reduced-rank regression approaches were considered. The first one is the Adaptive Nuclear Norm procedure (ANN, Chen et al., 2013) that implements rank reduction by adaptively thresholding the singular values of *B* via a tuning parameter *ν*.

Each value of *ν* uniquely determines an estimated rank, yielding a set of candidate ranks *ℛ ⊆ {*1,…, min*{n, p, q}}*. The optimal rank is then selected by minimizing an information criterion over *ℛ*. Importantly, ANN does not incorporate a ridge penalty, and addresses only rank selection. The ANN procedure is implemented in the `rrr` function of the R package `rrpack` (version 0.1-13; Chen and Wang 2022). The choice of the regularization parameter can be performed using any of the information criteria listed in Table 1, or alternatively using the Stability Approach to Regularization Selection for Reduced-Rank Regression (`StARS`, Wen et al., 2023); see Appendix E for details. Hereafter, these criteria-based ANN methods are denoted as `ANN.AIC, ANN.BIC`, etc.

Secondly, the sparse RDA (`sRDA`, Csala et al., 2017) was considered. The `sRDA` approach implements Redundancy Analysis with an Elastic Net penalty to obtain parsimonious solutions in high-dimensional omics datasets. It can perform ridge RDA when the Elastic Net penalty is tuned toward its *ℓ*_2_ component. `sRDA` defines the number of latent variables within a partial least squares regression framework, and therefore differs from `rrda` in both its objective function and rank reduction strategy. In this study it serves as a recent RDA method for comparison in terms of predictive accuracy and computational performance, as presented in later sections. See Appendix D for details on `sRDA`.

In summary, the methods compared in this study differ in their objective functions, regularization strategies, and tuning criteria; a structured overview is provided in Table 2.

**Table 2.** Summary of the compared methods, including their objective functions and tuning strategies. `rrda` and `c-rda` operate within a reduced-rank regression framework with and without ridge regularization, respectively. ANN implements rank reduction via adaptive nuclear norm thresholding. `sRDA` is based on a latent-variable redundancy analysis framework with elastic-net penalization.

| **Method** | **Objective function** | **Hyperparameters** | **Tuning strategy** |
|---|---|---|---|
| rrda | $\|Y - XB\|^2 + \lambda\|B\|^2,\ \mathrm{rank}(B) \le r$ | λ, *r* | CV / AIC / BIC / GIC / GCV / BICP |
| c-rda | $\|Y - XB\|^2,\ \mathrm{rank}(B) \le r$ | *r* | CV / AIC / BIC / GIC / GCV / BICP |
| ANN | $\frac{1}{2}\|Y - XB\|^2 + \nu \sum_i w_i d_i(XB),\ w_i = d_i^{-\gamma}(\hat{Y}_{\mathrm{LS}})$ | ν | AIC / BIC / GIC / GCV / BICP / StARS |
| sRDA | $\|Y\beta_h - X\alpha_h\|^2 + \lambda_2\|\alpha_h\|^2 + \lambda_1\|\alpha_h\|_1,\ h = 1, \ldots, r$ | λ<sub>1</sub>, λ<sub>2</sub>, *r* | CV |

### 2.6. Simulation data

In this subsection, we consider two data generation models to illustrate the performance of Ridge RDA.

Model 1: Latent space model

In Model 1, both the predictor and response variables are generated from a shared latent space. Let *H ∈* ℝ^*nxk*^ be a latent variable matrix, where *n* is the number of observations and *k* is the dimension of the latent space. The entries of *H* are independently drawn from a standard normal distribution:

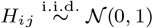

Let 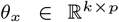 and 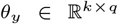 be coefficient matrices, with their entries also independently drawn from standard normal distributions:

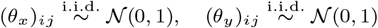

The observed predictor matrix *X ∈* ℝ^*nxp*^ and response matrix *Y ∈* R^*nxq*^ are then generated as:

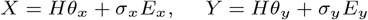

where

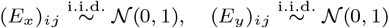

The scaling parameters *σ*_*x*_ and *σ*_*y*_ are chosen afterwards in order to ensure chosen signal-to-noise ratio *s*_*x*_ and *s*_*y*_, i.e.,

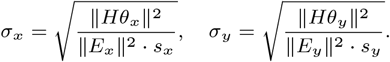

We tested the signal-to-noise (s2n) values *s*_*x*_ = *s*_*y*_ = 0.1, 1, 10; the results presented in the following sections correspond to *s*_*x*_ = *s*_*y*_ = 1.

This first setting does not explicitly rely on a true coefficient matrix *B* to which the ridge RDA estimator should be compared.

Model 2: Matrix factorization model

Let *X ∈*ℝ^*nxp*^ be the predictor matrix, with entries sampled from a standard normal distribution:

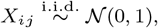

and *A ∈* ℝ^*pxk*^ and *C ∈* ℝ^*kxq*^ be two coefficient matrices, with *k* the true latent dimension. Both *A* and *C* are drawn from standard normal distributions:

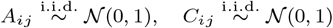

The columns of *A* and *C* are then normalized to form *A*^*\**^ and *C*^*\**^, respectively, and the coefficient matrix *B* is constructed as:

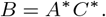

Lastly, the observed response matrix *Y ∈* ℝ^*nxq*^ is generated as

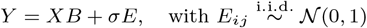

where the standard deviation *σ* defines the signal-to-noise ratio so that

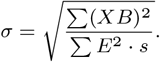

We considered the signal-to-noise (s2n) values *s* = 0.1, 1, 10.

This simulation setting corresponds to equation (1), where the true *B* matrix is obtained as the product of two low-rank matrices. It can be used to evaluate the performance of the Ridge RDA estimator in terms of i) recovering the latent dimension of the coefficient matrix, ii) extremely high-dimensional computing, and iii) prediction accuracy on new data.

### 2.7. Simulation scenarios

The performance of our ridge redundancy analysis procedure, hereafter referred to as RRDA, was evaluated using 3 different criteria: computational performance (Scenario 1), rank estimation (Scenario 2), and prediction performance (Scenario 3).

#### Scenario 1: Computational performance

To assess the computational effciency of the newly implemented procedure in high-dimensional settings, we evaluated the computation time of our `rrda.fit` function from the R package `rrda` and compared it with the conventional `rrs.fit` function from the R package `rrpack`. While `rrs.fit` is not designed for high-dimensional settings, it is the only available alternative implementation of RRDA. For both `rrs.fit` and `rrda.fit`, we fixed parameters: *λ* = 1 and *r* = 5. We also included `sRDA` (Csala et al., 2017) as a non-sparse setting (nonzero = *p*) with *λ* = 1 and the number of latent variables set to 5. For simulations, we generated data based on Model 1, with the parameters *n* = 100 and *k* = 5, and tested several values *p* = *q* ∈ {10^2^, 10^3^, 10^4^, 10^5^, 10^6^}. The computation time for a single model fitting was recorded 5 times. In addition, peak memory usage during model fitting was recorded using R’s built-in garbage collector. See Table 1S for session details.

#### Scenario 2: Rank estimation

Rank estimation performance was evaluated on Models 1 and 2. The matrix dimensions were set at *p*=*q* ∈ {50, 100, 200, 500}, and the number of samples set at *n* ∈ {100, 200}. The true rank *k* was chosen from {2, 5, 10}. For each setting, simulation replicates were conducted 20 times. We compared the performance of the following procedures: information criteria applied to RRDA, information criteria applied to ANN, and `StARS` approach. A 5-fold CV was applied to either classic non-ridge RDA (`c-rda.cv`) for selecting rank, or RRDA (`rrda.cv`) for selecting both the rank and the ridge penalty parameter.

#### Scenario 3: Prediction performance

The predictive performance of the different parameter tuning methods was evaluated. We fixed the parameters at *n* = 100, *p* = 200, *q* = 200, and *k* = 5, using both Model 1 and Model 2. The data were randomly split into a training set (*n*_train_ = 90) and a test set (*n*_test_ = 10), and this procedure was repeated 100 times. Within each training set, the tuning parameters were selected by one of the following two strategies: (i) internal 5-fold cross-validation (CV), applied either to classic non-ridge RDA (`c-rda.cv`, selecting only 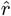) or to RRDA (`rrda.cv`, jointly selecting 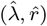); and (ii) information criteria (AIC, BIC, GIC, GCV, BICP) and `StARS`, applied to the ANN procedure (denoted `ANN.AIC`, `ANN.BIC`, etc.) to select the rank 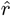. The qualifier “internal” indicates that, in both strategies, parameter selection is performed solely within the training split to avoid data leakage and to ensure a fair comparison across tuning methods. The model fitted with the tuned parameters was then applied to the test set, and predictive performance was quantified by the mean squared prediction error (MSPE):

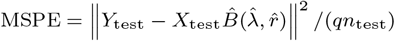

and averaged across the 100 splits. Note that, for ANN-based methods, which do not incorporate a ridge penalty, the fitted model is a reduced-rank regression with the selected rank 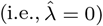. See Appendix C for details of the evaluation protocol.

### 2.8 Application to high-dimensional omics data

Three publicly available omics datasets were considered for the evaluation of the RRDA procedure (see Appendix A for details):

- The Breast Cancer dataset (Chin et al., 2006) for investigating the relationship between gene expression profiles and copy number variations (CNVs) in patients. Two versions were analyzed: a chromosome 13 subset (*X* ∈ ℝ^89*×*319,^ *Y* ∈ ℝ^89*×*58^) (also used in Wen et al. 2023), and a whole-genome set (*X* ∈ ℝ^89*×*19,672,^ *Y* ∈ ℝ^89*×*2,149^).
- The Soybean dataset of Dang et al. (2025) for investigating the relationship between microbiome and metabolome in the plant rhizosphere, where *X* ∈ ℝ^179*×*4771^ represents microbiome data, and *Y* ∈ ℝ^179*×*253^ represents metabolome data.
- The Cancer Genome Atlas (TCGA) methylation and expression dataset (Colaprico et al., 2016) to evaluate the association between DNA methylation and gene expression. We define *X* ∈ ℝ^20*×*392,277^ as DNA methylation and *Y* ∈ ℝ^20*×*67,528^ as gene expression.

For the Breast Cancer and Soybean datasets, the same MSPE evaluation framework as described in Simulation Scenario 3 was applied: the data were randomly split into training and test sets 100 times, regularization parameters 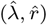 were tuned via internal 5-fold CV within each training set, and MSPE was evaluated on the held-out test set and averaged across the 100 random splits. The training/test splits were set to *n*_train_ = 79 and *n*_test_ = 10 for the Breast Cancer dataset, and *n*_train_ = 159 and *n*_test_ = 20 for the Soybean dataset. For the TCGA dataset, due to its limited sample size (*n* = 20), a leave-one-out cross-validation strategy was adopted instead, where each sample was held out once as the test set, resulting in 20 iterations with *n*_train_ = 19 and *n*_test_ = 1. See Appendix C for full details of the evaluation protocol.

For visualization purposes, a whole-sample estimate 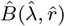 was obtained by refitting the model on the entire dataset, with 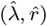 selected by 5-fold CV on the same full sample. This estimate was then visualized as a heatmap through the reduced submatrix 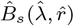 defined in Equation (10), with *m*_*x*_ = *m*_*y*_ = 30. The top 30 predictor and response variables were selected based on the row-wise *ℓ*_2_ norms of the low-rank factor matrices 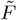 and 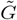, highlighting the most influential feature associations in the *r*-dimensional space.

## 3. Results

### 3.1. Simulation Results

#### Visualization

We first illustrate the parameter selection strategy implemented in the `rrda` R package, using simulation models 1 and 2 with parameters *n* = 100, *p* = *q* = 200, and *k* = 5. Similarly to the classical output in ridge regression, the regularization path can be visualized through a graph representing the cross-validated mean squared error (CV-MSE) as a function of the (log) value of the regularization parameter *λ*, as shown in Figure 1. Note that each curve corresponds to a candidate rank value *r*. The optimal combination 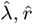 is then obtained as the location of the minimum among all curves, corresponding to the left vertical dotted line on the graph. The *λ* value corresponding to the one-standard-error rule is also displayed (right vertical dotted line). In what follows, only the results obtained with 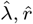 are displayed, as the one-standard-error rule did not show any improvement (Table 9S).

#### Computational performance

Table 3 displays the computational times (obtained on a single CPU core) and peak memory usage of 3 different implementations of the Ridge RDA, corresponding to R packages `sRDA`, `rrpack`, and `rrda`, respectively. The computational time corresponds to a single fit of the model with a rank set to 5 and a predetermined lambda value of 1. For mild values of (*p, q*) = 10^3^, `rrda` achieves a ×100 speed-up over its best competitor. For higher values of (*p, q*) ≥ 10^4^, the computational times become prohibitive for both `sRDA` and `rrpack` while remaining lower than 5 seconds for `rrda`. At (*p, q*) = 10^6^, conventional implementations in `rrpack` and `sRDA` required over 7,450 GB of memory to form the *p* × *p* matrix *X*×*X*, rendering computation infeasible. In contrast, `rrda` performed the task in 45.66 seconds. See Table 1S for session details.

**Table 3.**
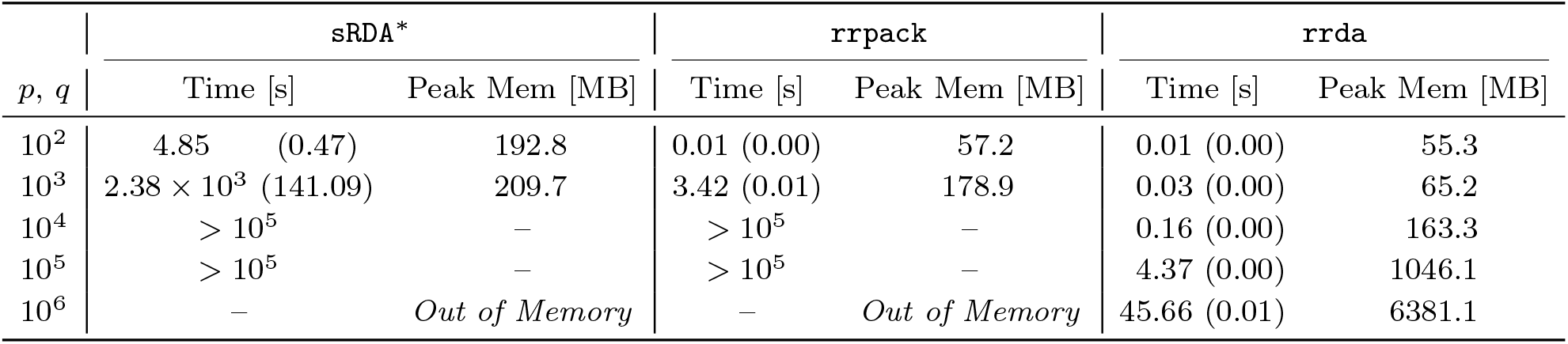
Benchmark of computation time and peak memory usage for procedures sRDA^*\**^, rrpack (using function rrs.fit), and rrda, based on Scenario 1. Computation time is reported as mean (standard error) in seconds [s]; peak memory is in megabytes [MB]. Peak memory is the maximum vector memory in use at any point during execution. “*>* 10^5^” indicates that the computation did not converge within the time limit (10^5^ seconds), in which case peak memory could not be recorded (denoted “–”). “*Out of Memory*” indicates that the computation failed due to exceeding the memory limit on the computational server (see Table 1S); at (*p, q*)= 10^6^, `sRDA` and `rrpack` required at least 7,450 GB of memory. ^*\**^`sRDA` is configured in a non-sparse setting of the Elastic Net to approximate a ridge prediction model.

#### Rank estimation

Figure 2 displays the behavior of classic criteria or CV-MSE (Y-axis) across a rank grid (X-axis), with the estimated rank indicated in red and the true rank in blue. The parameters were set to *n* = 100, *p* = *q* ∈ {50, 100, 200}, *k* = 5, and *s*2*n* = 1 (mid-noise) with the best *λ* selected for each rank. As shown, all of the tested classic criteria (AIC, BIC, GIC, BICP, GCV) severely overestimated the rank due to overfitting, as indicated by the sum of squared errors (SSE) converging to zero. In contrast, `rrda.cv` provided a reliable rank estimation. Similar patterns were observed across broad parameter settings (Tables 2S, 3S, and 4S).

**Fig. 2:**
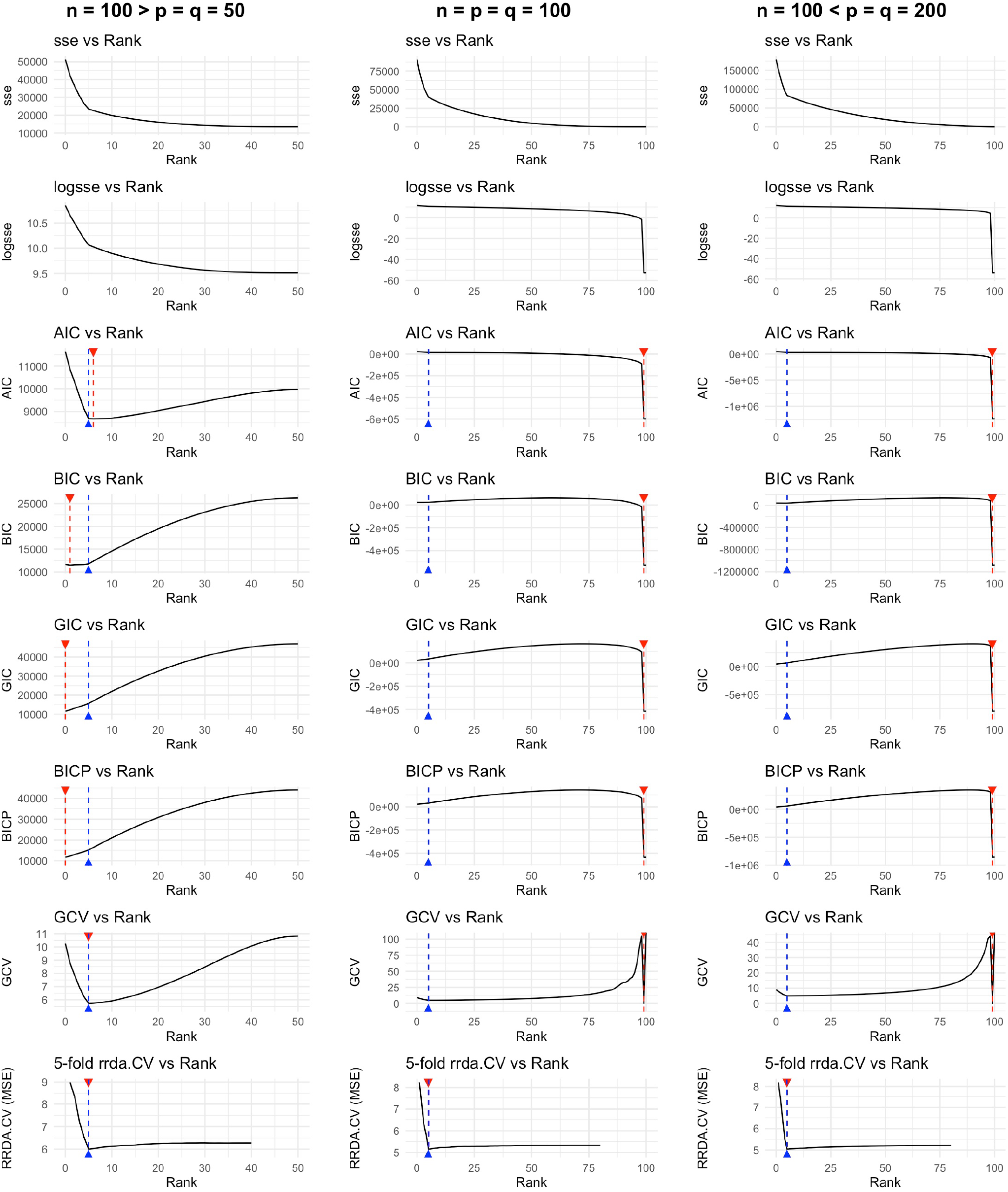
Performance of different information criteria (in rows) in different simulation settings (in columns), computed on data simulated using Model 1 (mid-noise: s2n = 1). The first two rows correspond to 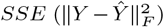 and log(*SSE*), the remaining ones corresponding to penalized criteria (AIC, BIC, GIC, BICP, GCV) and cross-validation (`rrda.cv`). Each graph displays the evolution of a specific criterion (y-axis) with respect to the rank (x-axis). For each rank value the best λ is selected. The true rank is indicated by a blue upward triangle and dashed vertical line, while the rank selected by the current criterion is indicated by a red downward triangle and dashed vertical line.

Next, we evaluated the scenario where information criteria are applied to ANN (information criterion-based ANN), comparing `StARS` with RRDA-based CV. Although these information criterion-based ANN methods were more stable than the classic grid search of information criteria, overfitting was still observed. Especially when *n* = *p* = *q* = 100 or 200, `ANN.BIC`, `ANN.BICP`, `ANN.GIC`, and `ANN.GCV` showed extreme overestimation of the rank (Figure 2S).

For a detailed comparison between information criteria-based ANN methods and RRDA-based CV, the results of the parameter setting *n* = 100, *p* = *q* = 200 are presented in Table 4. `ANN.AIC` and `StARS` tended to overestimate the rank, while `ANN.BIC`, `ANN.GIC`, and `ANN.BICP` often underestimated it. Especially in high-noise settings (*s*2*n* = 0.1), `ANN.BIC`, `ANN.GIC`, and `ANN.BICP` returned an estimated rank of 0, indicating failure to detect any signal. A broader landscape across matrix sizes is provided in Tables 5S, 6S, and 7S. In low-dimensional settings, `ANN.AIC` and `StARS` yielded relatively more accurate rank estimates, consistent with Wen et al. (2023). In contrast, in high-dimensional settings, both `ANN.AIC` and `StARS` consistently estimated the rank at its maximum value. See also Appendix E for details.

**Table 4.** Rank estimation performance of procedures ANN and RDA, based on Scenario 2 with *n* = 100, *p* = *q* = 200 and considering different signal-to-noise ratios (s2n), models (1 or 2) and true rank values (2, 5, 10). Estimated rank values are averaged across iterations and displayed with standard errors in parentheses.

|  |  |  | ANN |  |  |  |  |  | RDA |  |
|---|---|---|---|---|---|---|---|---|---|---|
| **s2n** | **Model** | **rank (*k*)** | **AIC** | **BIC** | **GIC** | **BICP** | **GCV** | **StARS** | **c-rda.cv** | **rrda.cv** |
| **0.1** | Model 1 | 2 | 99.00 (0.00) | 0.00 (0.00) | 0.00 (0.00) | 0.00 (0.00) | 2.40 (0.18) | 99.00 (0.00) | **2.00** (0.00) | **2.00** (0.00) |
|  |  | 5 | 99.00 (0.00) | 0.00 (0.00) | 0.00 (0.00) | 0.00 (0.00) | 6.95 (0.25) | 99.00 (0.00) | 1.25 (0.10) | **5.15** (0.08) |
|  |  | 10 | 99.00 (0.00) | 0.00 (0.00) | 0.00 (0.00) | 0.00 (0.00) | 7.45 (0.29) | 99.00 (0.00) | 1.00 (0.00) | **9.45** (0.59) |
|  | Model 2 | 2 | 99.00 (0.00) | 0.00 (0.00) | 0.00 (0.00) | 0.00 (0.00) | 2.35 (0.15) | 99.00 (0.00) | 1.60 (0.11) | **1.95** (0.05) |
|  |  | 5 | 99.00 (0.00) | 0.00 (0.00) | 0.00 (0.00) | 0.00 (0.00) | 6.95 (0.26) | 99.00 (0.00) | 1.25 (0.12) | **4.25** (0.29) |
|  |  | 10 | 99.00 (0.00) | 0.00 (0.00) | 0.00 (0.00) | 0.00 (0.00) | 7.00 (0.35) | 99.00 (0.00) | 1.00 (0.00) | **8.00** (0.90) |
| **1** | Model 1 | 2 | 99.00 (0.00) | **2.00** (0.00) | **2.00** (0.00) | **2.00** (0.00) | 2.30 (0.15) | 99.00 (0.00) | **2.00** (0.00) | **2.00** (0.00) |
|  |  | 5 | 99.00 (0.00) | 4.95 (0.05) | 0.00 (0.00) | 0.00 (0.00) | 5.05 (0.05) | 99.00 (0.00) | **5.00** (0.00) | **5.00** (0.00) |
|  |  | 10 | 99.00 (0.00) | 0.70 (0.18) | 0.00 (0.00) | 0.00 (0.00) | 10.30 (0.16) | 99.00 (0.00) | **10.00** (0.00) | **10.00** (0.00) |
|  | Model 2 | 2 | 99.00 (0.00) | **2.00** (0.00) | **2.00** (0.00) | **2.00** (0.00) | 2.30 (0.18) | 99.00 (0.00) | **2.00** (0.00) | **2.00** (0.00) |
|  |  | 5 | 99.00 (0.00) | **5.00** (0.00) | 0.00 (0.00) | 0.00 (0.00) | 5.05 (0.05) | 99.00 (0.00) | 4.90 (0.07) | **5.00** (0.00) |
|  |  | 10 | 99.00 (0.00) | 1.05 (0.18) | 0.00 (0.00) | 0.00 (0.00) | 10.35 (0.18) | 99.00 (0.00) | 9.45 (0.15) | **9.65** (0.15) |
| **10** | Model 1 | 2 | 99.00 (0.00) | **2.00** (0.00) | **2.00** (0.00) | **2.00** (0.00) | 2.25 (0.16) | 99.00 (0.00) | **2.00** (0.00) | **2.00** (0.00) |
|  |  | 5 | 99.00 (0.00) | **5.00** (0.00) | **5.00** (0.00) | **5.00** (0.00) | 5.10 (0.10) | 99.00 (0.00) | **5.00** (0.00) | 5.05 (0.05) |
|  |  | 10 | 99.00 (0.00) | **10.00** (0.00) | 0.00 (0.00) | 0.05 (0.05) | **10.00** (0.00) | 99.00 (0.00) | **10.00** (0.00) | 10.05 (0.05) |
|  | Model 2 | 2 | 99.00 (0.00) | **2.00** (0.00) | **2.00** (0.00) | **2.00** (0.00) | 2.40 (0.15) | 99.00 (0.00) | **2.00** (0.00) | **2.00** (0.00) |
|  |  | 5 | 99.00 (0.00) | **5.00** (0.00) | **5.00** (0.00) | **5.00** (0.00) | 5.40 (0.20) | 99.00 (0.00) | **5.00** (0.00) | **5.00** (0.00) |
|  |  | 10 | 99.00 (0.00) | **10.00** (0.00) | 0.00 (0.00) | 0.10 (0.07) | **10.00** (0.00) | 99.00 (0.00) | **10.00** (0.00) | **10.00** (0.00) |

**Table 5.** Scenario 3: The rrda evaluation for simulation data. The signal-to-noise (s2n) = 1. The estimations of rank, MSPE, *λ*, and Computation time of parameter tuning for random-splitted samples (100 iterations), and tested on the test data by MSPE 100 times. Comparison of model fit with various tuning parameters. CV (5-fold) is performed for the `rrda` model. Mean values are presented with standard error in the parentheses. For session details, see Table 1S.

|  | **ANN** |  |  |  |  |  | **RDA** |  |
|---|---|---|---|---|---|---|---|---|
|  | **AIC** | **BIC** | **GIC** | **BICP** | **GCV** | **StARS** | **c-rda.cv** | **rrda.cv** |
| **Model 1: Latent Space Model** |  |  |  |  |  |  |  |  |
| Rank | 89.00<br>(0.00) | 4.58<br>(0.05) | 0.00<br>(0.00) | 0.00<br>(0.00) | 5.00<br>(0.00) | 89.00<br>(0.00) | 5.00<br>(0.00) | 5.00<br>(0.00) |
| *λ* | -<br>- | -<br>- | -<br>- | -<br>- | -<br>- | -<br>- | -<br>- | 888<br>(7.61) |
| MSPE | 8.93<br>(3.41e-2) | 5.47<br>(3.96e-2) | 9.06<br>(7.78e-2) | 9.06<br>(7.78e-2) | 5.18<br>(1.72e-2) | 8.93<br>(3.41e-2) | 5.18<br>(1.72e-2) | **5.00**<br>(1.57e-2) |
| **Model 2: Matrix Factorization Model** |  |  |  |  |  |  |  |  |
| Rank | 89.00<br>(0.00) | 4.09<br>(0.03) | 0.00<br>(0.00) | 0.11<br>(0.03) | 5.00<br>(0.00) | 89.00<br>(0.00) | 4.78<br>(0.04) | 4.94<br>(0.02) |
| *λ* | -<br>- | -<br>- | -<br>- | -<br>- | -<br>- | -<br>- | -<br>- | 28.2<br>(1.00) |
| MSPE | 2.62<br>(1.64e-2) | 1.69<br>(9.90e-3) | 1.97<br>(1.59e-2) | 1.97<br>(1.58e-2) | 6.13<br>(4.44) | 2.62<br>(1.64e-2) | 6.13<br>(4.44) | **1.62**<br>(9.30e-3) |

RRDA-based CV methods were the most reliable compared with other methods. Notably, `rrda.cv` outperformed `c-rda.cv`, especially in high-noise settings (*s*2*n* = 0.1), highlighting the role of ridge regularization in accurate rank estimation.

#### Prediction performance

Table 5 displays the rank estimation and the prediction accuracy (computed as MSPE using random data splitting) on Scenario 3. Across both Model 1 and Model 2, `rrda.cv` achieved the lowest MSPE among all competing tuning strategies, including the information criteria and `StARS` applied to ANN. Among the ANN-based methods, `ANN.GCV` performed best under Model 1 and `ANN.BIC` under Model 2, yet both remained inferior to `rrda.cv`. The `rrda.cv` formula also clearly outperformed `c-rda.cv` by incorporating the ridge penalty. Similar trends were observed under various noise conditions (Table 8S). Using our newly implemented procedure, the computation time of CV was less than 5 seconds for all settings. The RRDA also outperformed a standard ridge regression and a multivariate linear regression without regularization (Table 9S).

### 3.2. Application Results

#### 3.2.1. Breast cancer data

We first evaluated the performance of `rrda` on the chromosome 13 subset (*X* ∈ ℝ^89*×*319^, *Y* ∈ ℝ^89*×*58^), using 100 random splits of the dataset (*n*_train_ = 79, *n*_test_ = 10). Hyperparameter tuning was conducted via internal 5-fold CV within each training split. We also considered the criterion-based ANN methods and `StARS`, as described in Wen et al. (2023). Contrary to their findings, `c-rda.cv` outperformed `StARS` in terms of both rank estimation and prediction accuracy. Across all tested methods, `rrda.cv` yielded the best prediction accuracy (Table 10S).

The `rrda` was then applied to the whole-genome dataset (*X* ∈ ℝ^89*×*19,672^, *Y* ∈ ℝ^89*×*2,149^) for scalability assessment. Among all methods, `rrda.cv` yielded the lowest MSPE. The `rrda` procedure completed inference in ~ 40 seconds using a single core and ~ 6 seconds using parallel computing (Table 13S). In comparison, `StARS` completed inference in ~ 6 minutes, with all other criteria requiring 20 seconds or more each.

RRDA was further compared to sparse RDA (`sRDA`; Csala et al. 2017) on the whole-genome dataset. Due to the high computational burden of `sRDA`, we limited its evaluation to 5 random splits (train/test = 79/10). For both methods, hyperparameter tuning was conducted via internal 5-fold CV within each training split. The parameter grid was designed to balance computational feasibility with effective coverage (see Figure 4S). For `sRDA`, parameters were tuned over the predefined grid to maximize the sum of absolute correlations, as proposed by Csala et al. (2017). We also tested MSE-based tuning, which yielded similar performance.

As summarized in Table 11S, `rrda` consistently achieved lower MSPE and significantly faster computation. While `sRDA` with Elastic Net required over 24 hours to complete five splits, `rrda` completed in under 2 minutes on a single core.

Additionally, we investigated the prediction performance at the chromosome-level in the whole-genome scenario. Two approaches were compared: (i) *Chr-Chr*, where gene expression values from each chromosome were used to predict CNV values on the same chromosome; and (ii) *All-Chr*, where gene expression values from all chromosomes were used to predict CNV values on each individual chromosome. MSPE was evaluated via 100 random-splitting (train/test = 79/10). The results indicate comparable performance for both approaches (Figure 3a). Alternatively, the chromosome-level scenario was tested in the inverted setting, i.e., using CNV as predictors and gene expression as response variables to evaluate how CNV affects gene expression. Figure 3b indicates that the *All-Chr* approach yielded more accurate predictions than the *Chr-Chr* approach in the inverted setting.

**Fig. 3:**
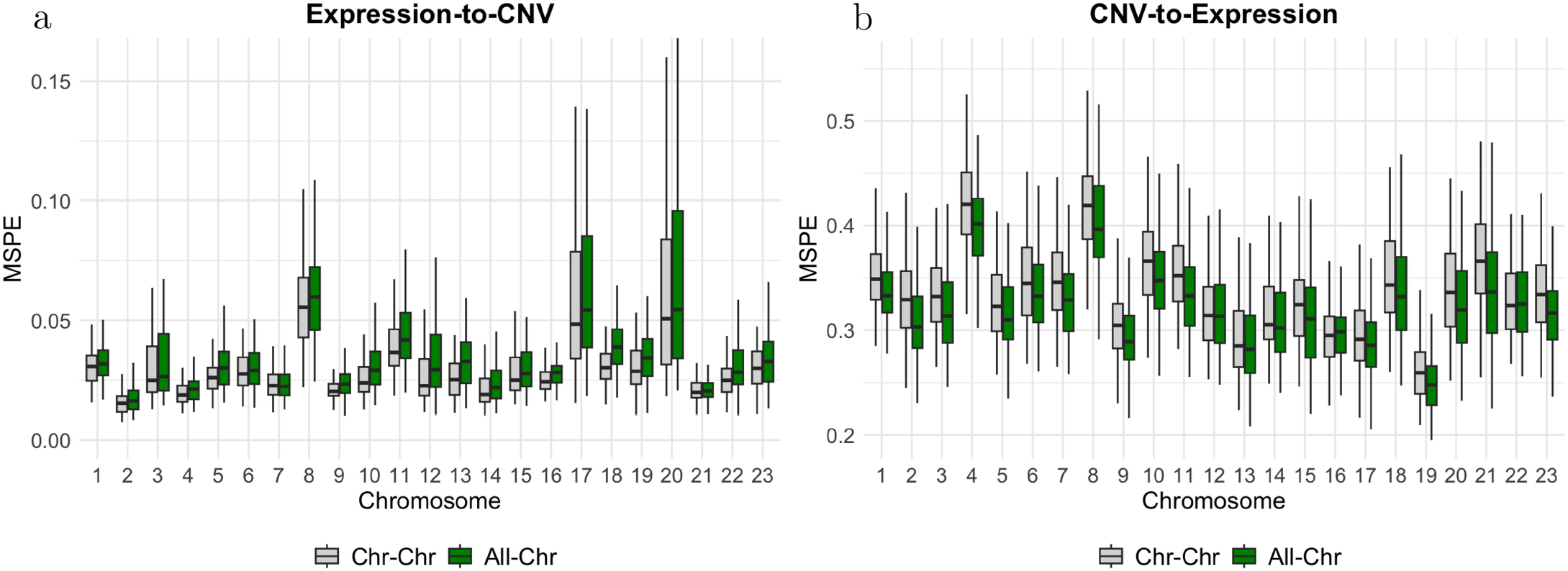
Chromosome-wise prediction performance for the Breast Cancer application, over 100 splits of the initial dataset into a training set (n=79) and a test (n=10) set. Within each training set, regularization parameters 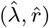 were tuned via internal 5-fold CV. The average MSPE is shown for each chromosome under two modeling strategies: *Chr-Chr* (gray), where the model was trained using data from the corresponding chromosome only, and *All-Chr* (green), where the model was trained using whole genome data. Two prediction settings were evaluated: (a) Expression-to-CNV, where gene expression data were used to predict CNV values, and (b) CNV-to-Expression, where CNV data were used to predict gene expression values.

To gain further biological insight, we performed CV on the entire sample set, which yielded the estimated coeffcient matrix 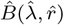 with the optimally selected parameters 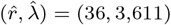. This result was visualized as a heatmap to highlight CNV–gene expression interactions (Figure 3S). The model identified several genes, such as *CEACAM5* and *SCGB2A2*, which are known to be overexpressed in breast cancer carcinomas (Sjödin et al., 2008; Bechmann et al., 2020). These genes showed strong correlations with clusters of CNVs located on chromosomes 11 and 17.

#### 3.2.2. Soybean multi-omics data

For the Soybean dataset (*X* ∈ ℝ^179×4771^, *Y* ∈ ℝ ^179×253^), we conducted 100 random-splitting (*n*_train_ =159,*n*_test_ = 20). Over the splits, internal CV tended to select a rank between 10 and 12 (Figure 4a), and RRDA yielded the lowest MSPE among all tested methods (Table 12S).

**Fig. 4:**
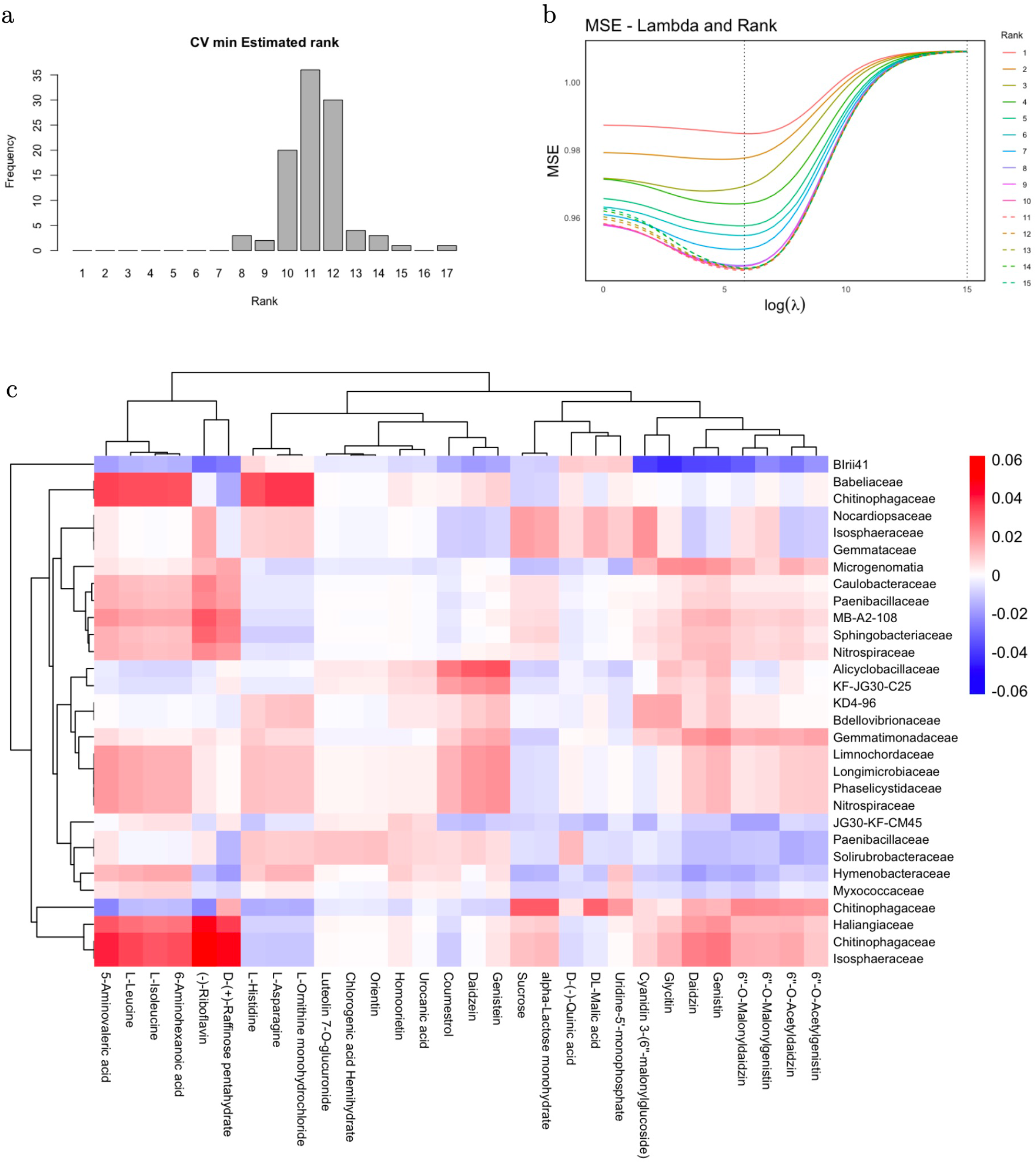
Application of `rrda` to the Soybean multi-omics dataset. (a) Empirical distribution of the optimally selected rank 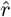 over 100 random splits (*n*_train_ = 159, *n*_test_ = 20). Parameters were tuned via internal CV in each split for performance evaluation. (b) Regularization path of the MSE as a function of tuning parameters *r* and *λ*, estimated via 5-fold CV using the entire sample set (*n* = 179). (c) Heatmap of `rrda` coefficients for the microbiome (rows) and metabolome (columns). The coefficient matrix 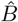 was estimated using the entire sample set with the best-tuned parameters from the CV (*r* = 11, *λ* = 335.74). Only the top 30 variables from each domain are presented, selected based on the *ε*_2_-norm of their low-rank factors.

Additionally, the tuning process via 5-fold CV using the entire sample set is illustrated by the regularization path in Figure 4b. Based on this procedure, the resulting estimated coefficient matrix 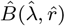 was obtained and is visualized as a low-dimensional representation in Figure 4c. This representation highlights structured associations between microbiome and metabolome features, where specific microbial families are linked to distinct metabolome profiles. Consistent with the analysis of Dang et al. (2025), the `rrda` method identified the *Chitinophagaceae* family (Carrión et al., 2019) as a key microbial variable. As illustrated in Figure 4c, the `rrda` method further identified microbe–metabolite associations. In particular, *Paenibacillaceae* showed positive correlations with soybean-derived aglycones (daidzein, genistein) and negative correlations with their glycosylated forms (daidzin, genistin). This pattern reflects the known ability of *Paenibacillus* to convert glycosides into bioactive aglycones (Park et al., 2013), highlighting its role in shaping the rhizosphere metabolome.

#### 3.2.3. TCGA methylation and expression data

The TCGA dataset (*X* ∈ ℝ^179×4771^, *Y* ∈ `ℝ`^179×253^) was tested using a leave-one-out (LOO) approach. Parameter tuning was performed for each training set through internal 5fold CV. Note that alternative approaches (e.g., `rrpack`, `StARS`) required approximately 100 GB of memory, and were unable to handle matrices of this size, even on a computing server. `sRDA` was also unable to operate under a non-sparse setting (i.e., nonzero=p). Therefore, our approach was the only feasible method for conducting the RRDA on this dataset. The computation time for parameter tuning (5-fold CV) using the `rrda.cv` function was 16 seconds using parallel computing (Table 13S).

The average estimated rank was 5.10 ± 1.01, and the regularization parameter was λ = 6.17 × 10^3^ ± 1.17 × 10^3^, with an MSPE of 7.27 × 10^−2^ ± 0.95× 10^−2^ across LOO evaluations. All values are reported as mean ± standard error.

Following 5-fold CV on the full sample for parameter tuning, the top 30 features of methylation sites and gene expression in the estimated low-dimensional coefficient matrix 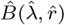 (Figure 5S) highlighted a group of methylation sites on chromosome X (e.g., `cg21983484`, `cg01742836`) that are strongly associated with specific gene expression (`TC0X001155.hg.1`, `TC0X002125.hg.1`) on the same chromosome. The observation is consistent with previously reported regulation, with `TC0X001155.hg.1` corresponding to a non-coding RNA known as *XIST*, which is known to play a critical role in X chromosome inactivation (Lee et al., 2019).

## 4. Discussion

The novel implementation of the Ridge Redundancy Analysis method, `rrda`, effectively addresses the computational challenges of high-dimensional settings and makes the procedure compatible with applications to large-scale datasets, including omics-integrative approaches as illustrated here.

Beyond scalability, the computational efficiency of our `rrda` procedure opened the way to efficient resampling strategies for the joint tuning of the regularization parameter ω and the rank r. Our study revealed the limitations of some classical information criteria for RRDA parameter tuning in high-dimensional settings. Indeed, whenever *p, q > n*, rank values larger than k can reduce the sum of squared errors to zero. The resulting overfitting is not compensated by the penalty term (Figure 2), leading to systematic overestimation of the rank. Although the ANN grid search partly mitigated this issue, its performance remained sensitive to dimensionality (Figure 2S). In both simulations and real data experiments, the *V*-fold CV of `rrda` led to improved rank estimation and yielded the best prediction performance compared to alternative selection criteria and competing models (Table 9S).

In our experiments, we observed a clear overestimation of the rank when using `StARS` in high-dimensional settings (Tables 5S, 6S, and 7S) that was not reported in the seminal paper. This overestimation may be partially explained by the implementation of `StARS` provided by Wen et al. (2023). In their implementation, `StARS` is first applied with the smallest threshold *v* - i.e., with the largest rank - and then moves to higher values of *v* until stability is reached via a sub-sampling method. However, in high-dimensional settings, when *r = n*_sampie_ (i.e., when the rank equals the sample size), the ANN model fits perfectly. This corresponds to the first “stable” rank estimate reached, which stops the grid search procedure and results in an overestimation of the rank (Figure 2S).

The various applications to real datasets illustrated the practical advantages of `rrda` compared to existing RDA methods. In the Breast Cancer application, `rrda` consistently outperformed `sRDA` both in terms of accuracy and computational efficiency (Table 11S). Although sparsity has often been advocated as leading to more interpretable models, it has been acknowledged that the identified support of variables with non-zero coefficients may be quite unstable under subsampling or slight changes of the experimental conditions, especially in high-dimensional settings (see e.g. Meinshausen and Bühlmann 2010; Baldassarre et al. 2017). In comparison, the `rrda` coefficient matrix in such settings was estimated to be relatively low-dimensional, resulting in a limited number of components to interpret and facilitating graphical representations, hence getting around the classical limitation of non-rank-restricted ridge procedures.

The Breast Cancer analysis illustrates that integrating whole-genome data (*All-Chr*) can improve prediction accuracy: Figure 3b demonstrates a clear improvement in prediction when CNV data are used to predict gene expression. This may reflect the trans-regulatory nature of CNVs, making genome-wide integration beneficial. More generally, having access to efficient and scalable whole-genome prediction procedures may be advantageous in scenarios where detailed genomic mapping is unavailable (Meuwissen et al., 2001; Bermingham et al., 2015).

Furthermore, the ability of `rrda` to offer a compact low-dimensional representation of feature associations was demonstrated in all the applications, where it revealed biologically meaningful structures across diverse omics layers (Figures 4c, 3S, and 5S). These structures, visualized through interpretable heatmaps, were obtained with minimal computational cost.

The RRDA framework we developed provides practical tools for multivariate data analysis in genomics and related fields, and is available as the R package `rrda` on CRAN.

## Acknowledgements

This work was supported by JSPS-KAKENHI (JP23KJ0506, JP22K21352), JST-CREST (JPMJCR16O2), JST-ALCA-Next (JPMJAN23D1), and JST-Mirai (JPMJMI120C7). This work has been supported by funds from the project AgroStat (reference: ANR-23-EXMA-0002) of the Maths-VivES France 2030 program handled by the French Agence Nationale de la Recherche. The authors thank the researchers who contributed to the soybean data collection: Y. Fuji, M. H. Yokota, Y. Ichihashi.

