## Appendix for "Ridge Redundancy Analysis for High-Dimensional Omics Data"

#### A. Details of Application data

##### A.1. General information

All datasets were used as obtained from their original public sources, in an already preprocessed format; consequently, no additional normalization or filtering was performed in this study. Note that the preprocessing step was therefore applied prior to cross-validation, which does not favor any particular procedure over the others in the benchmarking analysis.

For all three datasets, variable centering was conducted strictly within the CV framework to prevent information leakage, as implemented in the `rrda.cv` function: centering parameters (feature means) were estimated exclusively using the training data in each split or iteration, and these training-set statistics were then applied to center the corresponding test data.

##### A.2. Breast cancer data

We analyzed the Breast Cancer dataset (Chin et al., 2006). This dataset has been used in previous studies (Witten et al., 2009; Chen and Wang, 2022; Wen et al., 2023) to investigate the relationship between gene expression profiles and copy number variations (CNVs), with the aim of identifying potential therapeutic targets in breast cancer. Genome-wide copy number profiles were measured using bacterial artificial chromosome array comparative genomic hybridization, while gene expression profiles were obtained using Affymetrix U133A microarrays from primary breast tumors, as described in Chin et al. (2006).

The dataset comprises $n = 89$ patients, with gene expression measurements available for 19,672 transcripts, used as explanatory variables, and copy number variation measurements at 2,149 genomic loci, used as response variables.

Two versions of the breast dataset were analyzed: one limited to chromosome 13, where $X \in \mathbb{R}^{89\times319}$ and $Y \in \mathbb{R}^{89\times58}$, and another based on whole-genome data, where $X \in \mathbb{R}^{89\times19,672}$ and $Y \in \mathbb{R}^{89\times2,149}$.

##### A.3. Soybean multi-omics data

The Soybean multi-omics dataset consists of $n = 179$ samples, where microbiome amplicon sequence variants (ASVs) serve as explanatory variables $X \in \mathbb{R}^{179\times4771}$, and metabolome data serve as responses $Y \in \mathbb{R}^{179\times253}$, measured from different soybean accessions under drought conditions. It is known that the plant rhizosphere supports intricate communication and symbiosis between microbial communities and host metabolic processes. The motivation of this dataset is to investigate the relationship between the microbiome and metabolome in the plant rhizosphere.

As mentioned in Dang et al. (2025), microbiome profiles were obtained by sequencing the V4 region of the bacterial 16S rRNA gene using an Illumina MiSeq platform. Amplicons were purified, quantified, normalized, and pooled prior to sequencing. Raw reads were processed using QIIME2 (ver. 2020.6.0), including adapter trimming (Cutadapt), denoising and ASV inference (DADA2), read merging, and chimera removal. Taxonomic assignment was performed using the SILVA database (ver. 138). Non-bacterial sequences (Archaea, Eukaryota, mitochondria, chloroplasts) were removed.

Metabolome features were measured using liquid chromatography tandem mass spectrometry (LC–MS/MS) after extraction and purification from root tissues. This dataset was previously analyzed using a clustering method I-SVVS (Integrative Stochastic Variational Variable Selection) proposed by Dang et al. (2025).

In this study, we filtered out microbiome features with zero abundance across all samples. The Soybean multi-omics data is provided in the GitHub repository.

##### A.4. TCGA methylation and expression data

The methylation and expression dataset used in this study is derived from The Cancer Genome Atlas (TCGA), a landmark project that has publicly released a vast collection of clinical and molecular phenotypes from over 10,000 tumor patients across 33 cancer types (Colaprico et al., 2016). It is known that DNA methylation plays a key regulatory role in gene expression, making it a relevant target for integrative epigenomic analysis. TCGA has contributed to numerous discoveries by providing high-throughput molecular data, including DNA methylation and gene expression profiles. Colaprico et al. (2016) provide a guided workflow to query, download, and perform integrative analyses of TCGA data. The methylation-expression dataset contains data from $n = 20$ samples, with DNA methylation as input $X \in \mathbb{R}^{20\times392,277}$, and gene expression as output $Y \in \mathbb{R}^{20\times67,528}$. The motivation of this dataset is to evaluate whether DNA methylation patterns are associated with gene expression changes or external factors. Its extreme dimensionality ($p \gg n$ and $q \gg n$) presents a challenging test case for multivariate prediction methods.

This dataset is available via the Bioconductor package `brgedata` version 1.26.0 (Pelegri-Siso et al., 2024). DNA methylation and gene expression profiles were generated using high-throughput array-based and RNA-seq platforms, respectively.

#### B. Supporting Equations

##### B.1. Definition of Generalized Singular Value Decomposition

The GSVD for matrix $A$ with row and column metrics $LL'$ and $RR'$ is defined as follows

$$\mathcal{GSVD}(A)_{LL',RR'} = L'UDV'R$$

where $U$, $D$, and $V$ are the results of the SVD of $A$, and:

$$U'LL'U = V'RR'V = I$$

The GSVD results, $U^*$, $D^*$, and $V^*$, can be written as:

$$L'^{-}(\mathcal{SVD}(L'AR))R^{-} = UDV'$$

where

$$\mathcal{SVD}(L'AR) = U^*D^*V^{*\prime}$$

Thus,

$$U = (L')^{-}U^*, \quad V = (R')^{-}V^*, \quad D = D^*$$

For the process of GSVD, you can also refer to the description by Takane and Hunter (2001).

##### B.2. Computational Shortcuts When $p > n$

In optimizing the model, it is necessary to test multiple values for both the penalty parameter $\lambda$ and the rank. To efficiently handle this, we introduce a computational shortcut that reduces the dimension when the number of predictors $p$ exceeds the number of observations $n$. When $p > n$, computations can be performed in an $n$-dimensional space rather than a $p$-dimensional one, using the singular value decomposition (SVD). Geometrically, this corresponds to the fact that $n$ points in a $p$-dimensional space will always lie in an $(n-1)$-dimensional affine subspace.

Given the $n \times p$ data matrix $X$, the ridge regression estimate $\hat{B}(\lambda)$ in matrix form is given by:

$$\hat{B}(\lambda) = (X'X + \lambda I)^{-1} X'Y$$

This can alternatively be expressed as:

$$\hat{B}(\lambda) = X' (XX' + \lambda I)^{-1} Y$$

For the equation, we use the following theorem:

$$(A + BB')^{-1}B = A^{-1}B(I + B'A^{-1}B)^{-1}$$

Applying this, we replace $X'X$ with $XX'$, which allows us to reduce the computational cost of calculating $\hat{B}(\lambda)$ from $\mathcal{O}(p^3)$ to $\mathcal{O}(pn^2)$ when $p > n$ (Hastie et al. 2001, Section 18.3.5). The SVD of $X$ is written as:

$$X = UDV'$$

where $U$ is an $n \times n$ orthogonal matrix, $D$ is a diagonal matrix containing the singular values $d_1 \geq d_2 \geq \cdots \geq d_n \geq 0$, and $V$ is a $p \times n$ matrix with orthonormal columns. Using the SVD, we can express the inverse of $XX' + \lambda I$ as:

$$[XX' + \lambda I]^{-1} = [UD^2U' + \lambda UIU']^{-1} = U[D^2 + \lambda I]^{-1}U'$$

After some further manipulations, the solution can be rewritten as:

$$\begin{aligned}\hat{B}(\lambda) &= X' (XX' + \lambda I)^{-1} Y \\ &= VDU'U \left[D^2 + \lambda I\right]^{-1} U'Y \\ &= V \left(D \left[D^2 + \lambda I\right]^{-1}\right) U'Y\end{aligned}$$

This formulation leverages the SVD to significantly reduce computational complexity in scenarios where $p$ is large relative to $n$.

To simplify the formulations, we additionally define $P_X(\lambda)$ as follows:

$$P_X(\lambda) = X (X'X + \lambda P_{X'})^{-} X'$$

The $P_X(\lambda)$ is called the ridge operator. Using it,

$$Q_X(\lambda) = I - P_X(\lambda)$$

The objective is to reformulate the key components of the RRDA, including the projection matrices $P_{X'}$, $P_X(\lambda)$, and $Q_X(\lambda)$, the row metric for $LL' = X'X + \lambda P_{X'}$, the ridge estimated coefficient $\hat{B}(\lambda)$ using a computational trick.

Finally, we get the reduced-rank ridge estimate of $B$ by performing GSVD on $\hat{B}(\lambda)$ with all these elements. This reformulation facilitates the computation of the regularization path by exploiting the structural properties of the model to reduce the computational complexity associated with varying the regularization parameter $\lambda$ and the rank. Here, we show the proofs.

##### Matrix $P_{X'}$

A matrix $P_{X'}$ is defined as

$$P_{X'} = X' (XX')^{-} X = VV' \tag{12}$$

The proof proceeds as follows: Here, $X$ is an $n \times p$ matrix. We assume $n < p$ and $XX'$ is invertible. Assuming the orthogonality of the matrix $U$, where $UU' = U'U = I$, and $D$ is a diagonal matrix, the inverse of $XX'$ can be expressed as:

$$(XX')^{-} = \left(UD^2U'\right)^{-1} = UD^{-2}U'$$

Thus, we can simplify the projection matrix $P_{X'}$ as follows:

$$\begin{aligned}X' (XX')^{-} X &= VDU' \left(UD^2U'\right)^{-1} UDV' \\ &= VDU'UD^{-2}U'UDV' \\ &= VV'\end{aligned}$$

##### The ridge operator $P_X(\lambda)$

The ridge operator $P_X(\lambda)$ is defined as:

$$P_X(\lambda) = X (X'X + \lambda P_{X'})^{-} X' \tag{13}$$
$$= UD^2 \left(D^2 + \lambda I\right)^{-1} U' \tag{14}$$

The proof proceeds as follows: We assume that $\lambda > 0$ (or $\text{rank}(X) = n$). Utilizing the expression for $P_{X'}$ from (12), we can rewrite the inverse of $X'X + \lambda P_{X'}$ as:

$$\begin{aligned}(X'X + \lambda P_{X'})^{-1} &= \left(VD^2V' + \lambda VV'\right)^{-} \\ &= V \left(D^2 + \lambda I\right)^{-1} V'\end{aligned}$$

Substituting this into the expression for $P_X(\lambda)$, we obtain:

$$\begin{aligned}P_X(\lambda) &= X (X'X + \lambda P_{X'})^{-} X' \\ &= UDV'V \left(D^2 + \lambda I\right)^{-1} V'VDU' \\ &= UD^2 \left(D^2 + \lambda I\right)^{-1} U'\end{aligned}$$

This derivation highlights that the computation of the inverse is relatively inexpensive due to the diagonal structure of $D^2 + \lambda I$.

##### $Q_X(\lambda)$

The matrix $Q_X(\lambda)$ is given by

$$Q_X(\lambda) = U(\lambda I(D^2 + \lambda I)^{-1})U'$$

The proof proceeds as follows:

$$Q_X(\lambda) = I - P_X(\lambda)$$

According to (13),

$$P_X(\lambda) = UD^2(D^2 + \lambda I)^{-1}U'$$

Thus,

$$\begin{aligned}Q_X(\lambda) &= UU' - UD^2(D^2 + \lambda I)^{-1}U' \\ &= U(I - D^2(D^2 + \lambda I)^{-1})U' \\ &= U \left(\lambda(D^2 + \lambda I)^{-1}\right) U' \\ &= U \operatorname{diag}\left(\frac{\lambda}{d^2 + \lambda}\right) U'\end{aligned}$$

where we assume $\lambda > 0$. $d$ represents the vector of the diagonal elements of $D$.

##### Row Metric $LL'$

Row metric can be expressed using (12) as follows:

$$\begin{aligned} LL' &= X'X + \lambda P_{X'} \\ &= V(D^2 + \lambda I)V' \end{aligned}$$

Thus, the matrix $L$ and $L'$ can be written as:

$$L = V(D^2 + \lambda I)^{1/2} \tag{15}$$

$$L' = (D^2 + \lambda I)^{1/2} V'$$

The generalized inverse of $L'$ is derived as follows:

$$L'^{-} = V(D^2 + \lambda I)^{-1/2} \tag{16}$$

Here, $V$ is orthogonal.

##### Ridge Estimator $\hat{B}(\lambda)$

The ridge estimator $\hat{B}(\lambda)$ is given by:

$$\hat{B}(\lambda) = VD\left(D^2 + \lambda I\right)^{-1} U'Y$$

The proof proceeds as follows: According to (2) and (12)

$$\begin{aligned} \hat{B}(\lambda) &= (X'X + \lambda P_{X'})^{-} X'Y \\ &= (X'X + \lambda VV')^{-} X'Y \\ &= \left(VD^2V' + \lambda VV'\right)^{-} X'Y \\ &= \left(V(D^2 + \lambda I)V'\right)^{-} (UDV')'Y \\ &= \left(V(D^2 + \lambda I)^{-1}V'\right) VDU'Y \\ &= VD\left(D^2 + \lambda I\right)^{-1} U'Y \end{aligned}$$

##### GSVD for $\hat{B}$

The Generalized Singular Value Decomposition (GSVD) of matrix $\hat{B}(\lambda)$ is expressed as

$$\mathcal{GSVD}(\hat{B}(\lambda))_{LL',I} = V(D^2+\lambda I)^{-1/2}\mathcal{SVD}\left((D^2 + \lambda I)^{-1/2}DU'Y\right)$$

where $U$, $D$, and $V$ are matrices arising from the decomposition of $X$, and $SVD$ represents the singular value decomposition. The proof follows these steps: Using (15) and (16),

$$\begin{aligned} &\mathcal{GSVD}(\hat{B}(\lambda))_{LL',I} \\ &= L'^{-}(\mathcal{SVD}(L'\hat{B}(\lambda))) \\ &= L'^{-}(\mathcal{SVD}(L'VD\left(D^2 + \lambda I\right)^{-1} U'Y)) \\ &= V(D^2 + \lambda I)^{-1/2}\mathcal{SVD}\left((D^2 + \lambda I)^{1/2}V'VD\left(D^2 + \lambda I\right)^{-1} U'Y\right) \\ &= V(D^2 + \lambda I)^{-1/2}\mathcal{SVD}\left((D^2 + \lambda I)^{-1/2}DU'Y\right) \end{aligned}$$

#### C. Evaluation protocol for predictive performance

We evaluated predictive performance across both simulation and real-data analyses. In all experiments, we clearly distinguished between (i) internal CV used for hyperparameter tuning and (ii) validation for held-out samples for performance assessment, in order to avoid information leakage.

##### C.1. General evaluation framework

For each dataset, the data were split into training and test sets. Model parameters, including the ridge penalty $\lambda$ and rank $r$, were tuned exclusively within the training set using the internal 5-fold CV. The selected parameters $(\hat{\lambda}, \hat{r})$ were then used to fit the model on the full training data, and predictive performance was evaluated on the held-out test set.

The prediction accuracy was quantified using the mean squared prediction error (MSPE), defined as

$$\text{MSPE} = \left\| Y_{\text{test}} - X_{\text{test}}\hat{B}(\hat{\lambda}, \hat{r}) \right\|^2 / (qn_{\text{test}}),$$

where the error is normalized by the number of responses $q$. The reported performance corresponds to the average MSPE over repeated data splits.

Parameter tuning strategies included CV-based methods (e.g., `rrda.cv`, `c-rda.cv`), information criterion methods, ANN, and `StARS`, as described in Section 2.4.

##### C.2. Simulation scenario 3

In the simulation study (Scenario 3), we considered datasets with $n = 100$, $p = 200$, and $q = 200$. For each replicate, the data were randomly split into a training set ($n_{\text{train}} = 90$) and a test set ($n_{\text{test}} = 10$). This procedure was repeated 100 times.

For each split, hyperparameters $(\lambda, r)$ were selected using the internal 5-fold CV within the training set. The model was then trained on the full training data with the selected parameters and evaluated on the test set. The MSPE was computed for each split and averaged across repetitions.

##### C.3. Breast cancer data

For the Breast Cancer dataset ($n = 89$), we used a repeated random-splitting strategy. At each iteration, the data were randomly divided into a training set ($n_{\text{train}} = 79$) and a test set ($n_{\text{test}} = 10$). This process was repeated 100 times.

Within each training set, hyperparameters were tuned using an internal 5-fold CV. The final model, fitted using $(\hat{\lambda}, \hat{r})$, was evaluated on the corresponding test set. The predictive performance was summarized by the average MSPE across all splits.

##### C.4. Soybean multi-omics data

For the Soybean dataset ($n = 179$), we similarly performed 100 repeated random splits into training ($n_{\text{train}} = 159$) and test ($n_{\text{test}} = 20$) sets.

For each split, parameter tuning was conducted via an internal 5-fold CV within the training data. The model was then trained using the selected parameters and evaluated on the test set. The final performance was reported as the average MSPE across all repetitions.

##### C.5. TCGA methylation and expression data

Due to the limited sample size ($n = 20$), we adopted a leave-one-out (LOO) CV strategy. At each iteration, one sample was held out as the test set ($n_{\text{test}} = 1$), and the remaining $n_{\text{train}} = 19$ samples were used for training.

Within each training set, hyperparameters $(\lambda, r)$ were tuned using an internal 5-fold CV. The model was then fitted on the full training set using the selected parameters and evaluated on the held-out sample.

This procedure was repeated for all samples, and the final predictive performance was computed as the average MSPE across all LOO iterations.

### D. Sparse RDA model

#### D.1. Comparison with sparse RDA

We compared sparse RDA (`sRDA`, version 1.0.0, Csala et al. 2017) and `rrda`. While the two methods differ in formulation—`sRDA` is based on a partial least squares (PLS) framework with Lasso or Elastic Net regularization, and `rrda` uses ridge-regularized reduced-rank regression—they both target high-dimensional prediction modeling. For the scalability, `sRDA` promotes sparsity through penalization, while `rrda` leverages a low-rank structure via SVD. `sRDA` can approximate ridge-like behavior when the Elastic Net penalty is tuned toward its $\ell_2$ component (i.e., a non-sparse setting). However, even in this ridge-like configuration, the response of the function is not compatible with our `rrda` due to fundamental differences in their modeling approaches for rank reduction. This comparison highlights the methodological contrast between sparsity- and SVD-based approaches in high-dimensional settings, particularly in terms of computational efficiency in simulation (scenario 1) and predictive accuracy in application (Breast Cancer data).

#### D.2. Sparse RDA formulation

Let $Y \in \mathbb{R}^{n\times q}$ be the response matrix and $X \in \mathbb{R}^{n\times p}$ be the predictor matrix. We assume that all variables are centered and scaled.

Following the redundancy analysis framework of Csala et al. (2017), we define latent variables as linear combinations of $X$ and $Y$,

$$\xi = X\alpha, \qquad \eta = Y\beta,$$

where $\alpha \in \mathbb{R}^p$ and $\beta \in \mathbb{R}^q$ are weight vectors.

The objective of RDA is to determine $\alpha$ such that the correlations between $\xi$ and the columns of $Y$ are maximized. In the sparse RDA (sRDA) framework, we impose an elastic net penalty on $\alpha$ to handle high-dimensional settings and perform variable selection.

For fixed $\eta$, the weight vector $\alpha$ is estimated by

$$\hat{\alpha} = \arg\min_{\alpha} \left\{ \alpha' \left( \frac{X'X + \lambda_2 I}{1+\lambda_2} \right) \alpha - 2\eta' X\alpha + \lambda_1 \|\alpha\|_1 \right\}.$$

This formulation is equivalent to the elastic net regression problem

$$\min_{\alpha} \left\{ \|\eta - X\alpha\|^2 + \lambda_2 \|\alpha\|^2 + \lambda_1 \|\alpha\|_1 \right\},$$

up to a constant, obtained by expanding the squared loss. The factor $(1+\lambda_2)^{-1}$ serves as a normalization to ensure that the scale of the latent variable remains invariant with respect to the ridge parameter.

For fixed $\xi$, the response weights are estimated by ordinary least squares,

$$\hat{\beta} = (\xi'\xi)^{-1}\xi' Y.$$

The estimation proceeds iteratively by alternating updates:

$$\eta^{(t)} = Y\beta^{(t-1)},$$

$$\alpha^{(t)} = \arg\min_{\alpha} \left\{ \alpha' \left( \frac{X'X + \lambda_2 I}{1+\lambda_2} \right) \alpha - 2(\eta^{(t)})' X\alpha + \lambda_1 \|\alpha\|_1 \right\},$$

$$\xi^{(t)} = X\alpha^{(t)},$$

$$\beta^{(t)} = \left[ (\xi^{(t)})'\xi^{(t)} \right]^{-1} (\xi^{(t)})' Y.$$

The iterations are repeated until convergence, for example when

$$\sum_{k=1}^{p} \left( \alpha_k^{(t)} - \alpha_k^{(t-1)} \right)^2 < \varepsilon.$$

Multiple latent variables can be obtained sequentially by deflation. After extracting the $h$-th component $\xi_h$, the predictor matrix is updated as

$$X^{(h+1)} = X^{(h)} - \xi_h(\xi_h'\xi_h)^{-1}\xi_h' X^{(h)},$$

with $X^{(1)} = X$. The same procedure is then applied to $X^{(h+1)}$ and $Y$ to obtain the next component.

This deflation ensures that the latent variables $\{\xi_h\}$ are mutually orthogonal. In principle, up to $\min(n,p)$ latent variables can be extracted, although in practice the number is determined by a stopping criterion based on the additional variance explained in $Y$.

The first latent variable maximizes the total squared correlation with $Y$, while subsequent components explain progressively smaller portions of the variation in $Y$.

### E. ANN and StARS

#### E.1. Penalized Reduced-Rank Regression via Adaptive Nuclear Norm

This subsection outlines the reduced-rank estimation procedure using the adaptive nuclear norm (ANN) penalization (Chen et al., 2013).

ANN penalization is another approach to selecting the rank in multivariate models, implemented in the R package `rrpack`. ANN was recently extended to the `StARS` framework, and the formula is provided in Wen et al. (2023) (not a package). See Appendix G for details of the implementation.

The coefficient matrix $\mathbf{B} \in \mathbb{R}^{p\times q}$ is estimated via the following penalized optimization problem:

$$\widehat{\mathbf{B}}_\nu = \arg\min_{\mathbf{B}} \left\{ \frac{1}{2} \|\mathbf{Y} - \mathbf{XB}\|^2 + \nu \|\mathbf{XB}\|_w \right\}$$

where $\|\cdot\|^2$ is the squared Frobenius norm, and $\|\mathbf{XB}\|_w = \sum_{i=1}^{p\wedge q} w_i d_i(\mathbf{XB})$ is the adaptive nuclear norm with weights $w_i = d_i^{-\gamma}(\mathbf{PY})$, $\mathbf{P} = \mathbf{X}(\mathbf{X}'\mathbf{X})^{-1}\mathbf{X}'$, and $\gamma = 2$ by default.

The estimated rank of $\widehat{\mathbf{B}}_\nu$ is given by:

$$\widehat{r}_\nu = \max\left\{ r : d_r(\mathbf{PY}) > \nu^{1/(\gamma+1)} \right\}$$

$\nu$ is a thresholding parameter, not a ridge penalty parameter. The optimal value of $\nu$ determines the rank and final estimator $\widehat{\mathbf{B}}$. After defining the rank, the model can be tested through a standard reduced-rank regression model (RRR). Thus, the ANN is a kind of rank parameter tuning method.

While the ridge penalty shrinks (mostly) smaller singular values, the adaptive nuclear norm (ANN) approach removes singular values below the threshold $\nu$. Also, in our **rrda** approach, we define the rank $r$ (the number of singular values) explicitly.

#### E.2. Parameter range settings

Both the `rrpack` and `StARS` implementations define the same $\nu$ range for parameter selection. $\nu$ is a thresholding parameter, so it is not a ridge parameter.

Specifically, let $\nu_{\text{rrpack}}$ denote the set of $\nu$ values generated internally by the `rrpack` package for given data $(X, Y)$.

To explicitly compute the $\nu$ grid, we follow the strategy implemented in the `rrpack` package. This is equivalent to the following procedure: First, we obtain the fitted values of the ordinary least squares regression of $Y$ on $X$:

$$\hat{Y} = X\hat{B}_{\text{LS}}.$$

The candidate $\nu$ values are then generated based on the singular values of $\hat{Y}$. Specifically, we compute:

$$\text{Let } \hat{Y} = U_{\hat{Y}}\, \Sigma_{\hat{Y}}\, V_{\hat{Y}}^T, \quad \text{and let } \sigma_{\hat{Y}} = \text{diag}(\Sigma_{\hat{Y}}).$$

$$\text{Then} \quad Ad = \sigma_{\hat{Y}}^2, \quad Ad_x = Ad^{\frac{3}{2}}.$$

The sequence of $\nu$ values is then defined as:

$$\exp\Big(\text{seq}\big(\log(\max(Ad_x)),\ \log\left(Ad_{x\,[\min(n_{\text{rank}}+1,\ r_{\max})]}\right),\ 100\big)\Big),$$

where $n_{\text{rank}}$ is the maximum rank considered.

Putting this together, the resulting $\nu$ grid is:

$$\nu_{\text{rrpack}} = \{\nu_1, \ldots, \nu_{100}\}.$$

For the rank parameter, both `rrpack` and `StARS` are evaluated over the range up to $\min(\text{rank}(X), \text{rank}(Y), n_{\text{train}})$, where $n_{\text{train}}$ is a training sample size. For `rrda`, the default maximum rank is set to 15, which is generally sufficient as long as the estimated rank remains within this limit. This setting is reasonable because `rrda` is specifically designed for situations where the true rank of $B$ is much smaller than the matrix dimensions $n$, $p$, and $q$.

However, in cases where the estimated rank can be higher—such as in the Breast Cancer dataset, where `rrda.cv` produced optimal ranks greater than 15 (e.g., around 45)—we adjusted the maximum rank to $\min(\text{rank}(X), \text{rank}(Y), n_{\text{train}})$ accordingly. It is important to note that due to the use of CV in `rrda.cv`, the practically achievable maximum rank may be further constrained by the training sample size $n_{\text{train}}$, which is smaller than the total sample size $n$. The classic criteria parameter tuning implemented in `rrpack` was sensitive to the sample size, and overestimated without setting maxrank.

In both simulation and real data applications, we aim to fairly compare the performance of the proposed ridge redundancy analysis (`rrda`) with that of the existing `StARS` and `rrpack` methods. We ensure that any differences in performance arise solely from the methodological differences in estimation and rank selection.

#### E.3. ANN challenges in high-dimensional settings

The Adaptive Nuclear Norm (ANN) is a thresholding-based method used to estimate the rank of the coefficient matrix $B$. It applies a range of threshold values $\nu \in \mathcal{T}$ to the singular values of $B$, producing a set of candidate ranks $\mathcal{R} \subseteq \{1, \ldots, \min(n, p, q)\}$. In principle, each value of $\nu$ yields a corresponding rank estimate.

However, the ANN procedure does not directly suggest an optimal value of $\nu$ or rank for model selection. Therefore, a common strategy is to use model selection criteria—such as AIC, BIC, GIC, BICP, or GCV—to select the optimal rank from $\mathcal{R}$ in the context of reduced-rank regression (RRR). These criteria are implemented in the `rrr` function of the `R` package `rrpack` (version 0.1-13; Chen and Wang 2022), which provides ANN-based rank selection using each of the five criteria.

When a grid of 100 threshold values is used, the ANN approach provides up to 100 candidate rank estimates (note that different values of $\nu$ may yield the same rank). After simulations, rank estimation results are summarized in Tables 5S, 6S, and 7S.

These results show that in high-dimensional settings (e.g., $n = p = q = 100$ or 200), many classic information criteria (including BIC, GIC, BICP, GCV) occasionally overestimated the rank. This overestimation is further illustrated in Figure 2S, where the sum of squared errors (SSE; $\|Y - \hat{Y}\|^2$) converges toward zero as the model rank increases, causing log-likelihood–based criteria (which involve log(SSE)) to diverge. Because these criteria are inherently based on SSE, they are prone to overfitting—especially when the full-rank model achieves near-zero residual error.

While the ANN framework with grid-based search can sometimes avoid overfitting by chance—when the threshold does not suggest a maximal or near-maximal rank—this is not guaranteed. Thus, it depends on the grid generation. In certain cases, especially with large matrices, it may fail to prevent overestimation. Our results highlight the sensitivity of ANN methods to matrix dimensions and threshold values.

In summary, although criteria-based ANN methods are computationally convenient, they rely heavily on external thresholding and SSE-based criteria, which makes them unreliable in high-dimensional settings. Their tendency to overfit underscores the risks of applying classic information criteria for rank selection in such contexts.

#### E.4. StARS challenges in high-dimensional settings

The rank estimation pattern, shown in Figure 2S, highlights the systematic challenges of the `StARS` procedure in high-dimensional scenarios. `StARS` is a stability-based approach that estimates rank by measuring the variability across multiple sub-samples. The implementation can be found in the article of Wen et al. (2023). Specifically, it repeatedly performs rank estimation on random sub-samples of the data, by default, using 70% of the observations, and computes the variance of the results. This process is repeated (by default, 100 times) until a "stable" point is identified—defined as the smallest value of the thresholding parameter $\nu$ for which the variance of rank estimates falls below a predefined threshold (default: $10^{-4}$). In other words, the approach is not a grid search but "one-sided" search (from small $\nu$ to large $\nu$). The corresponding $\nu$ is then used to compute the final rank estimate on the full dataset.

While `StARS` performs well in low-dimensional settings, it encounters difficulties in high-dimensional cases (where $n < p, q$). Because `StARS` is dependent on the ANN implementation in the `rrr` function, it is prone to overfitting. `StARS` is a subsampling approach, thus it inherently reduces the number of observations below $n$, which imposes a cap on the maximum estimated rank.

During the search for stability, the procedure starts with small values of $\nu$, which tend to yield high-rank estimates. However, once the estimated rank hits the ceiling imposed by subsampling (and high-dimensionality), the rank may appear stable—even though this stability is artificial. This false stability leads `StARS` to terminate early and select a combination of a relatively small $\nu$ and an overly high rank, which does not reflect the true underlying structure. The systematic problem is that `StARS` is not a grid search, but a one-sided search approach, which makes it prone to getting stuck in local minima. While a full grid search typically incurs high computational costs, it offers a more exhaustive evaluation, presenting a trade-off between accuracy and efficiency. In contrast, our method, `rrda`, facilitates the computation and alleviates this issue.

The tendency of `StARS` to overestimate the rank was not explicitly discussed by Wen et al. (2023). In our study, we explored a wider range of both low-dimensional and high-dimensional settings and observed that `StARS` tended to overestimate the rank—particularly in high-dimensional scenarios—as demonstrated by our simulation results (Tables 4, 5, 5S, 6S, and 7S). While tuning specific hyperparameters may help control overfitting for specific cases, `StARS` may not be the most suitable choice for high-dimensional settings. In our simulations and applications, which focused primarily on high-dimensional cases, CV consistently outperformed `StARS` in both rank estimation and prediction accuracy.

##### E.5. CV and StARS

Using the Breast Cancer chromosome 13 dataset, we performed 100 random splits to reproduce the results for `rrpack` and `StARS` reported by Wen et al. (2023) (Table 10S). However, contrary to their findings, CV with c-rda.cv formula outperformed both `rrpack` and `StARS` in rank estimation and prediction accuracy.

We openly provide all data and code used in this study on GitHub, whereas Wen et al. (2023) did not make theirs publicly available, so the configuration they used may differ. The results are also likely affected by the choice of rank ranges evaluated. In our approach, the design of our function allows for efficient evaluation of a wide grid, facilitating the selection of the optimal parameter.

#### F. Top-Feature Visualization of the Ridge RDA Coefficient Matrix

The full RRDA coefficient matrix $\hat{B}(\lambda, r) \in \mathbb{R}^{p\times q}$ is often too large to visualize or interpret directly. For example, in the methylation application (Section 3.2), where $p = 392,277$ and $q = 67{,}528$, storing or visualizing the entire coefficient matrix $\hat{B}(\lambda, r)$ becomes computationally impractical.

To address this issue, the `rrda` framework allows one to extract a lower-dimensional submatrix by choosing the most informative features. This selection facilitates both interpretation and exploratory analysis while preserving the essential structure of the model.

In particular, the component form output by `rrda.fit` provides low-rank factor matrices that can be used to perform simple feature selection based on row-wise norms.

As described in Equation (8), the coefficient matrix admits the factorization $\hat{B}(\lambda, r) = FG'$, where $F \in \mathbb{R}^{p\times r}$ and $G \in \mathbb{R}^{q\times r}$ are defined in Section 2.3. Define the symmetrically scaled counterparts as

$$\tilde{F} = U^{[r]}_{\hat{B}(\lambda)} D^{[r]1/2}_{\hat{B}(\lambda)} \in \mathbb{R}^{p\times r}, \quad \tilde{G} = V^{[r]}_{\hat{B}(\lambda)} D^{[r]1/2}_{\hat{B}(\lambda)} \in \mathbb{R}^{q\times r},$$

so that $\hat{B}(\lambda, r) = FG' = \tilde{F}\tilde{G}'$, where the singular values are symmetrically distributed between the left and right factors.

To extract a manageable submatrix for visualization, define the row scores

$$s_i^{(\tilde{F})} = \|\tilde{F}_{i,\cdot}\|_2, \quad s_j^{(\tilde{G})} = \|\tilde{G}_{j,\cdot}\|_2,$$

where $\tilde{F}_{i,\cdot}$ and $\tilde{G}_{j,\cdot}$ denote the $i$-th and $j$-th rows of $\tilde{F}$ and $\tilde{G}$, respectively. The top $m_x$ predictors and top $m_y$ responses are selected according to the largest values of $s_i^{(\tilde{F})}$ and $s_j^{(\tilde{G})}$, yielding submatrices $\tilde{F}_s \in \mathbb{R}^{m_x\times r}$ and $\tilde{G}_s \in \mathbb{R}^{m_y\times r}$. The reduced coefficient matrix is then

$$\hat{B}_s(\lambda, r) = \tilde{F}_s\tilde{G}_s' \in \mathbb{R}^{m_x\times m_y}.$$

This submatrix retains the dominant associations captured by the low-rank estimator while remaining interpretable and computationally tractable. In the applications of Section 3.2.3, $\hat{B}_s(\lambda, r)$ is visualized as a heatmap with $m_x = m_y = 30$. This procedure is implemented in the `rrda.top` function of the `rrda` package.

#### G. Implementation

##### G.1. `rrpack` implementation for RRR calculation

ANN is implemented in the `rrr` function from R package `rrpack`. In `rrpack`, it has a specific implementation to avoid errors caused by inversion of the covariance matrix in the prediction. There are `rrr`, `cv.rrr`, `rrr.fit` functions for reduced-rank regression, and `rrs.fit` for ridge reduced-rank regression. `cv.rrr`, `rrr.fit`, and `rrs.fit` have an implementation to avoid the error by applying a minimal ridge penalty (which happens only when inverse calculation fails). These store the coefficient matrix B in p times q form, and therefore still require excessive memory in super-high-dimensional settings (i.e., over 100 GB in TCGA datasets).

The `cv.rrr` function is considered to be a special case of `rrda.cv` when $\lambda = 0$. In simulations and applications, we employed `rrda.cv` because `cv.rrr` becomes computationally burdensome in high-dimensional settings. In `rrr`, the lowest possible rank estimate is 0, corresponding to the mean model when it provides the best predictions. By contrast, `rrda` assigns a minimum rank of 1, which is advantageous for visualization and further implementation.

##### G.2. `StARS` implementation

The ANN is extended to `StARS` frameworks by Wen et al. (2023). `StARS` is not implemented as a package, but the functions are available in supplemental material of the article by Wen et al. (2023).

##### G.3. `rrda` software implementation and installation requirements

The `rrda` package was developed and tested in R version 4.4.1 (2024-06-14), primarily on Resource A described in Table 1S (Apple M2 Max, macOS Sonoma 14.5, aarch64). In addition, selected large-scale simulation experiments were also run under the same R version on Resource B (Ubuntu 24.04 LTS). The package is distributed on CRAN as `rrda` (version 0.2.3, published in Oct 2025).

Installation does not require compilation, and no additional external system dependencies are specified in the CRAN metadata. The package can therefore be installed directly from CRAN using standard R installation procedures:

```
install.packages("rrda")
```

For routine use, the package relies on standard R package dependencies listed on CRAN. These dependencies are installed automatically by `install.packages()` when needed. No non-R prerequisites are required for standard installation and use.

#### G.4. Fitting function rrda.fit

All statistical analyses in this study were conducted using R version 4.4.1 (R Core Team, 2023), and figures were produced using the R package ggplot2 version 3.5.1 (Wickham, 2016). Our fitting method is implemented in the R package "`rrda`." The function `rrda.fit` is designed to perform dimension reduction and regularization on two matrices, $X$ and $Y$ (predictor and response variables), typically in the context of high-dimensional multivariate analysis. The function allows flexibility in choosing the model via rank parameter *nrank* and regularization parameter $\lambda$, and whether to return individual components or the full approximated matrix. The function stores $\hat{B}(\lambda, r)$ in component form by default instead of matrix form. Additionally, the functions are designed for parallel computing relying on the `future` package (Bengtsson, 2021), which facilitates the computation for multiple parameters $\lambda$ and rank. Furthermore, in omics studies, the rank $r$ is often smaller than both $n$ and $p$, enabling efficient SVD computation of a limited number of top components using an optimized algorithm, such as the one implemented in the `RSpectra` package (Qiu and Mei 2024). To summarize, we present the `rrda.fit` method as in Algorithm 1.

#### G.5. Prediction function rrda.predict

The function `rrda.predict` is designed to perform predictions via given $X$ and estimated low-rank components $F$ and $G$. The rank ($r$) can be manually chosen. In most cases, the rank $r$ can be much smaller than the $p$ and $q$, reducing the computation task. To summarize, we present the `rrda.predict` method as in Algorithm 2.

#### G.6. Parallel Computing

All computation times, including those for simulation scenarios and applications, were first measured using single-core execution to ensure fair comparison with conventional methods. The corresponding results are reported for Scenario 1 (Table 3), the Lambda Sampling experiment (Fig. 1S), and Scenario 3 (Tables 8S and 9S). For application studies, results are shown for the breast cancer dataset (Table 10S), the comparison between `sRDA` and `rrda` (Table 11S), and the soybean dataset (Table 12S).

Additionally, parallel processing was evaluated for all application datasets for benchmarking purposes, and the results are presented in Table 13S.

When specifying `plan(multisession)`, computations are performed in parallel using the `furrr` package (Vaughan and M., 2022). For example:

**Algorithm 1** rrda.fit Algorithm

**Input:** Matrix of response variables $Y$; Matrix of predictor variables $X$; Rank parameter *nrank*; Regularization parameter $\lambda$; Scaling options (e.g., centering, scaling);

**Output:** List of low-rank components or estimated matrix $\hat{B}(\lambda, r)$;

Initialize *nrank*, $\lambda$, scaling parameters for $X$ and $Y$
**if** *nrank* is NULL **then**
  Set $nrank = \{1, \ldots, \min(15, \min(\dim(X), \dim(Y)))\}$
**end if**
Apply centering or scaling to $X$ and $Y$ if required
Perform SVD on $X$ to obtain $U$, $D$, and $V$
Compute $(D^2 + \lambda I)^{-1/2}$ for all $\lambda$
Estimate the low-rank matrix $\hat{B}(\lambda)$ using:

$$\hat{B}(\lambda) = VD(D^2 + \lambda I)^{-1}U'Y$$

Perform GSVD on $\hat{B}(\lambda)$ to obtain $U_{\hat{B}(\lambda)}, D_{\hat{B}(\lambda)}, V'_{\hat{B}(\lambda)}$
Store the low-rank components $F_{.i} = U_{\hat{B}(\lambda).i} D_{\hat{B}(\lambda).i}$ and $G_{.i} = V_{\hat{B}(\lambda).i}$ for $i = 1, \ldots, r$
**if** Component form is not required **then**
  Reconstruct matrix $\hat{B}(\lambda, r) = \sum_{i=1}^{r}(F_{.i} G'_{.i})$
**end if**
Return the list of $F$, $G$, or the matrix $\hat{B}(\lambda, r)$ for all $\lambda$

**Algorithm 2** rrda.predict Algorithm

**Input:** Ridge RDA components $\hat{B}(\lambda, r)$; Matrix of predictor variables $X$; Rank parameter *nrank*; Regularization parameter $\lambda$; Scaling information from training;

**Output:** Predicted matrix $\hat{Y}_r$ for each $\lambda$;

Initialize *nrank*, $\lambda$ from $\hat{B}(\lambda, r)$
**if** *nrank* is NULL **then**
  Set *nrank* as rank values stored in $\hat{B}(\lambda, r)$
**end if**
**if** $\lambda$ is NULL **then**
  Set $\lambda$ as $\Lambda$ values stored in $\hat{B}(\lambda, r)$
**end if**
**if** Training $X$ was centered or scaled **then**
  center and/or scale $X$ using the stored values from training data
**end if**
**for** each $\lambda$ in $\Lambda$ **do**
  Initialize an empty list to store the results for each rank
  **for** each rank $r = 1, \ldots, \max(nrank)$ **do**
    Compute $\tilde{X} = XF$
    Compute $\tilde{X}_{.i} G'_{.i}$ using the outer product to obtain $\tilde{Y}_{.i}$
  **end for**
  Use the `Reduce` function to sum over all ranks to generate $\hat{Y}_r$
**end for**
**if** Training $Y$ was centered or scaled **then**
  Re-center and/or re-scale $\hat{Y}_r$ using the stored values from training data
**end if**
Return the list of predicted matrices $\hat{Y}_r$ for all $\lambda$

```
# R code
set.seed(10)
```

```
X <- matrix(rnorm(10 * 30), 10, 30)
Y <- matrix(rnorm(10 * 30), 10, 30)

library(furrr)
plan(multisession)

cv_result <- rrda.cv(Y = Y, X = X, maxrank = 2, nfold = 5)

plan(sequential)
rrda.summary(cv_result = cv_result)
```

In this example, the CV is executed in parallel when `plan(multisession)` is active. After the computation, the execution plan is set back to sequential mode using `plan(sequential)`.

#### H. Data availability

All scripts used in this study, along with the breast cancer and soybean datasets, are publicly available on GitHub at `https://github.com/Yoska393/rrda`.

The TCGA methylation and gene expression data (Colaprico et al., 2016) are not hosted in the GitHub repository due to their size; however, instructions for downloading the data are provided in the repository. The data can be obtained via the `brgedata` package as follows:

```
if (!requireNamespace("BiocManager", quietly = TRUE))
    install.packages("BiocManager")

BiocManager::install(c(
  "brgedata",
  "IlluminaHumanMethylation450kanno.ilmn12.hg19"
))

library(brgedata)
library(IlluminaHumanMethylation450kanno.ilmn12.hg19)

data(brge_methy)
data(brge_gexp)
```

Details on dataset download procedures, simulation codes, and application codes are available in the GitHub repository (`https://github.com/Yoska393/rrda`).

#### Supplementary Figures and Tables 1523

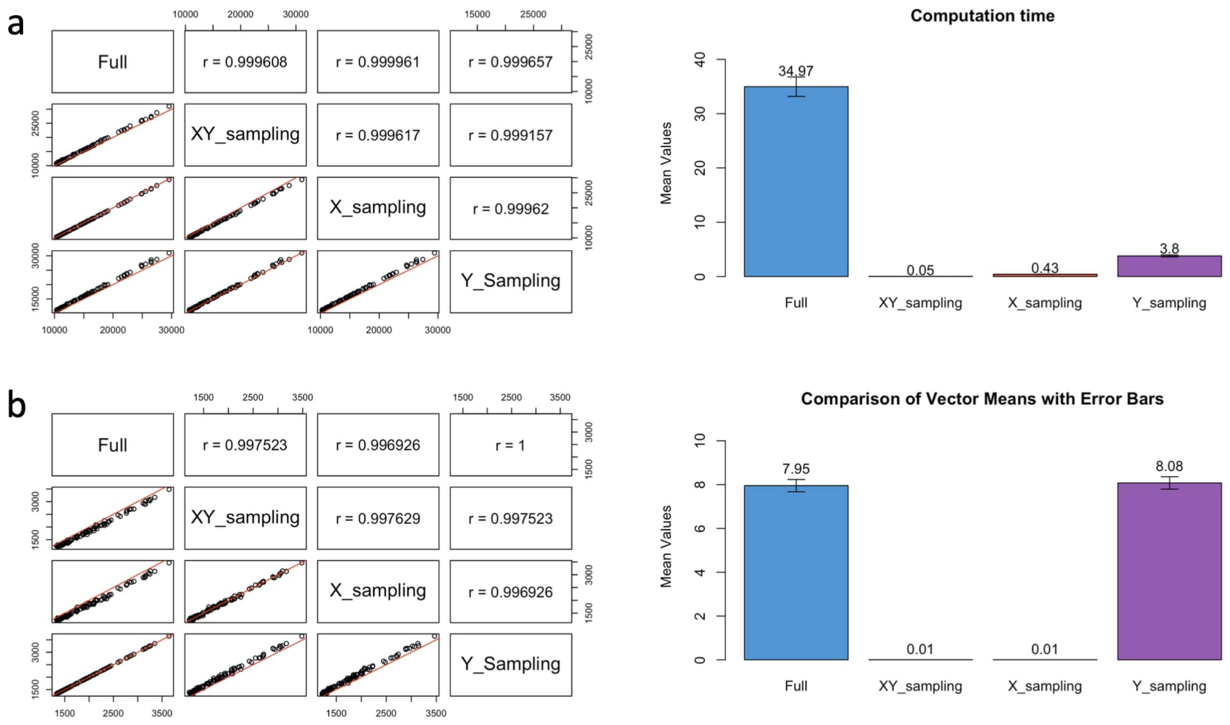


Fig. 1S: Simulation LS (Lambda Sampling). The results of the sampling method for estimating $\lambda_{\max}$. After generating the data sets on model 1 (Latent Space model), sampling of $10^3$ variables from the X and/or Y matrix was performed. Each point represents $\lambda_{\max}$ of each iteration. n is a random value from 10 to 100. The $\lambda$ estimated from the sampled matrices were compared with the one obtained from the full matrices, then Pearson correlation coefficient was calculated. The computation times given here are measured using sequential executions without parallel computing. (a) $p = 10^5$ and $q = 10^4$ (b) $p = 10^6$ and $q = 10^2$.

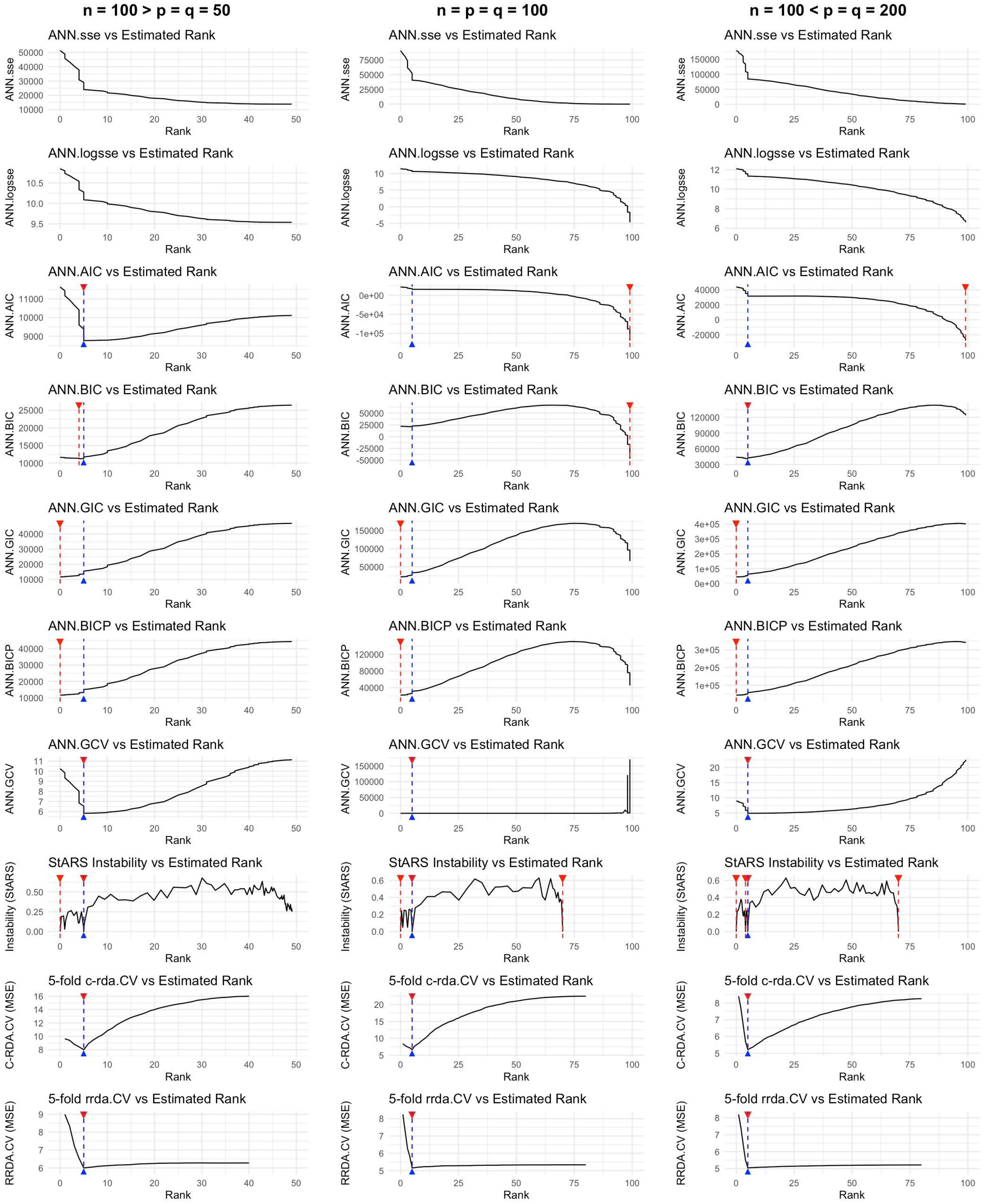


Fig. 2S: Rank vs ANN-based SSE (Sum of Squared Error), Log(SSE), information criteria. We also tested CV-MSE, which is not ANN based. The data were generated in each setting ($n = 100 > p = q = 50$, $n = p = q = 100$, $n = 100 < p = q = 200$) under Simulation Model 1 (mid-noise: s2n $= 1$). For SSE, logSSE, StARS, and all other criteria, the rank grid was generated based on ANN. For the CV approach, the rank grid was generated from 1 to $\min(n_{\text{train}}, p, q)$. Selection methods include the ANN based ANN.AIC, ANN.BIC, ANN.GIC, ANN.BICP, ANN.GCV, StARS criteria, and CV for both classic RDA (c-rda.cv) and ridge RDA (rrda.cv). The x-axis indicates the rank, and the y-axis shows the corresponding selection criterion to be minimized. StARS runs from the smallest $\nu$ (resulting in higher ranks) and proceeds toward largest $\nu$ (lower ranks) checking the rank estimate in subsample. The process immediately stops when instability drops below a threshold (default: $10^{-4}$). In each figure, the true rank is indicated by a blue upward triangle and dashed vertical line, while the estimated rank selected by the method is indicated by a red downward triangle and dashed vertical line. StARS displays all ranks selected based on stability, but the point furthest to the right is always selected as the estimated value of the function based on the algorithm.

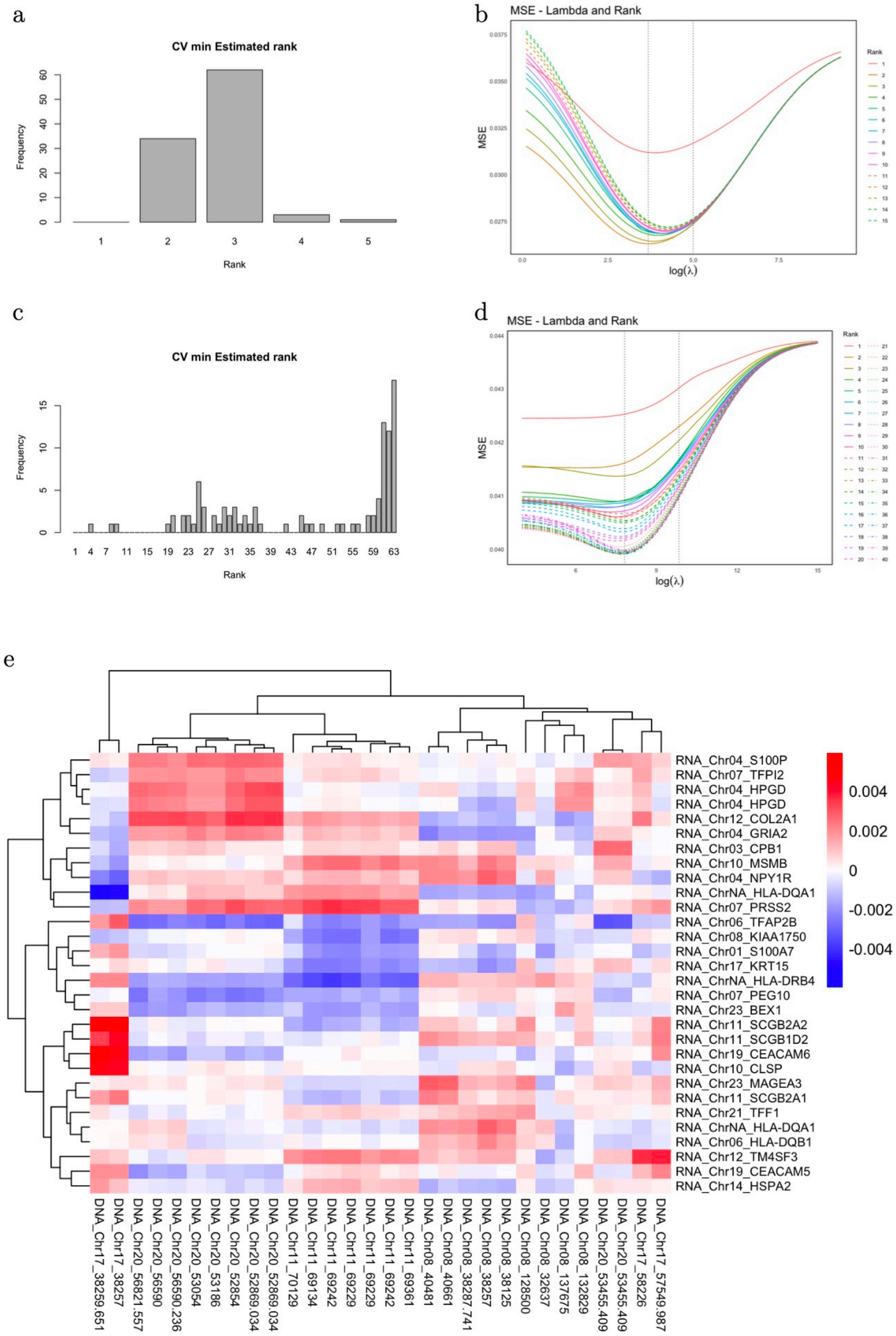


Fig. 3S: Application of `rrda` to the Breast Cancer dataset (Expression-to-CNV scenario). For the chromosome 13 subset ($n = 89, p = 319, q = 58$), (a) empirical distribution of the optimally selected rank $\hat{r}$ across 100 random splits ($n_{\text{train}} = 79, n_{\text{test}} = 10$) using internal CV, and (b) regularization path for parameter tuning via 5-fold CV using the entire sample set. For the whole-genome data ($n = 89, p = 19,672, q = 2,149$), (c) distribution of the selected rank $\hat{r}$ across 100 random splits using internal CV, and (d) regularization path for parameter tuning via 5-fold CV using the entire sample set. (e) Heatmap of `rrda` coefficients for the whole-genome data, estimated via 5-fold CV using the entire sample set ($n = 89$) with the best-tuned parameters ($r = 36, \lambda = 3611$). Only the top 30 gene expression features (rows) and CNV features (columns) are presented, selected based on the $\ell_2$-norm of their low-rank factors.

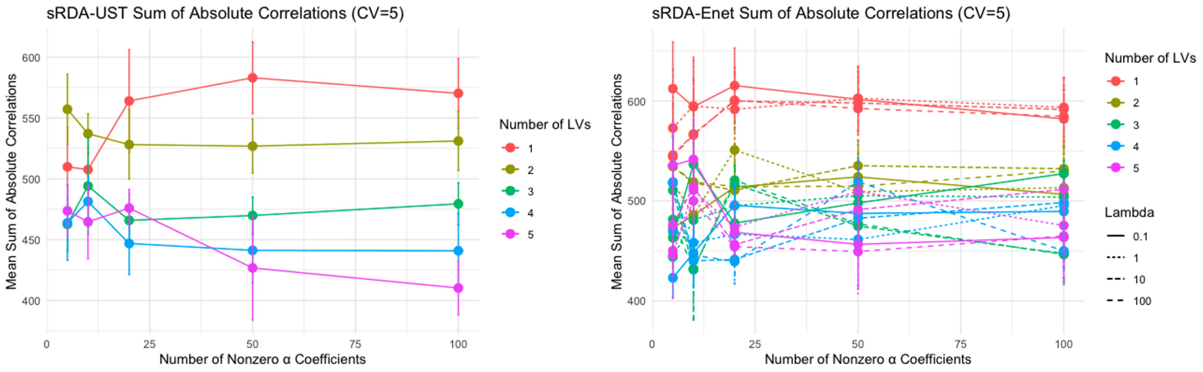


Fig. 4S: Parameter tuning of **sRDA** (Csala et al., 2017) on the whole-genome Breast Cancer dataset. The evaluation was based on five random splits ($n_{\text{train}} = 79, n_{\text{test}} = 10$). Within each training set, parameters were tuned via internal 5-fold CV to maximize the sum of absolute correlations, following the procedure by Csala et al. (2017). The mean sum of absolute correlations is plotted with error bars representing the standard error across CV folds. Both Univariate Soft Thresholding (UST) and Elastic Net penalties were evaluated with 1–5 latent variables, where the number of nonzero weights was selected from $\{5, 10, 20, 50, 100\}$ and the Elastic Net penalty from $\lambda \in \{0.1, 1, 10, 100\}$.

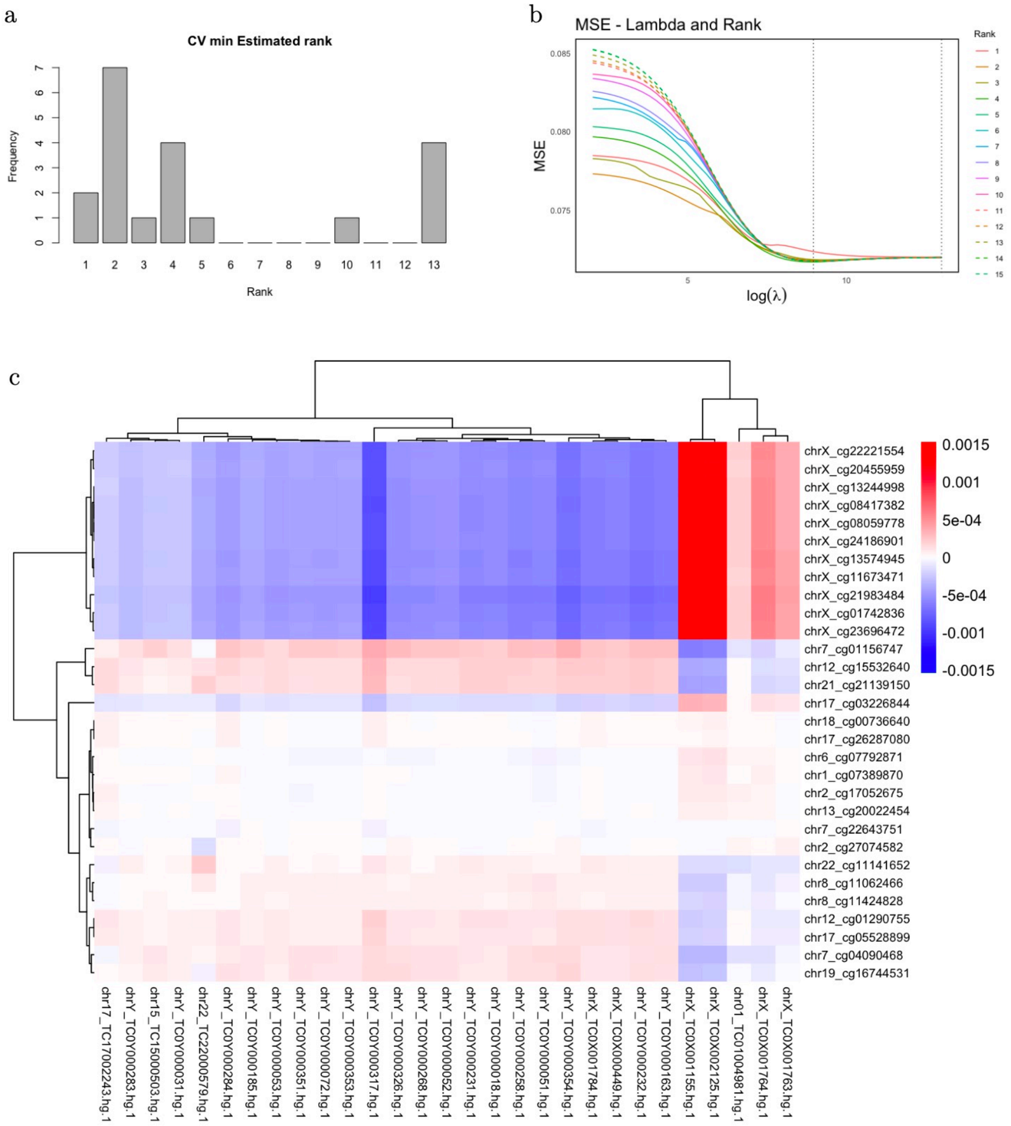


Fig. 5S: Application of **rrda** to the TCGA DNA methylation and gene expression dataset ($n = 20, p = 392,277, q = 67,528$). (a) Empirical distribution of the optimally selected rank $\hat{r}$ across the LOOCV procedure (20 iterations with $n_{\text{train}} = 19, n_{\text{test}} = 1$), where parameters were tuned via internal 5-fold CV in each iteration for performance evaluation. (b) Regularization path for parameter tuning via 5-fold CV using the entire sample set. (c) Heatmap of **rrda** coefficients as a low-dimensional representation. The coefficient matrix $\hat{B}$ was estimated using the entire sample set with the best-tuned parameters from the CV ($r = 4, \lambda = 9737$). Only the top 30 DNA methylation features (rows) and gene expression features (columns) are presented, selected based on the $\ell_2$-norm of their low-rank factors. For clarity, chromosome names have been prefixed to each feature label.

**Table 1S.** Computational resources. Resource B (server) was used for simulation Scenario 1 to evaluate extremely high-dimensional settings, where conventional methods often failed to converge. For details on parallel computing assessments, see Appendix G.

| | **Resource A** | **Resource B** |
|---|---|---|
| **CPU** | Apple M2 Max (12 cores) | AMD Ryzen Threadripper 3990X @2.9GHz (64 cores) |
| **Memory** | 64 GB | 256 GB |
| **R Version** | 4.4.1 (2024-06-14) | 4.4.1 (2024-06-14) |
| **OS** | macOS Sonoma 14.5 | Ubuntu 24.04 LTS |
| **System** | aarch64, darwin20 | x86 64, linux-gnu |
| **UI** | RStudio | RStudio |
| **Collate** | en US.UTF-8 | en US.UTF-8 |
| **CType** | en US.UTF-8 | en US.UTF-8 |
| **RStudio Version** | 2024.04.2+764 Chocolate Cosmos (desktop) | 2024.04.2+764 Chocolate Cosmos (server) |
| **Pandoc Version** | 3.1.11 | 3.1.11 |
| **Usage in experiments** | | |
| Scenario 1 (computational performance) | – | ✓(single core) |
| Scenario 2 (rank estimation) | ✓(single core) | – |
| Scenario 3 (prediction performance) | ✓(single core) | – |
| Lambda Sampling (LS) | ✓(single core) | – |
| Application (breast cancer, soybean, TCGA) | ✓(single core + parallel) | – |

**Table 2S.** Scenario 2: Estimated rank for classic information criteria with high noise; signal-to-noise ratio (s2n) = 0.1. For all combinations of parameters $(n, p, q, k)$. Mean values are presented with standard error.

| | | | | AIC | | BIC | | GIC | | BICP | | GCV | |
|---|---|---|---|---|---|---|---|---|---|---|---|---|---|
| Model | n | pq | k | mean | se | mean | se | mean | se | mean | se | mean | se |
| Model1 | 100 | 50 | 2 | 8.60 | 0.36 | 2.00 | 0.00 | 0.95 | 0.11 | 1.65 | 0.11 | 5.70 | 0.31 |
| Model1 | 200 | 50 | 2 | 2.00 | 0.00 | 2.00 | 0.00 | 0.20 | 0.09 | 1.80 | 0.09 | 2.00 | 0.00 |
| Model1 | 100 | 100 | 2 | 99.00 | 0.00 | 99.00 | 0.00 | 99.00 | 0.00 | 99.00 | 0.00 | 99.00 | 0.00 |
| Model1 | 200 | 100 | 2 | 2.00 | 0.00 | 2.00 | 0.00 | 0.50 | 0.11 | 1.95 | 0.05 | 2.00 | 0.00 |
| Model1 | 100 | 200 | 2 | 99.00 | 0.00 | 99.00 | 0.00 | 99.00 | 0.00 | 99.00 | 0.00 | 99.00 | 0.00 |
| Model1 | 200 | 200 | 2 | 200.00 | 0.00 | 200.00 | 0.00 | 200.00 | 0.00 | 200.00 | 0.00 | 199.00 | 0.00 |
| Model1 | 100 | 500 | 2 | 99.00 | 0.00 | 99.00 | 0.00 | 99.00 | 0.00 | 99.00 | 0.00 | 99.00 | 0.00 |
| Model1 | 200 | 500 | 2 | 199.20 | 0.09 | 200.00 | 0.00 | 200.00 | 0.00 | 200.00 | 0.00 | 199.00 | 0.00 |
| Model1 | 100 | 50 | 5 | 10.45 | 0.22 | 2.80 | 0.24 | 0.00 | 0.00 | 0.00 | 0.00 | 7.25 | 0.16 |
| Model1 | 200 | 50 | 5 | 5.00 | 0.00 | 1.60 | 0.13 | 0.00 | 0.00 | 0.00 | 0.00 | 5.00 | 0.00 |
| Model1 | 100 | 100 | 5 | 99.00 | 0.00 | 99.00 | 0.00 | 99.00 | 0.00 | 99.00 | 0.00 | 99.00 | 0.00 |
| Model1 | 200 | 100 | 5 | 5.00 | 0.00 | 1.80 | 0.27 | 0.00 | 0.00 | 0.00 | 0.00 | 5.00 | 0.00 |
| Model1 | 100 | 200 | 5 | 99.00 | 0.00 | 99.00 | 0.00 | 99.00 | 0.00 | 99.00 | 0.00 | 99.00 | 0.00 |
| Model1 | 200 | 200 | 5 | 200.00 | 0.00 | 200.00 | 0.00 | 200.00 | 0.00 | 200.00 | 0.00 | 199.00 | 0.00 |
| Model1 | 100 | 500 | 5 | 99.00 | 0.00 | 99.00 | 0.00 | 99.00 | 0.00 | 99.00 | 0.00 | 99.00 | 0.00 |
| Model1 | 200 | 500 | 5 | 199.95 | 0.05 | 200.00 | 0.00 | 200.00 | 0.00 | 200.00 | 0.00 | 199.00 | 0.00 |
| Model1 | 100 | 50 | 10 | 12.55 | 0.27 | 0.05 | 0.05 | 0.00 | 0.00 | 0.00 | 0.00 | 10.25 | 0.12 |
| Model1 | 200 | 50 | 10 | 9.80 | 0.09 | 0.00 | 0.00 | 0.00 | 0.00 | 0.00 | 0.00 | 9.40 | 0.13 |
| Model1 | 100 | 100 | 10 | 99.00 | 0.00 | 99.00 | 0.00 | 99.00 | 0.00 | 99.00 | 0.00 | 99.00 | 0.00 |
| Model1 | 200 | 100 | 10 | 10.00 | 0.00 | 0.00 | 0.00 | 0.00 | 0.00 | 0.00 | 0.00 | 10.00 | 0.00 |
| Model1 | 100 | 200 | 10 | 99.00 | 0.00 | 99.00 | 0.00 | 99.00 | 0.00 | 99.00 | 0.00 | 99.00 | 0.00 |
| Model1 | 200 | 200 | 10 | 200.00 | 0.00 | 200.00 | 0.00 | 200.00 | 0.00 | 200.00 | 0.00 | 199.00 | 0.00 |
| Model1 | 100 | 500 | 10 | 99.00 | 0.00 | 99.00 | 0.00 | 99.00 | 0.00 | 99.00 | 0.00 | 99.00 | 0.00 |
| Model1 | 200 | 500 | 10 | 200.00 | 0.00 | 200.00 | 0.00 | 200.00 | 0.00 | 200.00 | 0.00 | 199.00 | 0.00 |
| Model2 | 100 | 50 | 2 | 9.50 | 0.38 | 2.00 | 0.00 | 1.50 | 0.15 | 1.85 | 0.08 | 6.60 | 0.27 |
| Model2 | 200 | 50 | 2 | 2.00 | 0.00 | 2.00 | 0.00 | 0.75 | 0.14 | 1.75 | 0.10 | 2.00 | 0.00 |
| Model2 | 100 | 100 | 2 | 99.00 | 0.00 | 99.00 | 0.00 | 99.00 | 0.00 | 99.00 | 0.00 | 99.00 | 0.00 |
| Model2 | 200 | 100 | 2 | 2.00 | 0.00 | 2.00 | 0.00 | 0.50 | 0.15 | 1.95 | 0.05 | 2.00 | 0.00 |
| Model2 | 100 | 200 | 2 | 99.00 | 0.00 | 99.00 | 0.00 | 99.00 | 0.00 | 99.00 | 0.00 | 99.00 | 0.00 |
| Model2 | 200 | 200 | 2 | 200.00 | 0.00 | 200.00 | 0.00 | 200.00 | 0.00 | 200.00 | 0.00 | 199.00 | 0.00 |
| Model2 | 100 | 500 | 2 | 99.00 | 0.00 | 99.00 | 0.00 | 99.00 | 0.00 | 99.00 | 0.00 | 99.00 | 0.00 |
| Model2 | 200 | 500 | 2 | 199.55 | 0.11 | 200.00 | 0.00 | 200.00 | 0.00 | 200.00 | 0.00 | 199.00 | 0.00 |
| Model2 | 100 | 50 | 5 | 11.10 | 0.29 | 3.20 | 0.20 | 0.00 | 0.00 | 0.00 | 0.00 | 7.95 | 0.18 |
| Model2 | 200 | 50 | 5 | 5.00 | 0.00 | 2.70 | 0.18 | 0.00 | 0.00 | 0.00 | 0.00 | 5.00 | 0.00 |
| Model2 | 100 | 100 | 5 | 99.00 | 0.00 | 99.00 | 0.00 | 99.00 | 0.00 | 99.00 | 0.00 | 99.00 | 0.00 |
| Model2 | 200 | 100 | 5 | 5.00 | 0.00 | 3.25 | 0.28 | 0.00 | 0.00 | 0.00 | 0.00 | 5.00 | 0.00 |
| Model2 | 100 | 200 | 5 | 99.00 | 0.00 | 99.00 | 0.00 | 99.00 | 0.00 | 99.00 | 0.00 | 99.00 | 0.00 |
| Model2 | 200 | 200 | 5 | 200.00 | 0.00 | 200.00 | 0.00 | 200.00 | 0.00 | 200.00 | 0.00 | 199.00 | 0.00 |
| Model2 | 100 | 500 | 5 | 99.00 | 0.00 | 99.00 | 0.00 | 99.00 | 0.00 | 99.00 | 0.00 | 99.00 | 0.00 |
| Model2 | 200 | 500 | 5 | 200.00 | 0.00 | 200.00 | 0.00 | 200.00 | 0.00 | 200.00 | 0.00 | 199.00 | 0.00 |
| Model2 | 100 | 50 | 10 | 14.95 | 0.28 | 0.50 | 0.14 | 0.00 | 0.00 | 0.00 | 0.00 | 11.75 | 0.19 |
| Model2 | 200 | 50 | 10 | 9.85 | 0.08 | 0.15 | 0.08 | 0.00 | 0.00 | 0.00 | 0.00 | 9.60 | 0.11 |
| Model2 | 100 | 100 | 10 | 99.00 | 0.00 | 99.00 | 0.00 | 99.00 | 0.00 | 99.00 | 0.00 | 99.00 | 0.00 |
| Model2 | 200 | 100 | 10 | 10.00 | 0.00 | 0.00 | 0.00 | 0.00 | 0.00 | 0.00 | 0.00 | 10.00 | 0.00 |
| Model2 | 100 | 200 | 10 | 99.00 | 0.00 | 99.00 | 0.00 | 99.00 | 0.00 | 99.00 | 0.00 | 99.00 | 0.00 |
| Model2 | 200 | 200 | 10 | 200.00 | 0.00 | 200.00 | 0.00 | 200.00 | 0.00 | 200.00 | 0.00 | 199.00 | 0.00 |
| Model2 | 100 | 500 | 10 | 99.00 | 0.00 | 99.00 | 0.00 | 99.00 | 0.00 | 99.00 | 0.00 | 99.00 | 0.00 |
| Model2 | 200 | 500 | 10 | 200.00 | 0.00 | 200.00 | 0.00 | 200.00 | 0.00 | 200.00 | 0.00 | 199.00 | 0.00 |

**Table 3S.** Scenario 2: Estimated rank for classic information criteria with mid noise; signal-to-noise ratio (s2n) = 1. For all combinations of parameters $(n, p, q, k)$. Mean values are presented with standard error.

| | | | | AIC | | BIC | | GIC | | BICP | | GCV | |
|---|---|---|---|---|---|---|---|---|---|---|---|---|---|
| Model | n | pq | k | mean | se | mean | se | mean | se | mean | se | mean | se |
| Model1 | 100 | 50 | 2 | 8.60 | 0.36 | 2.00 | 0.00 | 0.95 | 0.11 | 1.65 | 0.11 | 5.70 | 0.31 |
| Model1 | 200 | 50 | 2 | 2.00 | 0.00 | 2.00 | 0.00 | 0.20 | 0.09 | 1.80 | 0.09 | 2.00 | 0.00 |
| Model1 | 100 | 100 | 2 | 99.00 | 0.00 | 99.00 | 0.00 | 99.00 | 0.00 | 99.00 | 0.00 | 99.00 | 0.00 |
| Model1 | 200 | 100 | 2 | 2.00 | 0.00 | 2.00 | 0.00 | 0.50 | 0.11 | 1.95 | 0.05 | 2.00 | 0.00 |
| Model1 | 100 | 200 | 2 | 99.00 | 0.00 | 99.00 | 0.00 | 99.00 | 0.00 | 99.00 | 0.00 | 99.00 | 0.00 |
| Model1 | 200 | 200 | 2 | 200.00 | 0.00 | 200.00 | 0.00 | 200.00 | 0.00 | 200.00 | 0.00 | 199.00 | 0.00 |
| Model1 | 100 | 500 | 2 | 99.00 | 0.00 | 99.00 | 0.00 | 99.00 | 0.00 | 99.00 | 0.00 | 99.00 | 0.00 |
| Model1 | 200 | 500 | 2 | 199.20 | 0.09 | 200.00 | 0.00 | 200.00 | 0.00 | 200.00 | 0.00 | 199.00 | 0.00 |
| Model1 | 100 | 50 | 5 | 10.45 | 0.22 | 2.80 | 0.24 | 0.00 | 0.00 | 0.00 | 0.00 | 7.25 | 0.16 |
| Model1 | 200 | 50 | 5 | 5.00 | 0.00 | 1.60 | 0.13 | 0.00 | 0.00 | 0.00 | 0.00 | 5.00 | 0.00 |
| Model1 | 100 | 100 | 5 | 99.00 | 0.00 | 99.00 | 0.00 | 99.00 | 0.00 | 99.00 | 0.00 | 99.00 | 0.00 |
| Model1 | 200 | 100 | 5 | 5.00 | 0.00 | 1.80 | 0.27 | 0.00 | 0.00 | 0.00 | 0.00 | 5.00 | 0.00 |
| Model1 | 100 | 200 | 5 | 99.00 | 0.00 | 99.00 | 0.00 | 99.00 | 0.00 | 99.00 | 0.00 | 99.00 | 0.00 |
| Model1 | 200 | 200 | 5 | 200.00 | 0.00 | 200.00 | 0.00 | 200.00 | 0.00 | 200.00 | 0.00 | 199.00 | 0.00 |
| Model1 | 100 | 500 | 5 | 99.00 | 0.00 | 99.00 | 0.00 | 99.00 | 0.00 | 99.00 | 0.00 | 99.00 | 0.00 |
| Model1 | 200 | 500 | 5 | 199.95 | 0.05 | 200.00 | 0.00 | 200.00 | 0.00 | 200.00 | 0.00 | 199.00 | 0.00 |
| Model1 | 100 | 50 | 10 | 12.55 | 0.27 | 0.05 | 0.05 | 0.00 | 0.00 | 0.00 | 0.00 | 10.25 | 0.12 |
| Model1 | 200 | 50 | 10 | 9.80 | 0.09 | 0.00 | 0.00 | 0.00 | 0.00 | 0.00 | 0.00 | 9.40 | 0.13 |
| Model1 | 100 | 100 | 10 | 99.00 | 0.00 | 99.00 | 0.00 | 99.00 | 0.00 | 99.00 | 0.00 | 99.00 | 0.00 |
| Model1 | 200 | 100 | 10 | 10.00 | 0.00 | 0.00 | 0.00 | 0.00 | 0.00 | 0.00 | 0.00 | 10.00 | 0.00 |
| Model1 | 100 | 200 | 10 | 99.00 | 0.00 | 99.00 | 0.00 | 99.00 | 0.00 | 99.00 | 0.00 | 99.00 | 0.00 |
| Model1 | 200 | 200 | 10 | 200.00 | 0.00 | 200.00 | 0.00 | 200.00 | 0.00 | 200.00 | 0.00 | 199.00 | 0.00 |
| Model1 | 100 | 500 | 10 | 99.00 | 0.00 | 99.00 | 0.00 | 99.00 | 0.00 | 99.00 | 0.00 | 99.00 | 0.00 |
| Model1 | 200 | 500 | 10 | 200.00 | 0.00 | 200.00 | 0.00 | 200.00 | 0.00 | 200.00 | 0.00 | 199.00 | 0.00 |
| Model2 | 100 | 50 | 2 | 9.50 | 0.38 | 2.00 | 0.00 | 1.50 | 0.15 | 1.85 | 0.08 | 6.60 | 0.27 |
| Model2 | 200 | 50 | 2 | 2.00 | 0.00 | 2.00 | 0.00 | 0.75 | 0.14 | 1.75 | 0.10 | 2.00 | 0.00 |
| Model2 | 100 | 100 | 2 | 99.00 | 0.00 | 99.00 | 0.00 | 99.00 | 0.00 | 99.00 | 0.00 | 99.00 | 0.00 |
| Model2 | 200 | 100 | 2 | 2.00 | 0.00 | 2.00 | 0.00 | 0.50 | 0.15 | 1.95 | 0.05 | 2.00 | 0.00 |
| Model2 | 100 | 200 | 2 | 99.00 | 0.00 | 99.00 | 0.00 | 99.00 | 0.00 | 99.00 | 0.00 | 99.00 | 0.00 |
| Model2 | 200 | 200 | 2 | 200.00 | 0.00 | 200.00 | 0.00 | 200.00 | 0.00 | 200.00 | 0.00 | 199.00 | 0.00 |
| Model2 | 100 | 500 | 2 | 99.00 | 0.00 | 99.00 | 0.00 | 99.00 | 0.00 | 99.00 | 0.00 | 99.00 | 0.00 |
| Model2 | 200 | 500 | 2 | 199.55 | 0.11 | 200.00 | 0.00 | 200.00 | 0.00 | 200.00 | 0.00 | 199.00 | 0.00 |
| Model2 | 100 | 50 | 5 | 11.10 | 0.29 | 3.20 | 0.20 | 0.00 | 0.00 | 0.00 | 0.00 | 7.95 | 0.18 |
| Model2 | 200 | 50 | 5 | 5.00 | 0.00 | 2.70 | 0.18 | 0.00 | 0.00 | 0.00 | 0.00 | 5.00 | 0.00 |
| Model2 | 100 | 100 | 5 | 99.00 | 0.00 | 99.00 | 0.00 | 99.00 | 0.00 | 99.00 | 0.00 | 99.00 | 0.00 |
| Model2 | 200 | 100 | 5 | 5.00 | 0.00 | 3.25 | 0.28 | 0.00 | 0.00 | 0.00 | 0.00 | 5.00 | 0.00 |
| Model2 | 100 | 200 | 5 | 99.00 | 0.00 | 99.00 | 0.00 | 99.00 | 0.00 | 99.00 | 0.00 | 99.00 | 0.00 |
| Model2 | 200 | 200 | 5 | 200.00 | 0.00 | 200.00 | 0.00 | 200.00 | 0.00 | 200.00 | 0.00 | 199.00 | 0.00 |
| Model2 | 100 | 500 | 5 | 99.00 | 0.00 | 99.00 | 0.00 | 99.00 | 0.00 | 99.00 | 0.00 | 99.00 | 0.00 |
| Model2 | 200 | 500 | 5 | 200.00 | 0.00 | 200.00 | 0.00 | 200.00 | 0.00 | 200.00 | 0.00 | 199.00 | 0.00 |
| Model2 | 100 | 50 | 10 | 14.95 | 0.28 | 0.50 | 0.14 | 0.00 | 0.00 | 0.00 | 0.00 | 11.75 | 0.19 |
| Model2 | 200 | 50 | 10 | 9.85 | 0.08 | 0.15 | 0.08 | 0.00 | 0.00 | 0.00 | 0.00 | 9.60 | 0.11 |
| Model2 | 100 | 100 | 10 | 99.00 | 0.00 | 99.00 | 0.00 | 99.00 | 0.00 | 99.00 | 0.00 | 99.00 | 0.00 |
| Model2 | 200 | 100 | 10 | 10.00 | 0.00 | 0.00 | 0.00 | 0.00 | 0.00 | 0.00 | 0.00 | 10.00 | 0.00 |
| Model2 | 100 | 200 | 10 | 99.00 | 0.00 | 99.00 | 0.00 | 99.00 | 0.00 | 99.00 | 0.00 | 99.00 | 0.00 |
| Model2 | 200 | 200 | 10 | 200.00 | 0.00 | 200.00 | 0.00 | 200.00 | 0.00 | 200.00 | 0.00 | 199.00 | 0.00 |
| Model2 | 100 | 500 | 10 | 99.00 | 0.00 | 99.00 | 0.00 | 99.00 | 0.00 | 99.00 | 0.00 | 99.00 | 0.00 |
| Model2 | 200 | 500 | 10 | 200.00 | 0.00 | 200.00 | 0.00 | 200.00 | 0.00 | 200.00 | 0.00 | 199.00 | 0.00 |

**Table 4S.** Scenario 2: Estimated rank for classic information criteria with low noise; signal-to-noise ratio (s2n) = 10. For all combinations of parameters $(n, p, q, k)$. Mean values are presented with standard error.

| | | | | AIC | | BIC | | GIC | | BICP | | GCV | |
|---|---|---|---|---|---|---|---|---|---|---|---|---|---|
| Model | n | pq | k | mean | se | mean | se | mean | se | mean | se | mean | se |
| Model1 | 100 | 50 | 2 | 20.40 | 0.30 | 2.00 | 0.00 | 2.00 | 0.00 | 2.00 | 0.00 | 17.00 | 0.21 |
| Model1 | 200 | 50 | 2 | 2.35 | 0.11 | 2.00 | 0.00 | 2.00 | 0.00 | 2.00 | 0.00 | 2.30 | 0.11 |
| Model1 | 100 | 100 | 2 | 99.00 | 0.00 | 99.00 | 0.00 | 99.00 | 0.00 | 99.00 | 0.00 | 99.00 | 0.00 |
| Model1 | 200 | 100 | 2 | 3.55 | 0.21 | 2.00 | 0.00 | 2.00 | 0.00 | 2.00 | 0.00 | 2.80 | 0.12 |
| Model1 | 100 | 200 | 2 | 99.00 | 0.00 | 99.00 | 0.00 | 99.00 | 0.00 | 99.00 | 0.00 | 99.00 | 0.00 |
| Model1 | 200 | 200 | 2 | 200.00 | 0.00 | 200.00 | 0.00 | 200.00 | 0.00 | 200.00 | 0.00 | 200.00 | 0.00 |
| Model1 | 100 | 500 | 2 | 99.00 | 0.00 | 99.00 | 0.00 | 99.00 | 0.00 | 99.00 | 0.00 | 99.00 | 0.00 |
| Model1 | 200 | 500 | 2 | 199.00 | 0.00 | 199.00 | 0.00 | 199.10 | 0.07 | 199.05 | 0.05 | 199.00 | 0.00 |
| Model1 | 100 | 50 | 5 | 20.70 | 0.25 | 5.00 | 0.00 | 5.00 | 0.00 | 5.00 | 0.00 | 17.60 | 0.23 |
| Model1 | 200 | 50 | 5 | 5.05 | 0.05 | 5.00 | 0.00 | 5.00 | 0.00 | 5.00 | 0.00 | 5.00 | 0.00 |
| Model1 | 100 | 100 | 5 | 99.00 | 0.00 | 99.00 | 0.00 | 99.00 | 0.00 | 99.00 | 0.00 | 99.00 | 0.00 |
| Model1 | 200 | 100 | 5 | 7.05 | 0.18 | 5.00 | 0.00 | 5.00 | 0.00 | 5.00 | 0.00 | 5.50 | 0.17 |
| Model1 | 100 | 200 | 5 | 99.00 | 0.00 | 99.00 | 0.00 | 99.00 | 0.00 | 99.00 | 0.00 | 99.00 | 0.00 |
| Model1 | 200 | 200 | 5 | 200.00 | 0.00 | 200.00 | 0.00 | 200.00 | 0.00 | 200.00 | 0.00 | 200.00 | 0.00 |
| Model1 | 100 | 500 | 5 | 99.00 | 0.00 | 99.00 | 0.00 | 99.00 | 0.00 | 99.00 | 0.00 | 99.00 | 0.00 |
| Model1 | 200 | 500 | 5 | 199.05 | 0.05 | 199.70 | 0.11 | 199.95 | 0.05 | 199.95 | 0.05 | 199.30 | 0.11 |
| Model1 | 100 | 50 | 10 | 21.05 | 0.21 | 10.00 | 0.00 | 9.85 | 0.08 | 9.95 | 0.05 | 17.95 | 0.18 |
| Model1 | 200 | 50 | 10 | 10.00 | 0.00 | 10.00 | 0.00 | 9.85 | 0.08 | 10.00 | 0.00 | 10.00 | 0.00 |
| Model1 | 100 | 100 | 10 | 99.00 | 0.00 | 99.00 | 0.00 | 99.00 | 0.00 | 99.00 | 0.00 | 99.00 | 0.00 |
| Model1 | 200 | 100 | 10 | 11.40 | 0.23 | 10.00 | 0.00 | 10.00 | 0.00 | 10.00 | 0.00 | 10.05 | 0.05 |
| Model1 | 100 | 200 | 10 | 99.00 | 0.00 | 99.00 | 0.00 | 99.00 | 0.00 | 99.00 | 0.00 | 99.00 | 0.00 |
| Model1 | 200 | 200 | 10 | 200.00 | 0.00 | 200.00 | 0.00 | 200.00 | 0.00 | 200.00 | 0.00 | 200.00 | 0.00 |
| Model1 | 100 | 500 | 10 | 99.00 | 0.00 | 99.00 | 0.00 | 99.00 | 0.00 | 99.00 | 0.00 | 99.00 | 0.00 |
| Model1 | 200 | 500 | 10 | 199.10 | 0.07 | 200.00 | 0.00 | 200.00 | 0.00 | 200.00 | 0.00 | 199.90 | 0.07 |
| Model2 | 100 | 50 | 2 | 20.80 | 0.20 | 2.00 | 0.00 | 2.00 | 0.00 | 2.00 | 0.00 | 17.50 | 0.25 |
| Model2 | 200 | 50 | 2 | 2.75 | 0.18 | 2.00 | 0.00 | 2.00 | 0.00 | 2.00 | 0.00 | 2.55 | 0.14 |
| Model2 | 100 | 100 | 2 | 99.00 | 0.00 | 99.00 | 0.00 | 99.00 | 0.00 | 99.00 | 0.00 | 99.00 | 0.00 |
| Model2 | 200 | 100 | 2 | 4.35 | 0.36 | 2.00 | 0.00 | 2.00 | 0.00 | 2.00 | 0.00 | 3.00 | 0.18 |
| Model2 | 100 | 200 | 2 | 99.00 | 0.00 | 99.00 | 0.00 | 99.00 | 0.00 | 99.00 | 0.00 | 99.00 | 0.00 |
| Model2 | 200 | 200 | 2 | 199.75 | 0.10 | 199.95 | 0.05 | 200.00 | 0.00 | 200.00 | 0.00 | 199.90 | 0.07 |
| Model2 | 100 | 500 | 2 | 99.00 | 0.00 | 99.00 | 0.00 | 99.00 | 0.00 | 99.00 | 0.00 | 99.00 | 0.00 |
| Model2 | 200 | 500 | 2 | 199.00 | 0.00 | 199.80 | 0.09 | 200.00 | 0.00 | 200.00 | 0.00 | 199.25 | 0.10 |
| Model2 | 100 | 50 | 5 | 22.75 | 0.24 | 5.00 | 0.00 | 5.00 | 0.00 | 5.00 | 0.00 | 19.05 | 0.28 |
| Model2 | 200 | 50 | 5 | 5.40 | 0.11 | 5.00 | 0.00 | 5.00 | 0.00 | 5.00 | 0.00 | 5.15 | 0.08 |
| Model2 | 100 | 100 | 5 | 99.00 | 0.00 | 99.00 | 0.00 | 99.00 | 0.00 | 99.00 | 0.00 | 99.00 | 0.00 |
| Model2 | 200 | 100 | 5 | 8.20 | 0.28 | 5.00 | 0.00 | 5.00 | 0.00 | 5.00 | 0.00 | 5.95 | 0.15 |
| Model2 | 100 | 200 | 5 | 99.00 | 0.00 | 99.00 | 0.00 | 99.00 | 0.00 | 99.00 | 0.00 | 99.00 | 0.00 |
| Model2 | 200 | 200 | 5 | 199.85 | 0.08 | 200.00 | 0.00 | 200.00 | 0.00 | 200.00 | 0.00 | 199.95 | 0.05 |
| Model2 | 100 | 500 | 5 | 99.00 | 0.00 | 99.00 | 0.00 | 99.00 | 0.00 | 99.00 | 0.00 | 99.00 | 0.00 |
| Model2 | 200 | 500 | 5 | 199.10 | 0.07 | 200.00 | 0.00 | 200.00 | 0.00 | 200.00 | 0.00 | 200.00 | 0.00 |
| Model2 | 100 | 50 | 10 | 24.20 | 0.20 | 10.00 | 0.00 | 9.95 | 0.05 | 10.00 | 0.00 | 20.95 | 0.18 |
| Model2 | 200 | 50 | 10 | 10.20 | 0.09 | 10.00 | 0.00 | 9.75 | 0.10 | 9.90 | 0.07 | 10.00 | 0.00 |
| Model2 | 100 | 100 | 10 | 99.00 | 0.00 | 99.00 | 0.00 | 99.00 | 0.00 | 99.00 | 0.00 | 99.00 | 0.00 |
| Model2 | 200 | 100 | 10 | 13.75 | 0.32 | 10.00 | 0.00 | 10.00 | 0.00 | 10.00 | 0.00 | 10.45 | 0.11 |
| Model2 | 100 | 200 | 10 | 99.00 | 0.00 | 99.00 | 0.00 | 99.00 | 0.00 | 99.00 | 0.00 | 99.00 | 0.00 |
| Model2 | 200 | 200 | 10 | 199.95 | 0.05 | 200.00 | 0.00 | 200.00 | 0.00 | 200.00 | 0.00 | 200.00 | 0.00 |
| Model2 | 100 | 500 | 10 | 99.00 | 0.00 | 99.00 | 0.00 | 99.00 | 0.00 | 99.00 | 0.00 | 99.00 | 0.00 |
| Model2 | 200 | 500 | 10 | 199.50 | 0.11 | 200.00 | 0.00 | 200.00 | 0.00 | 200.00 | 0.00 | 200.00 | 0.00 |

**Table 5S.** Scenario 2: Estimated rank for information criterion-based ANN and RDA-based CV with high noise; signal-to-noise ratio (s2n) = 0.1. For all combinations of parameters $(n, p, q, k)$. Mean values are presented with standard error.

| | | | | ANN | | | | | | | | | | | | RDA | | | |
|---|---|---|---|---|---|---|---|---|---|---|---|---|---|---|---|---|---|---|---|
| | | | | AIC | | BIC | | GIC | | BICP | | GCV | | StARS | | c-rda.cv | | rrda.cv | |
| Model | n | pq | k | mean | se | mean | se | mean | se | mean | se | mean | se | mean | se | mean | se | mean | se |
| Model 1 | 100 | 50 | 2 | 2.80 | 0.25 | 0.00 | 0.00 | 0.00 | 0.00 | 0.00 | 0.00 | 2.60 | 0.15 | 0.20 | 0.12 | 1.00 | 0.00 | 2.15 | 0.11 |
| Model 1 | 200 | 50 | 2 | 2.45 | 0.20 | 0.00 | 0.00 | 0.00 | 0.00 | 0.00 | 0.00 | 2.30 | 0.16 | 1.80 | 0.14 | 1.75 | 0.10 | 2.05 | 0.05 |
| Model 1 | 100 | 100 | 2 | 99.00 | 0.00 | 99.00 | 0.00 | 0.00 | 0.00 | 14.85 | 8.11 | 17.55 | 7.85 | 99.00 | 0.00 | 1.00 | 0.00 | 2.00 | 0.00 |
| Model 1 | 200 | 100 | 2 | 2.40 | 0.17 | 0.10 | 0.07 | 0.00 | 0.00 | 0.00 | 0.00 | 2.35 | 0.15 | 1.85 | 0.11 | 1.10 | 0.07 | 2.00 | 0.00 |
| Model 1 | 100 | 200 | 2 | 99.00 | 0.00 | 0.00 | 0.00 | 0.00 | 0.00 | 0.00 | 0.00 | 2.40 | 0.18 | 99.00 | 0.00 | 2.00 | 0.00 | 2.00 | 0.00 |
| Model 1 | 200 | 200 | 2 | 199.00 | 0.00 | 199.00 | 0.00 | 0.00 | 0.00 | 19.90 | 13.70 | 22.30 | 13.51 | 199.00 | 0.00 | 1.40 | 0.11 | 2.00 | 0.00 |
| Model 1 | 100 | 500 | 2 | 99.00 | 0.00 | 0.00 | 0.00 | 0.00 | 0.00 | 0.00 | 0.00 | 2.25 | 0.12 | 99.00 | 0.00 | 2.00 | 0.00 | 2.00 | 0.00 |
| Model 1 | 200 | 500 | 2 | 199.00 | 0.00 | 1.85 | 0.08 | 0.00 | 0.00 | 0.00 | 0.00 | 2.45 | 0.18 | 199.00 | 0.00 | 2.00 | 0.00 | 2.00 | 0.00 |
| Model 1 | 100 | 50 | 5 | 4.20 | 0.30 | 0.00 | 0.00 | 0.00 | 0.00 | 0.00 | 0.00 | 3.60 | 0.26 | 0.00 | 0.00 | 1.00 | 0.00 | 5.60 | 0.89 |
| Model 1 | 200 | 50 | 5 | 4.40 | 0.23 | 0.00 | 0.00 | 0.00 | 0.00 | 0.00 | 0.00 | 4.30 | 0.22 | 0.00 | 0.00 | 1.00 | 0.00 | 5.75 | 0.40 |
| Model 1 | 100 | 100 | 5 | 99.00 | 0.00 | 99.00 | 0.00 | 4.95 | 4.95 | 9.90 | 6.81 | 14.65 | 6.45 | 99.00 | 0.00 | 1.00 | 0.00 | 5.25 | 0.30 |
| Model 1 | 200 | 100 | 5 | 7.20 | 0.31 | 0.00 | 0.00 | 0.00 | 0.00 | 0.00 | 0.00 | 6.35 | 0.21 | 0.00 | 0.00 | 1.00 | 0.00 | 5.00 | 0.00 |
| Model 1 | 100 | 200 | 5 | 99.00 | 0.00 | 0.00 | 0.00 | 0.00 | 0.00 | 0.00 | 0.00 | 6.95 | 0.25 | 99.00 | 0.00 | 1.25 | 0.10 | 5.15 | 0.08 |
| Model 1 | 200 | 200 | 5 | 199.00 | 0.00 | 199.00 | 0.00 | 0.00 | 0.00 | 0.00 | 0.00 | 7.30 | 0.31 | 199.00 | 0.00 | 1.00 | 0.00 | 5.00 | 0.00 |
| Model 1 | 100 | 500 | 5 | 99.00 | 0.00 | 0.00 | 0.00 | 0.00 | 0.00 | 0.00 | 0.00 | 7.20 | 0.28 | 99.00 | 0.00 | 5.00 | 0.00 | 5.00 | 0.00 |
| Model 1 | 200 | 500 | 5 | 199.00 | 0.00 | 0.00 | 0.00 | 0.00 | 0.00 | 0.00 | 0.00 | 6.60 | 0.28 | 199.00 | 0.00 | 5.00 | 0.00 | 5.00 | 0.00 |
| Model 1 | 100 | 50 | 10 | 1.70 | 0.32 | 0.00 | 0.00 | 0.00 | 0.00 | 0.00 | 0.00 | 1.45 | 0.26 | 0.00 | 0.00 | 1.00 | 0.00 | 5.60 | 0.86 |
| Model 1 | 200 | 50 | 10 | 3.85 | 0.36 | 0.00 | 0.00 | 0.00 | 0.00 | 0.00 | 0.00 | 3.45 | 0.32 | 0.00 | 0.00 | 1.00 | 0.00 | 7.20 | 1.10 |
| Model 1 | 100 | 100 | 10 | 99.00 | 0.00 | 99.00 | 0.00 | 0.00 | 0.00 | 9.90 | 6.81 | 13.60 | 6.54 | 99.00 | 0.00 | 1.00 | 0.00 | 8.30 | 0.91 |
| Model 1 | 200 | 100 | 10 | 7.85 | 0.35 | 0.00 | 0.00 | 0.00 | 0.00 | 0.00 | 0.00 | 7.15 | 0.28 | 0.00 | 0.00 | 1.00 | 0.00 | 10.55 | 0.69 |
| Model 1 | 100 | 200 | 10 | 99.00 | 0.00 | 0.00 | 0.00 | 0.00 | 0.00 | 0.00 | 0.00 | 7.45 | 0.29 | 99.00 | 0.00 | 1.00 | 0.00 | 9.45 | 0.59 |
| Model 1 | 200 | 200 | 10 | 199.00 | 0.00 | 199.00 | 0.00 | 0.00 | 0.00 | 9.95 | 9.95 | 21.15 | 9.37 | 199.00 | 0.00 | 1.00 | 0.00 | 10.10 | 0.30 |
| Model 1 | 100 | 500 | 10 | 99.00 | 0.00 | 0.00 | 0.00 | 0.00 | 0.00 | 0.00 | 0.00 | 11.15 | 0.24 | 99.00 | 0.00 | 4.00 | 0.38 | 9.60 | 0.15 |
| Model 1 | 200 | 500 | 10 | 199.00 | 0.00 | 0.00 | 0.00 | 0.00 | 0.00 | 0.00 | 0.00 | 14.20 | 0.26 | 199.00 | 0.00 | 5.10 | 0.49 | 10.00 | 0.00 |
| Model 2 | 100 | 50 | 2 | 2.80 | 0.24 | 0.00 | 0.00 | 0.00 | 0.00 | 0.00 | 0.00 | 2.55 | 0.18 | 0.70 | 0.21 | 1.10 | 0.07 | 1.95 | 0.05 |
| Model 2 | 200 | 50 | 2 | 2.65 | 0.25 | 0.50 | 0.11 | 0.00 | 0.00 | 0.00 | 0.00 | 2.50 | 0.22 | 1.85 | 0.11 | 2.00 | 0.00 | 2.00 | 0.00 |
| Model 2 | 100 | 100 | 2 | 99.00 | 0.00 | 99.00 | 0.00 | 0.00 | 0.00 | 4.95 | 4.95 | 7.80 | 4.80 | 99.00 | 0.00 | 1.05 | 0.05 | 2.00 | 0.00 |
| Model 2 | 200 | 100 | 2 | 2.85 | 0.24 | 0.40 | 0.11 | 0.00 | 0.00 | 0.00 | 0.00 | 2.45 | 0.17 | 1.90 | 0.10 | 2.00 | 0.00 | 2.00 | 0.00 |
| Model 2 | 100 | 200 | 2 | 99.00 | 0.00 | 0.00 | 0.00 | 0.00 | 0.00 | 0.00 | 0.00 | 2.35 | 0.15 | 99.00 | 0.00 | 1.60 | 0.11 | 1.95 | 0.05 |
| Model 2 | 200 | 200 | 2 | 199.00 | 0.00 | 199.00 | 0.00 | 0.00 | 0.00 | 9.95 | 9.95 | 12.35 | 9.83 | 199.00 | 0.00 | 1.70 | 0.11 | 2.00 | 0.00 |
| Model 2 | 100 | 500 | 2 | 99.00 | 0.00 | 0.00 | 0.00 | 0.00 | 0.00 | 0.00 | 0.00 | 2.35 | 0.15 | 99.00 | 0.00 | 1.65 | 0.11 | 2.00 | 0.00 |
| Model 2 | 200 | 500 | 2 | 199.00 | 0.00 | 1.70 | 0.11 | 0.00 | 0.00 | 0.00 | 0.00 | 2.10 | 0.07 | 199.00 | 0.00 | 1.90 | 0.07 | 2.00 | 0.00 |
| Model 2 | 100 | 50 | 5 | 4.80 | 0.28 | 0.00 | 0.00 | 0.00 | 0.00 | 0.00 | 0.00 | 4.25 | 0.23 | 0.00 | 0.00 | 1.00 | 0.00 | 4.90 | 0.67 |
| Model 2 | 200 | 50 | 5 | 5.65 | 0.22 | 0.00 | 0.00 | 0.00 | 0.00 | 0.00 | 0.00 | 5.50 | 0.20 | 0.15 | 0.08 | 1.85 | 0.18 | 4.75 | 0.14 |
| Model 2 | 100 | 100 | 5 | 99.00 | 0.00 | 99.00 | 0.00 | 4.95 | 4.95 | 9.90 | 6.81 | 14.35 | 6.48 | 99.00 | 0.00 | 1.00 | 0.00 | 4.05 | 0.32 |
| Model 2 | 200 | 100 | 5 | 6.90 | 0.24 | 0.00 | 0.00 | 0.00 | 0.00 | 0.00 | 0.00 | 6.55 | 0.23 | 0.50 | 0.17 | 1.05 | 0.05 | 4.55 | 0.11 |
| Model 2 | 100 | 200 | 5 | 99.00 | 0.00 | 0.00 | 0.00 | 0.00 | 0.00 | 0.00 | 0.00 | 6.95 | 0.26 | 99.00 | 0.00 | 1.25 | 0.12 | 4.25 | 0.29 |
| Model 2 | 200 | 200 | 5 | 199.00 | 0.00 | 199.00 | 0.00 | 0.00 | 0.00 | 9.95 | 9.95 | 17.25 | 9.58 | 199.00 | 0.00 | 1.00 | 0.00 | 4.90 | 0.07 |
| Model 2 | 100 | 500 | 5 | 99.00 | 0.00 | 0.00 | 0.00 | 0.00 | 0.00 | 0.00 | 0.00 | 7.00 | 0.26 | 99.00 | 0.00 | 2.00 | 0.27 | 4.35 | 0.24 |
| Model 2 | 200 | 500 | 5 | 199.00 | 0.00 | 0.00 | 0.00 | 0.00 | 0.00 | 0.00 | 0.00 | 6.55 | 0.22 | 199.00 | 0.00 | 4.10 | 0.20 | 4.95 | 0.05 |
| Model 2 | 100 | 50 | 10 | 4.15 | 0.59 | 0.00 | 0.00 | 0.00 | 0.00 | 0.00 | 0.00 | 3.35 | 0.44 | 0.00 | 0.00 | 1.00 | 0.00 | 8.70 | 1.05 |
| Model 2 | 200 | 50 | 10 | 8.05 | 0.35 | 0.00 | 0.00 | 0.00 | 0.00 | 0.00 | 0.00 | 7.25 | 0.31 | 0.00 | 0.00 | 1.25 | 0.10 | 7.60 | 0.69 |
| Model 2 | 100 | 100 | 10 | 99.00 | 0.00 | 99.00 | 0.00 | 4.95 | 4.95 | 14.85 | 8.11 | 18.15 | 7.80 | 99.00 | 0.00 | 1.00 | 0.00 | 8.70 | 0.78 |
| Model 2 | 200 | 100 | 10 | 10.35 | 0.39 | 0.00 | 0.00 | 0.00 | 0.00 | 0.00 | 0.00 | 9.20 | 0.25 | 0.00 | 0.00 | 1.00 | 0.00 | 8.70 | 0.48 |
| Model 2 | 100 | 200 | 10 | 99.00 | 0.00 | 0.00 | 0.00 | 0.00 | 0.00 | 0.00 | 0.00 | 7.00 | 0.35 | 99.00 | 0.00 | 1.00 | 0.00 | 8.00 | 0.90 |
| Model 2 | 200 | 200 | 10 | 199.00 | 0.00 | 199.00 | 0.00 | 0.00 | 0.00 | 19.90 | 13.70 | 30.25 | 12.91 | 199.00 | 0.00 | 1.00 | 0.00 | 7.85 | 0.48 |
| Model 2 | 100 | 500 | 10 | 99.00 | 0.00 | 0.00 | 0.00 | 0.00 | 0.00 | 0.00 | 0.00 | 11.20 | 0.28 | 99.00 | 0.00 | 1.30 | 0.16 | 6.95 | 0.73 |
| Model 2 | 200 | 500 | 10 | 199.00 | 0.00 | 0.00 | 0.00 | 0.00 | 0.00 | 0.00 | 0.00 | 13.50 | 0.30 | 199.00 | 0.00 | 1.75 | 0.23 | 8.85 | 0.32 |

**Table 6S.** Scenario 2: Estimated rank for information criterion-based ANN and RDA-based CV with mid noise; signal-to-noise ratio (s2n) = 1. For all combinations of parameters $(n, p, q, k)$. Mean values are presented with standard error.

| | | | | **ANN** | | | | | | | | | | | | **RDA** | | | |
|---|---|---|---|---|---|---|---|---|---|---|---|---|---|---|---|---|---|---|---|
| | | | | AIC | | BIC | | GIC | | BICP | | GCV | | StARS | | c-rda.cv | | rrda.cv | |
| Model | n | pq | k | mean | se | mean | se | mean | se | mean | se | mean | se | mean | se | mean | se | mean | se |
| Model 1 | 100 | 50 | 2 | 2.20 | 0.16 | 2.00 | 0.00 | 1.95 | 0.05 | 2.00 | 0.00 | 2.10 | 0.10 | 1.65 | 0.15 | 2.00 | 0.00 | 2.05 | 0.05 |
| Model 1 | 200 | 50 | 2 | 2.30 | 0.16 | 2.00 | 0.00 | 2.00 | 0.00 | 2.00 | 0.00 | 2.15 | 0.08 | 1.95 | 0.05 | 2.00 | 0.00 | 2.00 | 0.00 |
| Model 1 | 100 | 100 | 2 | 99.00 | 0.00 | 99.00 | 0.00 | 1.95 | 0.05 | 2.00 | 0.00 | 2.25 | 0.14 | 99.00 | 0.00 | 2.00 | 0.00 | 2.00 | 0.00 |
| Model 1 | 200 | 100 | 2 | 2.40 | 0.23 | 2.00 | 0.00 | 2.00 | 0.00 | 2.00 | 0.00 | 2.40 | 0.23 | 1.95 | 0.05 | 2.00 | 0.00 | 2.00 | 0.00 |
| Model 1 | 100 | 200 | 2 | 99.00 | 0.00 | 2.00 | 0.00 | 2.00 | 0.00 | 2.00 | 0.00 | 2.30 | 0.15 | 99.00 | 0.00 | 2.00 | 0.00 | 2.00 | 0.00 |
| Model 1 | 200 | 200 | 2 | 199.00 | 0.00 | 199.00 | 0.00 | 2.00 | 0.00 | 2.00 | 0.00 | 2.00 | 0.00 | 199.00 | 0.00 | 2.00 | 0.00 | 2.00 | 0.00 |
| Model 1 | 100 | 500 | 2 | 99.00 | 0.00 | 2.00 | 0.00 | 2.00 | 0.00 | 2.00 | 0.00 | 2.30 | 0.18 | 99.00 | 0.00 | 2.00 | 0.00 | 2.00 | 0.00 |
| Model 1 | 200 | 500 | 2 | 199.00 | 0.00 | 2.00 | 0.00 | 2.00 | 0.00 | 2.00 | 0.00 | 2.05 | 0.05 | 199.00 | 0.00 | 2.00 | 0.00 | 2.00 | 0.00 |
| Model 1 | 100 | 50 | 5 | 5.20 | 0.16 | 3.85 | 0.18 | 0.00 | 0.00 | 0.00 | 0.00 | 5.00 | 0.00 | 4.55 | 0.31 | 4.70 | 0.11 | 5.15 | 0.11 |
| Model 1 | 200 | 50 | 5 | 5.00 | 0.00 | 5.00 | 0.00 | 3.30 | 0.16 | 3.80 | 0.19 | 5.00 | 0.00 | 4.65 | 0.24 | 5.00 | 0.00 | 5.00 | 0.00 |
| Model 1 | 100 | 100 | 5 | 99.00 | 0.00 | 99.00 | 0.00 | 0.00 | 0.00 | 4.95 | 4.95 | 9.90 | 4.69 | 99.00 | 0.00 | 4.95 | 0.05 | 5.20 | 0.20 |
| Model 1 | 200 | 100 | 5 | 5.25 | 0.18 | 5.00 | 0.00 | 1.10 | 0.18 | 3.15 | 0.22 | 5.10 | 0.10 | 4.55 | 0.31 | 5.00 | 0.00 | 5.00 | 0.00 |
| Model 1 | 100 | 200 | 5 | 99.00 | 0.00 | 4.95 | 0.05 | 0.00 | 0.00 | 0.00 | 0.00 | 5.05 | 0.05 | 99.00 | 0.00 | 5.00 | 0.00 | 5.00 | 0.00 |
| Model 1 | 200 | 200 | 5 | 199.00 | 0.00 | 199.00 | 0.00 | 0.20 | 0.09 | 1.70 | 0.22 | 5.00 | 0.00 | 199.00 | 0.00 | 5.00 | 0.00 | 5.00 | 0.00 |
| Model 1 | 100 | 500 | 5 | 99.00 | 0.00 | 5.00 | 0.00 | 0.00 | 0.00 | 0.00 | 0.00 | 5.00 | 0.00 | 99.00 | 0.00 | 5.00 | 0.00 | 5.00 | 0.00 |
| Model 1 | 200 | 500 | 5 | 199.00 | 0.00 | 5.00 | 0.00 | 1.45 | 0.14 | 4.85 | 0.08 | 5.00 | 0.00 | 199.00 | 0.00 | 5.00 | 0.00 | 5.00 | 0.00 |
| Model 1 | 100 | 50 | 10 | 11.35 | 0.23 | 0.20 | 0.09 | 0.00 | 0.00 | 0.00 | 0.00 | 10.10 | 0.10 | 0.25 | 0.12 | 2.75 | 0.57 | 10.10 | 0.10 |
| Model 1 | 200 | 50 | 10 | 10.05 | 0.05 | 4.10 | 0.28 | 0.00 | 0.00 | 0.15 | 0.08 | 10.05 | 0.05 | 8.05 | 0.82 | 10.00 | 0.00 | 10.00 | 0.00 |
| Model 1 | 100 | 100 | 10 | 99.00 | 0.00 | 99.00 | 0.00 | 4.95 | 4.95 | 9.90 | 6.81 | 19.60 | 6.07 | 99.00 | 0.00 | 2.60 | 0.50 | 10.10 | 0.07 |
| Model 1 | 200 | 100 | 10 | 10.35 | 0.13 | 5.00 | 0.26 | 0.00 | 0.00 | 0.00 | 0.00 | 10.00 | 0.00 | 9.50 | 0.50 | 9.95 | 0.05 | 10.00 | 0.00 |
| Model 1 | 100 | 200 | 10 | 99.00 | 0.00 | 0.70 | 0.18 | 0.00 | 0.00 | 0.00 | 0.00 | 10.30 | 0.16 | 99.00 | 0.00 | 10.00 | 0.00 | 10.00 | 0.00 |
| Model 1 | 200 | 200 | 10 | 189.05 | 9.95 | 189.05 | 9.95 | 9.95 | 9.95 | 19.90 | 13.70 | 19.00 | 9.49 | 189.05 | 9.95 | 9.90 | 0.10 | 10.00 | 0.00 |
| Model 1 | 100 | 500 | 10 | 99.00 | 0.00 | 1.95 | 0.18 | 0.00 | 0.00 | 0.00 | 0.00 | 10.00 | 0.00 | 99.00 | 0.00 | 10.00 | 0.00 | 10.00 | 0.00 |
| Model 1 | 200 | 500 | 10 | 199.00 | 0.00 | 9.95 | 0.05 | 0.00 | 0.00 | 0.00 | 0.00 | 10.00 | 0.00 | 199.00 | 0.00 | 10.00 | 0.00 | 10.00 | 0.00 |
| Model 2 | 100 | 50 | 2 | 2.10 | 0.07 | 2.00 | 0.00 | 2.00 | 0.00 | 2.00 | 0.00 | 2.00 | 0.00 | 1.85 | 0.08 | 2.00 | 0.00 | 2.00 | 0.00 |
| Model 2 | 200 | 50 | 2 | 2.25 | 0.12 | 2.00 | 0.00 | 2.00 | 0.00 | 2.00 | 0.00 | 2.25 | 0.12 | 1.90 | 0.07 | 2.00 | 0.00 | 2.00 | 0.00 |
| Model 2 | 100 | 100 | 2 | 99.00 | 0.00 | 99.00 | 0.00 | 16.55 | 7.95 | 16.55 | 7.95 | 16.65 | 7.94 | 99.00 | 0.00 | 2.00 | 0.00 | 2.00 | 0.00 |
| Model 2 | 200 | 100 | 2 | 2.60 | 0.29 | 2.00 | 0.00 | 2.00 | 0.00 | 2.00 | 0.00 | 2.30 | 0.18 | 1.85 | 0.08 | 2.00 | 0.00 | 2.00 | 0.00 |
| Model 2 | 100 | 200 | 2 | 99.00 | 0.00 | 2.00 | 0.00 | 2.00 | 0.00 | 2.00 | 0.00 | 2.30 | 0.18 | 99.00 | 0.00 | 2.00 | 0.00 | 2.00 | 0.00 |
| Model 2 | 200 | 200 | 2 | 199.00 | 0.00 | 199.00 | 0.00 | 2.00 | 0.00 | 2.00 | 0.00 | 2.10 | 0.10 | 199.00 | 0.00 | 2.00 | 0.00 | 2.00 | 0.00 |
| Model 2 | 100 | 500 | 2 | 99.00 | 0.00 | 2.00 | 0.00 | 2.00 | 0.00 | 2.00 | 0.00 | 2.05 | 0.05 | 99.00 | 0.00 | 1.90 | 0.07 | 1.95 | 0.05 |
| Model 2 | 200 | 500 | 2 | 199.00 | 0.00 | 2.00 | 0.00 | 2.00 | 0.00 | 2.00 | 0.00 | 2.10 | 0.10 | 199.00 | 0.00 | 2.00 | 0.00 | 2.00 | 0.00 |
| Model 2 | 100 | 50 | 5 | 5.70 | 0.28 | 3.70 | 0.21 | 0.05 | 0.05 | 0.15 | 0.11 | 5.10 | 0.10 | 4.75 | 0.25 | 5.00 | 0.00 | 5.00 | 0.00 |
| Model 2 | 200 | 50 | 5 | 5.35 | 0.17 | 5.00 | 0.00 | 3.75 | 0.18 | 4.20 | 0.16 | 5.15 | 0.08 | 4.25 | 0.35 | 5.00 | 0.00 | 5.00 | 0.00 |
| Model 2 | 100 | 100 | 5 | 99.00 | 0.00 | 99.00 | 0.00 | 0.00 | 0.00 | 14.85 | 8.11 | 19.10 | 7.70 | 99.00 | 0.00 | 4.80 | 0.12 | 5.00 | 0.00 |
| Model 2 | 200 | 100 | 5 | 5.00 | 0.30 | 4.75 | 0.25 | 1.95 | 0.20 | 3.45 | 0.22 | 4.90 | 0.28 | 4.00 | 0.40 | 5.00 | 0.00 | 5.00 | 0.00 |
| Model 2 | 100 | 200 | 5 | 99.00 | 0.00 | 5.00 | 0.00 | 0.00 | 0.00 | 0.00 | 0.00 | 5.05 | 0.05 | 99.00 | 0.00 | 4.90 | 0.07 | 5.00 | 0.00 |
| Model 2 | 200 | 200 | 5 | 199.00 | 0.00 | 199.00 | 0.00 | 20.25 | 13.67 | 41.35 | 18.08 | 24.40 | 13.35 | 199.00 | 0.00 | 5.00 | 0.00 | 5.00 | 0.00 |
| Model 2 | 100 | 500 | 5 | 99.00 | 0.00 | 5.00 | 0.00 | 0.00 | 0.00 | 0.05 | 0.05 | 5.00 | 0.00 | 99.00 | 0.00 | 4.95 | 0.05 | 5.00 | 0.00 |
| Model 2 | 200 | 500 | 5 | 199.00 | 0.00 | 5.00 | 0.00 | 1.90 | 0.19 | 4.85 | 0.08 | 5.00 | 0.00 | 199.00 | 0.00 | 5.00 | 0.00 | 5.00 | 0.00 |
| Model 2 | 100 | 50 | 10 | 12.80 | 0.78 | 1.25 | 0.18 | 0.00 | 0.00 | 0.00 | 0.00 | 10.00 | 0.55 | 1.00 | 0.25 | 7.35 | 0.26 | 9.25 | 0.14 |
| Model 2 | 200 | 50 | 10 | 10.40 | 0.13 | 6.30 | 0.24 | 0.45 | 0.15 | 1.00 | 0.19 | 10.15 | 0.08 | 7.75 | 0.87 | 10.00 | 0.00 | 10.00 | 0.00 |
| Model 2 | 100 | 100 | 10 | 99.00 | 0.00 | 99.00 | 0.00 | 9.90 | 6.81 | 14.85 | 8.11 | 24.05 | 7.23 | 99.00 | 0.00 | 6.20 | 0.26 | 9.85 | 0.08 |
| Model 2 | 200 | 100 | 10 | 11.50 | 0.45 | 6.00 | 0.22 | 0.00 | 0.00 | 0.15 | 0.08 | 10.30 | 0.15 | 7.35 | 0.93 | 9.95 | 0.05 | 10.00 | 0.00 |
| Model 2 | 100 | 200 | 10 | 99.00 | 0.00 | 1.05 | 0.18 | 0.00 | 0.00 | 0.00 | 0.00 | 10.35 | 0.18 | 99.00 | 0.00 | 9.45 | 0.15 | 9.65 | 0.15 |
| Model 2 | 200 | 200 | 10 | 199.00 | 0.00 | 199.00 | 0.00 | 0.00 | 0.00 | 9.95 | 9.95 | 10.00 | 0.00 | 199.00 | 0.00 | 9.70 | 0.13 | 10.00 | 0.00 |
| Model 2 | 100 | 500 | 10 | 99.00 | 0.00 | 2.50 | 0.17 | 0.00 | 0.00 | 0.00 | 0.00 | 10.10 | 0.10 | 99.00 | 0.00 | 9.50 | 0.26 | 9.65 | 0.11 |
| Model 2 | 200 | 500 | 10 | 199.00 | 0.00 | 9.75 | 0.10 | 0.00 | 0.00 | 0.00 | 0.00 | 10.00 | 0.00 | 199.00 | 0.00 | 10.00 | 0.00 | 10.00 | 0.00 |

**Table 7S.** Scenario 2: Estimated rank for information criterion-based ANN and RDA-based CV with low noise; signal-to-noise ratio (s2n) = 10. For all combinations of parameters $(n, p, q, k)$. Mean values are presented with standard error.

| | | | | ANN | | | | | | | | | | | | RDA | | | |
|---|---|---|---|---|---|---|---|---|---|---|---|---|---|---|---|---|---|---|---|
| | | | | AIC | | BIC | | GIC | | BICP | | GCV | | StARS | | c-rda.cv | | rrda.cv | |
| Model | n | pq | k | mean | se | mean | se | mean | se | mean | se | mean | se | mean | se | mean | se | mean | se |
| Model 1 | 100 | 50 | 2 | 2.25 | 0.12 | 2.00 | 0.00 | 2.00 | 0.00 | 2.00 | 0.00 | 2.25 | 0.12 | 1.30 | 0.19 | 2.00 | 0.00 | 2.30 | 0.13 |
| Model 1 | 200 | 50 | 2 | 2.40 | 0.13 | 2.00 | 0.00 | 2.00 | 0.00 | 2.00 | 0.00 | 2.25 | 0.10 | 1.70 | 0.13 | 2.00 | 0.00 | 2.05 | 0.05 |
| Model 1 | 100 | 100 | 2 | 98.85 | 0.08 | 98.85 | 0.08 | 2.00 | 0.00 | 2.00 | 0.00 | 2.25 | 0.10 | 98.85 | 0.08 | 2.00 | 0.00 | 2.00 | 0.00 |
| Model 1 | 200 | 100 | 2 | 2.00 | 0.00 | 2.00 | 0.00 | 2.00 | 0.00 | 2.00 | 0.00 | 2.00 | 0.00 | 1.50 | 0.14 | 2.00 | 0.00 | 2.00 | 0.00 |
| Model 1 | 100 | 200 | 2 | 99.00 | 0.00 | 2.00 | 0.00 | 2.00 | 0.00 | 2.00 | 0.00 | 2.25 | 0.16 | 99.00 | 0.00 | 2.00 | 0.00 | 2.00 | 0.00 |
| Model 1 | 200 | 200 | 2 | 198.65 | 0.11 | 198.65 | 0.11 | 2.00 | 0.00 | 2.00 | 0.00 | 2.25 | 0.16 | 198.65 | 0.11 | 2.00 | 0.00 | 2.15 | 0.11 |
| Model 1 | 100 | 500 | 2 | 99.00 | 0.00 | 2.00 | 0.00 | 2.00 | 0.00 | 2.00 | 0.00 | 2.20 | 0.12 | 99.00 | 0.00 | 2.00 | 0.00 | 2.00 | 0.00 |
| Model 1 | 200 | 500 | 2 | 199.00 | 0.00 | 2.00 | 0.00 | 2.00 | 0.00 | 2.00 | 0.00 | 2.25 | 0.16 | 199.00 | 0.00 | 2.00 | 0.00 | 2.00 | 0.00 |
| Model 1 | 100 | 50 | 5 | 5.25 | 0.18 | 5.00 | 0.00 | 5.00 | 0.00 | 5.00 | 0.00 | 5.00 | 0.00 | 3.25 | 0.50 | 5.00 | 0.00 | 5.05 | 0.05 |
| Model 1 | 200 | 50 | 5 | 5.10 | 0.07 | 5.00 | 0.00 | 5.00 | 0.00 | 5.00 | 0.00 | 5.10 | 0.07 | 4.45 | 0.30 | 5.00 | 0.00 | 5.05 | 0.05 |
| Model 1 | 100 | 100 | 5 | 99.00 | 0.00 | 99.00 | 0.00 | 19.10 | 7.70 | 23.80 | 8.63 | 19.10 | 7.70 | 99.00 | 0.00 | 5.00 | 0.00 | 5.05 | 0.05 |
| Model 1 | 200 | 100 | 5 | 5.20 | 0.16 | 5.00 | 0.00 | 5.00 | 0.00 | 5.00 | 0.00 | 5.15 | 0.15 | 3.80 | 0.42 | 5.00 | 0.00 | 5.00 | 0.00 |
| Model 1 | 100 | 200 | 5 | 99.00 | 0.00 | 5.00 | 0.00 | 5.00 | 0.00 | 5.00 | 0.00 | 5.10 | 0.10 | 99.00 | 0.00 | 5.00 | 0.00 | 5.05 | 0.05 |
| Model 1 | 200 | 200 | 5 | 198.90 | 0.07 | 198.90 | 0.07 | 5.00 | 0.00 | 24.40 | 13.35 | 14.70 | 9.70 | 198.90 | 0.07 | 5.00 | 0.00 | 5.00 | 0.00 |
| Model 1 | 100 | 500 | 5 | 99.00 | 0.00 | 5.00 | 0.00 | 5.00 | 0.00 | 5.00 | 0.00 | 5.10 | 0.10 | 99.00 | 0.00 | 5.00 | 0.00 | 5.00 | 0.00 |
| Model 1 | 200 | 500 | 5 | 199.00 | 0.00 | 5.00 | 0.00 | 5.00 | 0.00 | 5.00 | 0.00 | 5.30 | 0.18 | 199.00 | 0.00 | 5.00 | 0.00 | 5.05 | 0.05 |
| Model 1 | 100 | 50 | 10 | 10.25 | 0.18 | 10.00 | 0.00 | 0.05 | 0.05 | 0.20 | 0.16 | 10.00 | 0.00 | 8.60 | 0.77 | 10.00 | 0.00 | 10.00 | 0.00 |
| Model 1 | 200 | 50 | 10 | 10.05 | 0.05 | 10.00 | 0.00 | 10.00 | 0.00 | 10.00 | 0.00 | 10.00 | 0.00 | 9.15 | 0.59 | 10.00 | 0.00 | 10.05 | 0.05 |
| Model 1 | 100 | 100 | 10 | 99.00 | 0.00 | 99.00 | 0.00 | 14.85 | 8.11 | 34.65 | 10.83 | 18.90 | 6.13 | 99.00 | 0.00 | 10.00 | 0.00 | 10.05 | 0.05 |
| Model 1 | 200 | 100 | 10 | 10.75 | 0.38 | 10.00 | 0.00 | 10.00 | 0.00 | 10.00 | 0.00 | 10.00 | 0.00 | 8.20 | 0.83 | 10.00 | 0.00 | 10.05 | 0.05 |
| Model 1 | 100 | 200 | 10 | 99.00 | 0.00 | 10.00 | 0.00 | 0.00 | 0.00 | 0.05 | 0.05 | 10.00 | 0.00 | 99.00 | 0.00 | 10.00 | 0.00 | 10.05 | 0.05 |
| Model 1 | 200 | 200 | 10 | 199.00 | 0.00 | 199.00 | 0.00 | 19.45 | 9.45 | 47.80 | 17.34 | 19.55 | 9.45 | 199.00 | 0.00 | 10.00 | 0.00 | 10.00 | 0.00 |
| Model 1 | 100 | 500 | 10 | 99.00 | 0.00 | 10.00 | 0.00 | 0.00 | 0.00 | 0.00 | 0.00 | 10.00 | 0.00 | 99.00 | 0.00 | 10.00 | 0.00 | 10.00 | 0.00 |
| Model 1 | 200 | 500 | 10 | 199.00 | 0.00 | 10.00 | 0.00 | 10.00 | 0.00 | 10.00 | 0.00 | 10.20 | 0.20 | 199.00 | 0.00 | 10.00 | 0.00 | 10.00 | 0.00 |
| Model 2 | 100 | 50 | 2 | 2.75 | 0.29 | 2.00 | 0.00 | 2.00 | 0.00 | 2.00 | 0.00 | 2.25 | 0.12 | 1.50 | 0.17 | 2.00 | 0.00 | 2.00 | 0.00 |
| Model 2 | 200 | 50 | 2 | 2.30 | 0.21 | 2.00 | 0.00 | 2.00 | 0.00 | 2.00 | 0.00 | 2.20 | 0.12 | 1.45 | 0.14 | 2.00 | 0.00 | 2.00 | 0.00 |
| Model 2 | 100 | 100 | 2 | 98.85 | 0.08 | 98.85 | 0.08 | 6.85 | 4.85 | 16.55 | 7.95 | 16.75 | 7.93 | 98.85 | 0.08 | 2.00 | 0.00 | 2.00 | 0.00 |
| Model 2 | 200 | 100 | 2 | 2.25 | 0.14 | 2.00 | 0.00 | 2.00 | 0.00 | 2.00 | 0.00 | 2.25 | 0.14 | 1.55 | 0.11 | 2.00 | 0.00 | 2.00 | 0.00 |
| Model 2 | 100 | 200 | 2 | 99.00 | 0.00 | 2.00 | 0.00 | 2.00 | 0.00 | 2.00 | 0.00 | 2.40 | 0.15 | 99.00 | 0.00 | 2.00 | 0.00 | 2.00 | 0.00 |
| Model 2 | 200 | 200 | 2 | 188.50 | 9.92 | 188.50 | 9.92 | 1.90 | 0.10 | 1.90 | 0.10 | 2.05 | 0.18 | 188.50 | 9.92 | 2.00 | 0.00 | 2.00 | 0.00 |
| Model 2 | 100 | 500 | 2 | 99.00 | 0.00 | 2.00 | 0.00 | 2.00 | 0.00 | 2.00 | 0.00 | 2.40 | 0.28 | 99.00 | 0.00 | 2.00 | 0.00 | 2.00 | 0.00 |
| Model 2 | 200 | 500 | 2 | 199.00 | 0.00 | 2.00 | 0.00 | 2.00 | 0.00 | 2.00 | 0.00 | 2.20 | 0.16 | 199.00 | 0.00 | 2.00 | 0.00 | 2.00 | 0.00 |
| Model 2 | 100 | 50 | 5 | 6.00 | 0.25 | 5.00 | 0.00 | 5.00 | 0.00 | 5.00 | 0.00 | 5.40 | 0.13 | 3.25 | 0.50 | 5.00 | 0.00 | 5.00 | 0.00 |
| Model 2 | 200 | 50 | 5 | 5.55 | 0.23 | 5.00 | 0.00 | 5.00 | 0.00 | 5.00 | 0.00 | 5.10 | 0.07 | 3.35 | 0.42 | 5.00 | 0.00 | 5.00 | 0.00 |
| Model 2 | 100 | 100 | 5 | 98.90 | 0.07 | 98.90 | 0.07 | 9.70 | 4.70 | 9.70 | 4.70 | 10.00 | 4.69 | 98.90 | 0.07 | 5.00 | 0.00 | 5.00 | 0.00 |
| Model 2 | 200 | 100 | 5 | 5.55 | 0.26 | 5.00 | 0.00 | 5.00 | 0.00 | 5.00 | 0.00 | 5.15 | 0.11 | 2.50 | 0.44 | 5.00 | 0.00 | 5.00 | 0.00 |
| Model 2 | 100 | 200 | 5 | 99.00 | 0.00 | 5.00 | 0.00 | 5.00 | 0.00 | 5.00 | 0.00 | 5.40 | 0.20 | 99.00 | 0.00 | 5.00 | 0.00 | 5.00 | 0.00 |
| Model 2 | 200 | 200 | 5 | 198.55 | 0.11 | 198.55 | 0.11 | 5.00 | 0.00 | 24.40 | 13.35 | 14.90 | 9.69 | 198.55 | 0.11 | 5.00 | 0.00 | 5.00 | 0.00 |
| Model 2 | 100 | 500 | 5 | 99.00 | 0.00 | 5.00 | 0.00 | 5.00 | 0.00 | 5.00 | 0.00 | 5.15 | 0.15 | 99.00 | 0.00 | 4.75 | 0.14 | 4.90 | 0.10 |
| Model 2 | 200 | 500 | 5 | 199.00 | 0.00 | 5.00 | 0.00 | 5.00 | 0.00 | 5.00 | 0.00 | 5.00 | 0.00 | 199.00 | 0.00 | 5.00 | 0.00 | 5.00 | 0.00 |
| Model 2 | 100 | 50 | 10 | 13.05 | 0.54 | 10.00 | 0.00 | 0.40 | 0.15 | 1.05 | 0.41 | 10.10 | 0.07 | 3.95 | 1.02 | 10.00 | 0.00 | 10.00 | 0.00 |
| Model 2 | 200 | 50 | 10 | 10.25 | 0.14 | 10.00 | 0.00 | 10.00 | 0.00 | 10.00 | 0.00 | 10.05 | 0.05 | 5.75 | 0.98 | 10.00 | 0.00 | 10.00 | 0.00 |
| Model 2 | 100 | 100 | 10 | 98.90 | 0.07 | 98.90 | 0.07 | 19.80 | 9.08 | 29.75 | 10.40 | 28.15 | 8.13 | 98.90 | 0.07 | 10.00 | 0.00 | 10.00 | 0.00 |
| Model 2 | 200 | 100 | 10 | 12.10 | 0.55 | 10.00 | 0.00 | 10.00 | 0.00 | 10.00 | 0.00 | 10.25 | 0.16 | 4.40 | 0.96 | 10.00 | 0.00 | 10.00 | 0.00 |
| Model 2 | 100 | 200 | 10 | 99.00 | 0.00 | 10.00 | 0.00 | 0.00 | 0.00 | 0.10 | 0.07 | 10.00 | 0.00 | 99.00 | 0.00 | 10.00 | 0.00 | 10.00 | 0.00 |
| Model 2 | 200 | 200 | 10 | 198.50 | 0.14 | 198.50 | 0.14 | 10.00 | 0.00 | 10.00 | 0.00 | 10.00 | 0.00 | 198.50 | 0.14 | 10.00 | 0.00 | 10.00 | 0.00 |
| Model 2 | 100 | 500 | 10 | 99.00 | 0.00 | 10.00 | 0.00 | 0.00 | 0.00 | 0.25 | 0.10 | 10.00 | 0.00 | 99.00 | 0.00 | 9.95 | 0.05 | 9.95 | 0.05 |
| Model 2 | 200 | 500 | 10 | 199.00 | 0.00 | 10.00 | 0.00 | 10.00 | 0.00 | 10.00 | 0.00 | 10.00 | 0.00 | 199.00 | 0.00 | 10.00 | 0.00 | 10.00 | 0.00 |

**Table 8S.** Scenario 3: The `rrda` prediction performance for simulation data. Signal to noise = 0.1,1,10. Model 1 (The Latent Space Model) and Model 2 (the Matrix Factorization Model) were tested. The estimations of rank, MSPE, $\lambda$, and Computation time of parameter tuning for random-splitted samples (100 iterations), and tested on the test data by MSPE 100 times. Comparison of model fit with various tuning parameters. CV (5-fold) is performed for the `rrda` model. The computation times given here are measured using sequential executions without parallel computing. Mean values are presented with standard error in the parentheses. For details on the session information, see Table 1S.

| | | | **ANN** | | | | | | **RDA** | |
|---|---|---|---|---|---|---|---|---|---|---|
| **s2n** | **Model** | **Method** | **AIC** | **BIC** | **GIC** | **BICP** | **GCV** | **StARS** | **c-rda.cv** | **rrda.cv** |
| 0.1 | Model 1 | Rank | 89.00 (0.00) | 0.00 (0.00) | 0.00 (0.00) | 0.00 (0.00) | 7.12 (0.09) | 89.00 (0.00) | 1.00 (0.00) | 5.40 (0.10) |
| | | $\lambda$ | - - | - - | - - | - - | - - | - - | - - | 1.90e+04 (209.60) |
| | | MSPE | 8.98e+01 (3.34e-1) | 5.00e+01 (1.54e-1) | 5.00e+01 (1.54e-1) | 5.00e+01 (1.54e-1) | 7.60e+07 (2.31e+07) | 8.98e+01 (3.34e-1) | 2.13e+04 (1.66e+04) | **4.86e+01** (1.44e-1) |
| | | Time [s] | 0.488 (0.004) | 0.484 (0.005) | 0.497 (0.005) | 0.494 (0.005) | 0.486 (0.005) | 2.04 (0.009) | 0.57 (0.005) | 4.54 (0.006) |
| | Model 2 | Rank | 89.00 (0.00) | 0.00 (0.00) | 0.00 (0.00) | 0.00 (0.00) | 5.63 (0.07) | 89.00 (0.00) | 1.00 (0.00) | 3.03 (0.17) |
| | | $\lambda$ | - - | - - | - - | - - | - - | - - | - - | 191 (9.24) |
| | | MSPE | 1.97e+01 (9.10e-2) | 1.09e+01 (3.69e-2) | 1.09e+01 (3.69e-2) | 1.09e+01 (3.69e-2) | 1.04e+07 (3.70e+06) | 1.97e+01 (9.10e-2) | 2.65e+03 (5.58e+02) | **1.08e+01** (3.65e-2) |
| | | Time [s] | 0.522 (0.006) | 0.515 (0.007) | 0.511 (0.006) | 0.513 (0.006) | 0.520 (0.007) | 2.38 (0.014) | 0.58 (0.006) | 3.85 (0.010) |
| 1 | Model 1 | Rank | 89.00 (0.00) | 4.58 (0.05) | 0.00 (0.00) | 0.00 (0.00) | 5.00 (0.00) | 89.00 (0.00) | 5.00 (0.00) | 5.00 (0.00) |
| | | $\lambda$ | - - | - - | - - | - - | - - | - - | - - | 888 (9.39) |
| | | MSPE | 8.93 (3.41e-2) | 5.47 (3.96e-2) | 9.06 (7.78e-2) | 9.06 (7.78e-2) | 5.18 (1.72e-2) | 8.93 (3.41e-2) | 5.18 (1.72e-2) | **5.00** (1.57e-2) |
| | | Time [s] | 0.39 (0.003) | 0.389 (0.003) | 0.387 (0.003) | 0.386 (0.003) | 0.389 (0.003) | 1.89 (0.007) | 0.57 (0.005) | 4.38 (0.007) |
| | Model 2 | Rank | 89.00 (0.00) | 4.09 (0.03) | 0.00 (0.00) | 0.11 (0.03) | 5.00 (0.00) | 89.00 (0.00) | 4.78 (0.04) | 4.94 (0.02) |
| | | $\lambda$ | - - | - - | - - | - - | - - | - - | - - | 28.20 (1.00) |
| | | MSPE | 2.62 (1.64e-2) | 1.69 (9.90e-3) | 1.97 (1.59e-2) | 1.97 (1.58e-2) | 6.13 (4.44) | 2.62 (1.64e-2) | 6.13 (4.44) | **1.62** (9.30e-3) |
| | | Time [s] | 0.424 (0.005) | 0.414 (0.005) | 0.405 (0.002) | 0.414 (0.004) | 0.417 (0.005) | 2.21 (0.013) | 0.57 (0.004) | 3.77 (0.011) |
| 10 | Model 1 | Rank | 89.00 (0.00) | 5.00 (0.00) | 5.00 (0.00) | 5.00 (0.00) | 5.06 (0.03) | 89.00 (0.00) | 5.00 (0.00) | 5.03 (0.02) |
| | | $\lambda$ | - - | - - | - - | - - | - - | - - | - - | 152.00 (1.54) |
| | | MSPE | 8.94e-1 (3.50e-3) | 5.18e-1 (1.80e-3) | 5.18e-1 (1.80e-3) | 5.18e-1 (1.80e-3) | 1.20e+01 (7.83) | 8.94e-1 (3.50e-3) | 5.18e-1 (1.80e-3) | **5.01e-1** (1.60e-3) |
| | | Time [s] | 0.347 (0.002) | 0.348 (0.004) | 0.341 (0.002) | 0.342 (0.003) | 0.344 (0.003) | 1.89 (0.008) | 0.57 (0.004) | 4.51 (0.007) |
| | Model 2 | Rank | 89.00 (0.00) | 5.00 (0.00) | 5.00 (0.00) | 5.00 (0.00) | 5.00 (0.00) | 89.00 (0.00) | 4.89 (0.03) | 4.90 (0.02) |
| | | $\lambda$ | - - | - - | - - | - - | - - | - - | - - | 15.70 (0.78) |
| | | MSPE | 1.03 (1.82e-2) | 7.30e-1 (9.40e-3) | 7.30e-1 (9.40e-3) | 7.30e-1 (9.40e-3) | 7.30e-1 (9.40e-3) | 1.03 (1.82e-2) | 7.30e-1 (9.50e-3) | **7.03e-1** (8.80e-3) |
| | | Time [s] | 0.379 (0.003) | 0.376 (0.005) | 0.374 (0.004) | 0.376 (0.006) | 0.373 (0.005) | 2.19 (0.012) | 0.57 (0.004) | 3.82 (0.012) |

**Table 9S.** Scenario 3: Parameter selections for Model 1 (Latent Space Model) and Model 2 (the Matrix Factorization Model). Random-Splitting (100 iterations) for the following rank and $\lambda$ parameters. We compared the following six parameter settings: (1) **CV min**, which uses the best set of parameters selected by CV-i.e., the conventional CV approach; (2) **Rank.1se**, which selects the rank using the one-standard-error rule while keeping $\lambda$ the same as in CV min; (3) **$\boldsymbol{\lambda}$.1se**, which selects $\boldsymbol{\lambda}$ using the one-standard-error rule while keeping the rank the same as in CV min; (4) **Full Rank**, a full-rank multivariate model using the $\boldsymbol{\lambda}$ from CV min; (5) **$\boldsymbol{\lambda}$0**, which uses the rank estimated by CV min without applying $\boldsymbol{\lambda}$ regularization; and (6) **Full Rank + $\boldsymbol{\lambda}$0**, a standard multivariate linear model without rank restriction or regularization. Mean values are presented with standard error in the parentheses. Parameter selections. Signal to noise = 0.1,1,10. For details on the session information, see Table 1S.

| s2n | Model | Method | CV min | Rank.1se | $\lambda$.1se | Full Rank | $\lambda$0 | Full Rank+$\lambda$0 |
|---|---|---|---|---|---|---|---|---|
| 0.1 | Model 1 | Rank | 5.40<br>(0.10) | 2.91<br>(0.09) | 5.40<br>(0.10) | 90.00<br>(0.00) | 5.40<br>(0.10) | 90.00<br>(0.00) |
| | | $\lambda$ | 1.90e+04<br>(2.10e+02) | 1.90e+04<br>(2.10e+02) | 8.09e+04<br>(2.30e+03) | 1.90e+04<br>(2.10e+02) | 0.00<br>(0.00) | 0.00<br>(0.00) |
| | | MSPE | **4.87e+01**<br>(1.44e-1) | 4.92e+01<br>(1.48e-1) | 4.91e+01<br>(1.47e-1) | 5.07e+01<br>(1.50e-1) | 2.02e+07<br>(1.89e+07) | 8.83e+01<br>(3.05e-1) |
| | Model 2 | Rank | 3.03<br>(0.17) | 1.00<br>(0.00) | 3.03<br>(0.17) | 90.00<br>(0.00) | 3.03<br>(0.17) | 90.00<br>(0.00) |
| | | $\lambda$ | 190.89<br>(9.24) | 190.89<br>(9.24) | 5237.59<br>(222.81) | 190.89<br>(9.24) | 0.00<br>(0.00) | 0.00<br>(0.00) |
| | | MSPE | **1.08e+01**<br>(3.56e-2) | 1.08e+01<br>(3.65e-2) | 1.08e+01<br>(3.68e-2) | 1.19e+01<br>(8.19e-2) | 3.73e+05<br>(2.08e+05) | 1.91e+01<br>(8.42e-2) |
| 1 | Model 1 | Rank | 5.00<br>(0.00) | 5.00<br>(0.00) | 5.00<br>(0.00) | 90.00<br>(0.00) | 5.00<br>(0.00) | 90.00<br>(0.00) |
| | | $\lambda$ | 887.8<br>(7.61) | 887.8<br>(7.61) | 2246.88<br>(36.35) | 887.8<br>(7.61) | 0.00<br>(0.00) | 0.00<br>(0.00) |
| | | MSPE | **5.00**<br>(1.57e-2) | **5.00**<br>(1.57e-2) | 5.03<br>(1.63e-2) | 5.52<br>(1.82e-2) | 5.18<br>(1.72e-2) | 8.90<br>(3.34e-2) |
| | Model 2 | Rank | 4.94<br>(0.02) | 3.98<br>(0.06) | 4.94<br>(0.02) | 90.00<br>(0.00) | 4.94<br>(0.02) | 90.00<br>(0.00) |
| | | $\lambda$ | 28.25<br>(1.00) | 28.25<br>(1.00) | 126.02<br>(4.05) | 28.25<br>(1.00) | 0.00<br>(0.00) | 0.00<br>(0.00) |
| | | MSPE | **1.62**<br>(9.30e-3) | 1.66<br>(1.06e-2) | 1.66<br>(1.00e-2) | 2.03<br>(1.39e-2) | 6.13<br>(4.44) | 2.46<br>(1.27e-2) |
| 10 | Model 1 | Rank | 5.03<br>(0.02) | 5.00<br>(0.00) | 5.03<br>(0.02) | 90.00<br>(0.00) | 5.03<br>(0.02) | 90.00<br>(0.00) |
| | | $\lambda$ | 152.16<br>(1.54) | 152.16<br>(1.54) | 465.68<br>(7.82) | 152.16<br>(1.54) | 0.00<br>(0.00) | 0.00<br>(0.00) |
| | | MSPE | **5.01e-1**<br>(1.60e-3) | **5.01e-1**<br>(1.60e-3) | 5.04e-1<br>(1.70e-3) | 5.30e-1<br>(1.80e-3) | 1.82e+02<br>(1.82e+02) | 8.92e-1<br>(3.40e-3) |
| | Model 2 | Rank | 4.90<br>(0.03) | 4.06<br>(0.05) | 4.90<br>(0.03) | 90.00<br>(0.00) | 4.90<br>(0.03) | 90.00<br>(0.00) |
| | | $\lambda$ | 15.73<br>(0.78) | 15.73<br>(0.78) | 89.56<br>(2.94) | 15.73<br>(0.78) | 0.00<br>(0.00) | 0.00<br>(0.00) |
| | | MSPE | **7.03e-1**<br>(9.00e-3) | 7.29e-1<br>(9.90e-3) | 7.40e-1<br>(9.60e-3) | 7.50e-1<br>(9.20e-3) | 7.34e-1<br>(9.60e-3) | 8.09e-1<br>(9.30e-3) |

**Table 10S.** The `rrda` evaluation in Breast Cancer data in comparison with all criteria-based ANN and `StARS`. The estimations of rank, MSPE, $\lambda$, and Computation time of parameter tuning for random-splitted samples (100 iterations), and tested on the test data by MSPE 100 times. Comparison of model fit with various tuning parameters. CV (5-fold) is performed for the `rrda` model. The computation times given here are measured in a single core setting. Mean values are presented with standard error in the parentheses. For details on the session information, see Table 1S.

| | **ANN** | | | | | | **RDA** | |
|---|---|---|---|---|---|---|---|---|
| | **AIC** | **BIC** | **GIC** | **BICP** | **GCV** | **StARS** | **c-rda.cv** | **rrda.cv** |
| The Chromosome 13 subset | | | | | | | | |
| Rank | 56.95<br>(0.02) | 37.26<br>(2.64) | 1.19<br>(0.03) | 1.82<br>(0.03) | 24.47<br>(0.13) | 2.40<br>(0.08) | 2.05<br>(0.02) | 2.75<br>(0.05) |
| $\lambda$ | -<br>- | -<br>- | -<br>- | -<br>- | -<br>- | -<br>- | -<br>- | 52.8<br>(1.60) |
| MSPE | 4.39e-2<br>(1.00e-3) | 4.07e-2<br>(1.20e-3) | 3.68e-2<br>(1.20e-3) | 3.60e-2<br>(1.20e-3) | 4.29e-2<br>(1.00e-3) | 3.58e-2<br>(1.10e-3) | 3.46e-2<br>(1.00e-3) | **2.70e-2**<br>(9.00e-4) |
| Time [s] | 0.065<br>(0.00) | 0.062<br>(0.00) | 0.067<br>(0.00) | 0.064<br>(0.00) | 0.065<br>(0.00) | 32.34<br>(0.19) | 0.606<br>(0.00442) | 2.13<br>(0.00) |
| Whole Genome (Chromosome 1-23) | | | | | | | | |
| Rank | 78.00<br>(0.00) | 1.07<br>(0.02) | 0.00<br>(0.00) | 0.00<br>(0.00) | 67.86<br>(0.15) | 78.00<br>(0.00) | 5.94<br>(0.22) | 46.73<br>(1.62) |
| $\lambda$ | -<br>- | -<br>- | -<br>- | -<br>- | -<br>- | -<br>- | -<br>- | 3.66e+3<br>(101.94) |
| MSPE | 4.21e-2<br>(8.00e-4) | 4.64e-2<br>(2.20e-3) | 4.54e-2<br>(1.00e-3) | 4.54e-2<br>(1.00e-3) | 4.20e-2<br>(8.00e-4) | 4.21e-2<br>(8.00e-4) | 4.56e-1<br>(1.52e-1) | **4.11e-2**<br>(9.00e-4) |
| Time [s] | 24.02<br>(0.45) | 20.61<br>(0.02) | 20.59<br>(0.02) | 20.54<br>(0.02) | 63.22<br>(18.80) | 368.12<br>(0.26) | 2.71<br>(0.01) | 39.30<br>(0.04) |

**Table 11S.** Comparison of `rrda` and `sRDA` (Csala et al., 2017) in terms of MSPE and computation time on the Breast Cancer dataset ($X \in \mathbb{R}^{89\times19,672}$, $Y \in \mathbb{R}^{89\times2,149}$). The evaluation was based on five random splits (train: 79, test: 10); within each training set, a 5-fold CV was used for parameter tuning. For `rrda`, parameters were selected to minimize the mean squared error (MSE). For `sRDA`, parameters were chosen to maximize the sum of absolute correlations following the procedure described by Csala et al. (2017). We also evaluated MSE-based parameter selection for `sRDA`, which yielded similar performance. For `rrda`, ranks 1–40 and fifty values of $\lambda$, defined as $\lambda = 10^s$ with $s$ equally spaced over the interval $[0, 6]$, were tested. For `sRDA`, both UST (Univariate Soft Thresholding) and Elastic Net penalties were considered using 1–5 latent variables, with the number of nonzero weights selected from $\{5, 10, 20, 50, 100\}$. For the Elastic Net, $\lambda \in \{0.1, 1, 10, 100\}$. After observing the prediction accuracy, we confirmed the parameter range is plausible to maximize the sum of absolute correlations (Figure 4S). UST represents a limiting case of the Elastic Net where the $\ell_2$ component dominates (Waldmann et al., 2013; Zou and Hastie, 2005). MSPE values are averaged across the 5 splits, with standard errors in parentheses. For details on the session information, see Table 1S.

| | rrda | sRDA-UST | sRDA-Enet |
|---|---|---|---|
| Rank | 27.60 (0.6) | 1.00 (0.0) | 1.00 (0.0) |
| Nonzero weights | – | 50.00 (0.0) | 20.00 (0.0) |
| $\lambda$ | 2682.70 (0.0) | – | 0.10 (0.0) |
| MSPE | **3.63e-2 (0.04e-2)** | 8.30e-2 (0.27e-2) | 9.10e-2 (0.28e-2) |
| Time [s] | 20.80 (0.07) | 175.10 (0.80) | 24295.12 (455.88) |

**Table 12S.** The rrda evaluation in Soybean dataset. The estimations of rank, MSPE, $\lambda$, and Computation time for parameter tuning with random-splitting samples (100 iterations), and tested on the test data by MSPE 100 times. Comparison of model fit with various tuning parameters. CV (5-fold) is performed for the rrda model. The rank is estimated to be 0 when no variables are selected in the model. The computation times given here are measured using sequential executions without parallel computing. Mean values are presented with standard error in the parentheses. For details on the session information, see Table 1S.

| | **ANN** | | | | | | **RDA** | |
|---|---|---|---|---|---|---|---|---|
| | **AIC** | **BIC** | **GIC** | **BICP** | **GCV** | **StARS** | **c-rda.cv** | **rrda.cv** |
| Rank | 158.00<br>(0.00) | 3.00<br>(0.00) | 0.00<br>(0.00) | 0.00<br>(0.00) | 43.44<br>(0.38) | 158.00<br>(0.00) | 9.23<br>(0.13) | 11.24<br>(0.13) |
| $\lambda$ | -<br>- | -<br>- | -<br>- | -<br>- | -<br>- | -<br>- | -<br>- | 309<br>(9.80) |
| MSPE | 1.02<br>(1.62e-2) | 6.21<br>(1.20) | 1.01<br>(1.55e-2) | 1.01<br>(1.55e-2) | 0.99<br>(1.60e-2) | 1.02<br>(1.62e-2) | 5.03<br>(1.22) | **0.94**<br>(1.55e-2) |
| Time [s] | 1.32<br>(0.01) | 1.12<br>(0.01) | 1.12<br>(0.01) | 1.10<br>(0.01) | 1.15<br>(0.01) | 27.95<br>(0.03) | 2.42<br>(0.01) | 10.99<br>(0.02) |

**Table 13S.** The results of rrda.cv parallel computation with the variable information and means for CV (5-fold) with their standard errors in parentheses. For details on the session information, see Table 1S.

| | Breast Cancer (Ch13) | Breast Cancer (Whole) | Soybean | TCGA |
|---|---|---|---|---|
| $n$ | 89 | 89 | 179 | 20 |
| $p$ | 319 | 19,672 | 4771 | 392,277 |
| $q$ | 58 | 2,149 | 253 | 67,528 |
| Time [s] | 3.374 (0.037) | 5.719 (0.011) | 4.465 (0.018) | 15.60 (0.035) |
